# Reconstructing complex cancer evolutionary histories from multiple bulk DNA samples using Pairtree

**DOI:** 10.1101/2020.11.06.372219

**Authors:** Jeff A. Wintersinger, Stephanie M. Dobson, Lincoln D. Stein, John E. Dick, Quaid D. Morris

## Abstract

Cancers are composed of genetically distinct subpopulations of malignant cells. By sequencing DNA from cancer tissue samples, we can characterize the somatic mutations specific to each population and build *clone trees* describing the evolutionary relationships between these populations. These trees reveal critical points in disease development and inform treatment. Both liquid cancers and solid tumours can be profiled with this approach.

Pairtree is a new method for constructing clone trees using DNA sequencing data from one or more bulk samples of an individual cancer. It uses Bayesian inference to compute posterior distributions over the evolutionary relationships between every pair of identified subpopulations, then uses these distributions in a Markov Chain Monte Carlo algorithm to perform efficient inference of the posterior distribution over clone trees. Pairtree also uses the pairwise relationships to detect mutations that violate the infinite sites assumption. Unlike previous methods, Pairtree can perform clone tree reconstructions using as many as 100 samples per cancer that reveal 30 or more cell subpopulations. On simulated data, Pairtree is the only method whose performance reliably improves when provided with additional bulk samples from a cancer. On 14 B-progenitor acute lymphoblastic leukemias with up to 90 samples from each cancer, Pairtree was the only method that could reproduce or improve upon expert-derived clone tree reconstructions. By scaling to more challenging problems, Pairtree supports new biomedical research applications that can improve our understanding of the natural history of cancer, as well as better illustrate the interplay between cancer, host, and therapeutic interventions.

**Significance:** Clone trees describe the evolutionary history of a cancer and can provide insights into how the disease changed through time (e.g., between diagnosis and relapse). Pairtree uses DNA sequencing data from many samples of the same cancer to reconstruct a cancer’s evolutionary history with much greater detail and accuracy than previously possible.

## 1 Introduction

Individual cancers contain substantial genetic heterogeneity arising from an ongoing evolutionary process of random somatic mutation and selection [1]. Cancers typically arise from a small number of founder mutations that confer a growth advantage [2]. Over time, additional somatic mutations accrue, and their frequency and distribution are shaped by evolutionary forces such as selection and genetic drift, resulting in the emergence of multiple genetically distinct cell subpopulations [3] (Fig. 1a). A *clone tree* is the evolutionary tree delineating the cell subpopulations in a cancer, the genetic mutations specific to each, and the proportions of cells in each sample that arose from each subpopulation (Fig. 1). Within the tree, *subclones* correspond to a cell subpopulation and all its descendants.

**Figure 1:**
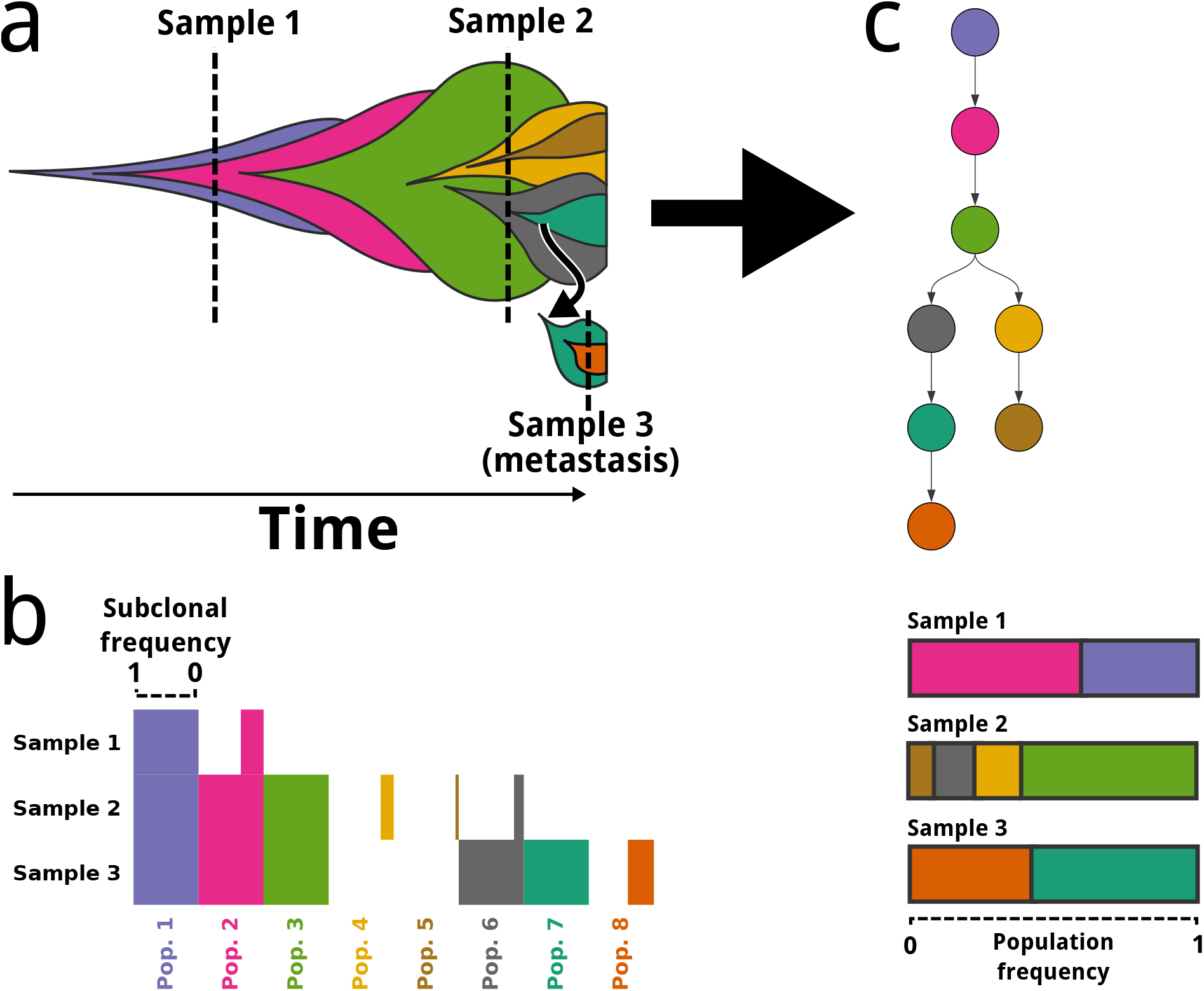
Construction of clone trees from multiple cancer samples. **a.** Schematic illustrates cancer development under the clonal evolution model. Each colour represents a genetically distinct subpopulation. Each subpopulation emerges within the mass of its parent. The leftmost point for a subpopulation denotes the cell that was its most recent common ancestor. Dashed vertical lines indicate when and where cancer samples were taken. The relative abundance of each subpopulation in a cancer sample, including any nested descendent subpopulations composing a subclone, is represented by the height of that subpopulation or subclone along the sample’s dashed line. **b.** Horizontal bar plot showing idealized input to clone tree reconstruction algorithms. Bar length indicates the subclonal frequency of each subpopulation and its descendants (column) in each sequenced sample (row). The clonal evolution model asserts that a subpopulation’s point mutations are inherited by its descendants. Consequently, mutation VAFs in DNA sequencing data provide estimates of subclonal frequencies, corresponding to the proportion of cells that originated from a subclonal population and its descendants. **c.** Clone tree representing the ancestry of subpopulations (top). Nodes indicate subpopulations. Arrows extend from each subpopulation to its direct descendants. Inferred frequencies of each subpopulation in each sample are based on the clone tree and mutation frequency data (bottom).

Clone trees built from bulk cancer samples have important biomedical applications. Those built from single samples already reveal important genomic events in evolution [3–5] and provide insights into heterogeneity [1]. But as sequencing costs continue to drop, sequencing different regions of the same tumour [6], multiple tumours of the same cancer [7], or longitudinal samples from different timepoints [8] will become more common. When bulk samples have different mixtures of subpopulations, each sample can provide unique information about the single clone tree that characterizes the cancer’s evolutionary history. This can include revealing new subpopulations or deconvolving large subpopulations into smaller constituents. Clone trees built from multiple samples of the same cancer have helped identify factors associated with metastasis [9] and probed how treatment [10–12] or tumour microenvironment [13, 14] shape evolution. This, in turn, can inform strategies to counteract treatment resistance [15]. Beyond cancer, clone trees have applications in other studies of somatic genetic heterogeneity [16, 17].

Current subclonal reconstruction methods [18–24] are severely limited in their ability to build clone trees based on large multi-sample studies. Most of these methods were designed for single cancer samples from which no more than three subclones can be discerned at typical whole-genome sequencing depths [1]. Recent studies with greater sequencing depth and multiple cancer samples have revealed that a single cancer can have dozens of resolvable subclones [6, 11]. Here we show that existing clone tree reconstruction methods become highly inaccurate on datasets with many subclones or many cancer samples, necessitating a new approach.

We introduce Pairtree, a new method that can accurately construct clone trees containing as many as 30 subclones. Pairtree outperforms a representative set of state-of-the-art clone tree reconstruction packages on simulated benchmark datasets of variable complexity. Pairtree is also the only method tested that can recover or improve upon expert reconstructions of clone trees for 14 B-progenitor acute lymphoblastic leukemias (B-ALLs) containing up to 90 samples and 26 subclones per cancer. The Pairtree method, along with an interactive visual interface for exploring the clone tree posterior, is available at https://github.com/morrislab/pairtree.

## 2 Results

### 2.1 Pairtree inputs and outputs

Fig. 1 outlines the process of constructing a clone tree to represent the evolutionary history of a cancer. Pairtree takes as input allele frequency data for point mutations detected in one or more samples from a single cancer. These data can be derived from whole-genome sequencing (WGS), whole-exome sequencing (WES), or more targeted sequencing. Each bulk cancer sample is a mixture of genetically heterogeneous cells (Fig. 1a). For each mutation, Pairtree uses counts of variant and reference reads in each sample to estimate the variant allele frequency (VAF), i.e., the proportion of reads at a mutation’s locus that contain the mutation. By correcting a mutation’s VAF for copy-number aberrations (CNAs) affecting the locus, Pairtree computes an estimate of the proportion of cells in each sample carrying the mutation, termed the mutation’s *subclonal frequency* [25] (Fig. 1b).

Pairtree outputs a set of possible clone trees explaining evolutionary relationships between the input mutations. Clone tree nodes correspond to cancerous subpopulations, while arrows (i.e., directed edges) extend from a subpopulation’s node to the nodes representing its direct descendants (Fig. 1c). We define a subpopulation as those cells containing exactly the same subset of the somatic mutations input into Pairtree. In each cancer sample, each subpopulation is assigned a population frequency, representing what proportion of cells in that sample share the same mutation subset. Note that many, if not most, of a cancer’s mutations will not be provided in the input because of incomplete genome coverage or because the mutations are too low in frequency to be detected.

Each subpopulation and its descendant subpopulations (both direct and indirect) form a subclone (Fig. 1a). Pairtree assigns a tree-constrained subclonal frequency to each subclone in each cancer sample, which is equal to the sum of the population frequencies of all the subpopulations contained within the subclone (Fig. 1a-b). This relationship follows from the infinite sites assumption (ISA), which states that no site is mutated more than once during cancer evolution. The ISA implies that subpopulations inherit all the mutations of their parent populations, and that each mutation appears only once in the evolutionary history of the cancer. Though violations of the ISA occur [26], it remains broadly valid [27], and if the input dataset includes ISA-violating mutations, Pairtree can detect and discard them before starting to build a clone tree (Section 9.6). Like most other clone tree reconstruction methods, Pairtree assumes the ISA when building trees. Other methods permit some, but not all, types of ISA-violations [28–31].

Pairtree identifies which mutations belong to each subclone based on the estimated subclonal frequencies provided by the VAF data (Fig. 1b), then searches for clone trees whose structures allow subclonal frequencies that best match these estimates (Fig. 1c). Pairtree’s output consists of a set of clone trees, each scored by a likelihood indicating how well the tree-constrained subclonal frequencies match the frequency estimates given by the VAF data. Although there is a single true clone tree explaining how subpopulations are related, this tree is not observed directly, and the input data often permit multiple solutions.

Grouping mutations into subclones is not necessary—algorithms can instead build clone trees in which each mutation is assigned to a unique subclone, yielding a mutation tree. However, because of limited resolution in the data’s estimated subclonal frequencies, sets of mutations often have subclonal frequency estimates that are too similar to separate the mutations into distinct subclones. As such, the first step in clone tree reconstruction is often clustering mutations with similar estimated subclonal frequencies across all input samples, and associating subclones with these clusters. Mutation clustering can be performed with Pairtree (Section 9.5.1) or by another method [32–34] and input into Pairtree. This step simplifies clone tree reconstruction by reducing the number of subclones. Additionally, this approach permits more precise estimates of each subclone’s subclonal frequency by combining data from the subclone’s mutations (Section 9.3.8), at the risk of grouping together mutations from different subclones. Increasing the number of cancer samples provides more subclonal frequency estimates for each mutation, thereby reducing the risk of improper mutation grouping.

### 2.2 Delineating ancestral relationships between pairs of subclones using the Pairs Tensor

Pairtree uses the estimated subclonal frequencies to predict the ancestral relationship between every subclone pair. These pairwise relationships then serve as a guide when Pairtree searches for clone trees that best fit the VAF data. Under the ISA, one of three mutually exclusive ancestral relationships exist between an ordered pair of subclones *A* and *B.* [35, 36]

#### Ancestor

*A* is ancestral to *B*. Here, the subpopulation associated with *A* contains *A*’s mutations but not *B*’s. No cell subpopulation has *B*’s mutations without also inheriting *A*’s.

#### Descendant

*B* is ancestral to *A*. As above but with the roles of *A* and *B* switched.

#### Branching

Neither *A* nor *B* is the ancestor of the other. I.e., they occur on different branches of the clone tree. Consequently, no subpopulations have both *A*’s and *B*’s mutations.

Each relationship constrains the frequencies that can be assigned to the two subclones (Section 9.2.3). For a given subclone pair, Pairtree combines a prior probability distribution incorporating these constraints with a likelihood distribution based on the VAF data for each subclone’s mutations, then uses Bayesian inference to compute the probability of each relationship type for the pair (Section 9.2). This yields a data structure termed the *Pairs Tensor*, the elements of which are the marginal posterior probability distributions over the three possible ancestral relationships for every subclone pair.

### 2.3 Using pairwise ancestry to guide the search for clone trees

Pairtree uses the Pairs Tensor to define a proposal distribution for a Markov Chain Monte Carlo (MCMC) algorithm [37] that samples from the posterior distribution over clone trees (Fig. 2). The algorithm’s Metropolis-Hastings scheme generates proposal trees using two distributions over subclones derived from the Pairs Tensor (Section 9.3.5). The first distribution helps choose a poorly placed subclone to move within the tree, with each subclone’s selection probability determined by the degree of discordance between the data-implied pairwise relationships and those imposed by its present position within the tree. The second distribution guides the choice of new parent for the selected subclone, evaluating potential destinations based on how much this discordance is decreased. Though other MCMC-based subclonal reconstruction methods also modify trees by moving subclones [18, 20, 38] or mutations [39, 40], Pairtree is the first to guide this decision with data, allowing the algorithm to rapidly navigate to and explore high-probability regions of the clone-tree posterior.

**Figure 2:**
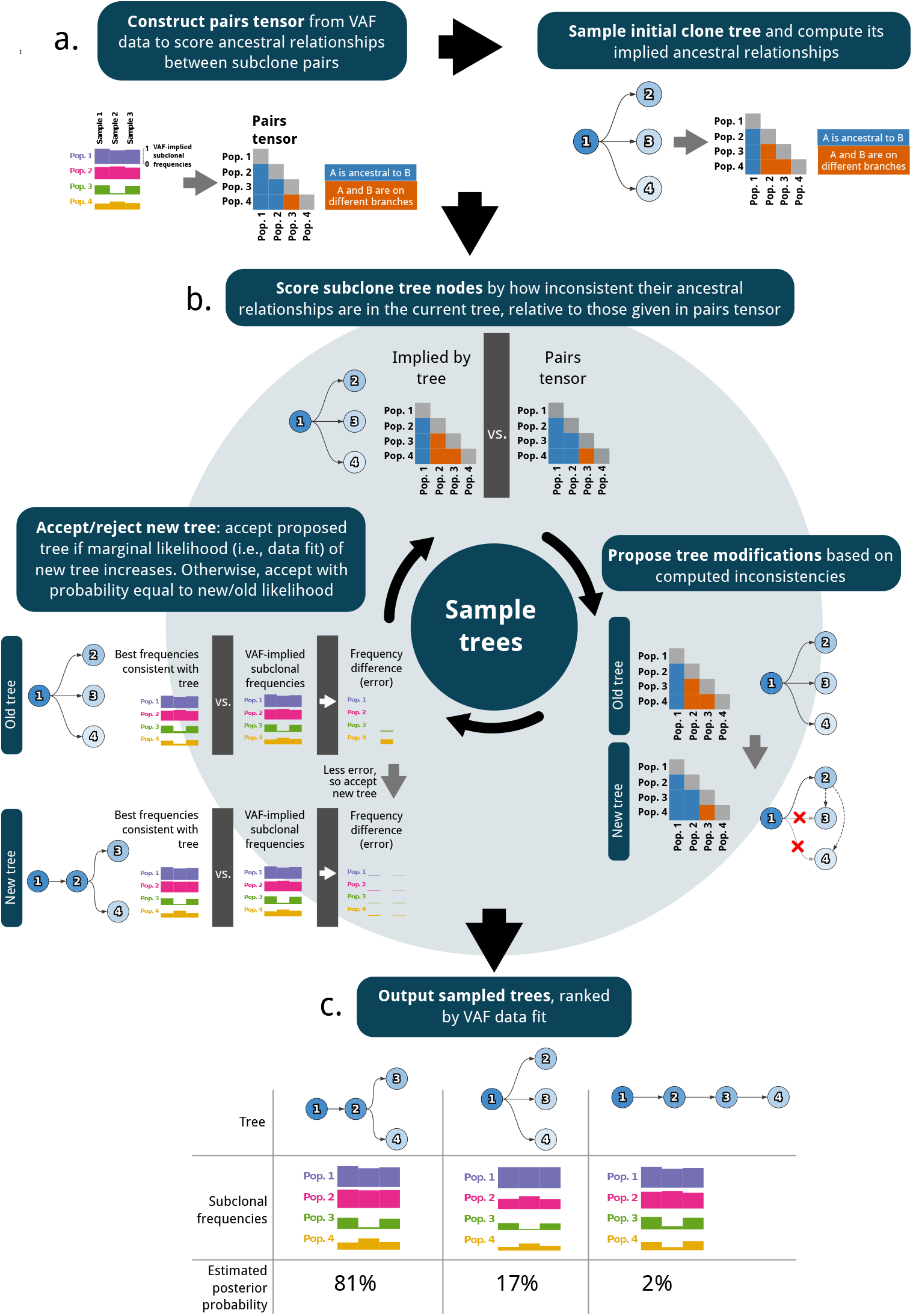
The Pairtree algorithm. **a.** Pairtree uses VAF data to compute the Pairs Tensor. This tensor denotes the probability of every possible pairwise ancestral relationship between subclones (left). An initial clone tree is built using relationships scored by the Pairs Tensor. **b.** Pairtree samples trees using Markov Chain Monte Carlo. The method proposes tree modifications by identifying a subclone whose ancestral relationships in the current tree are assigned low probability by the Pairs Tensor (top), then ascertaining where that subclone can be moved within the tree to increase its ancestral relationship probabilities (bottom right). Proposed trees are then accepted or rejected based on their likelihoods that reflect how well they fit the VAF data (bottom left). **c.** Sampled clone trees are returned along with posterior probability estimates proportional to the likelihood of each tree. *VAF*, variant allele frequency.

Pairtree uses a maximum a posteriori (MAP) approximation of the clone tree’s marginal likelihood, both for the Metropolis-Hastings accept-reject decision and for estimating the tree’s posterior probability. The Bayesian prior enforces tree constraints but is otherwise uninformative. By this constraint, the root subclone must have a subclonal frequency of 1 in every sample, as it corresponds to the germline and all subclones are descended from it. Additionally, the prior requires that every subclone has a frequency greater than or equal to the sum of its direct descendants’ subclonal frequencies. Pairtree can compute the MAP estimate either using a fast approximate scheme [41] or a slower exact one (Section 9.4). A clone tree’s likelihood scores how well the variant and reference read counts for each mutation match the MAP subclonal frequencies under a binomial sequencing noise model that includes the provided CNA correction for the mutation.

### 2.4 Benchmarking Pairtree performance using novel scoring metrics

Evaluating Pairtree against other common subclonal reconstruction methods required developing new metrics, as existing metrics are limited to datasets with single cancer samples [24], do not consider uncertainty about the best-fitting clone tree [35], or both. Below, we introduce two novel metrics well-suited for the multi-sample domain that also permit uncertainty about the best-fitting clone tree.

The first, termed *VAF reconstruction loss*, uses likelihood to compare the data fit of a tree’s subclonal frequencies to a baseline (Section 9.9.2). For simulated data, the baseline frequencies are the ground-truth frequencies used to generate the VAF data. For real data with an unknown ground truth, the baseline is MAP subclonal frequencies computed for an expert-constructed clone tree. If a method outputs multiple clone trees, the VAF reconstruction loss of this solution set is the average loss of each clone tree, weighted by the likelihood the method associated to the tree. Negative VAF losses indicate the evaluated frequencies have better data fit than the baseline. Importantly, this is an unbiased metric can be used even when the ground-truth is unknown, or when the simulated data supports a better-fitting clone tree than the one to generate it in the first place.

The second evaluation metric, termed *relationship reconstruction error*, compares the structure of candidate clone trees to the ground truth (Section 9.9.3) using the evolutionary relationships between subclone pairs. This metric is a generalization of previous pairwise-relation-dependent metrics [24, 35] to permit the comparison of distributions over clone trees to one another. The metric permits uncertainty in the ground truth clone tree while also rewarding methods that report multiple clone trees when the correct solution is indeed uncertain. To compute it, we construct an empirical Pairs Tensor from the clone tree solutions found by a method, then compare it via the Jensen-Shannon divergence (JSD) to a tensor based on the ground truth. As multiple clone trees may be consistent with the ground-truth subclonal frequencies, we construct the ground-truth Pairs Tensor by enumerating all trees consistent with these frequencies [42] and denoting the pairwise relationships between subclones that each expresses. Building the ground-truth collection of clone trees requires knowing the ground-truth subclonal frequencies with no measurement error, so this metric is best suited to simulated data.

### 2.5 Selecting comparison methods and generating simulated data

Clone tree reconstruction methods use one of two approaches: exhaustive enumeration or stochastic search. To evaluate Pairtree, a stochastic search method, we compared it against four exhaustive enumeration methods (CALDER [43], PASTRI [23], CITUP [19], and LICHeE [22]) and one stochastic search method (PhyloWGS [40]). All methods produce multiple candidate clone trees that are scored based on how well their tree-constrained subclonal frequencies fit the observed VAF data (see also [44]).

We assessed method performance on 576 simulated datasets with variable read depths and numbers of subclones, cancer samples, and mutations. These included trees with 3, 10, 30, and 100 subclones. Three subclones are the most that can typically be resolved at WGS read depths of 50x [1]. In multi-sample datasets, ten subclones are often discernible [6], while 30 was the approximate maximum we could resolve in the high-depth, many-sample B-ALL data evaluated here [11]. We also included trees with 100 subclones to probe the methods’ limits, anticipating challenges presented by future datasets. The number of simulated cancer samples ranged from 1 to 100. We designed the simulation process (Section 9.8.2) to generate realistic, diverse, and resolvable clone trees (Section 9.17). We did not include one- or three-sample datasets in the 30- and 100-subclone simulations, as resolving so many subclones from so few samples would be unrealistic. Methods were allowed up to 24 hours of wall-clock time to produce results.

Some caveats must be noted. LICHeE does not report subclonal frequencies for its solutions, so we used Pairtree to fit MAP frequencies to LICHeE’s trees. Though LICHeE does not produce a likelihood, unlike the other methods here, it reports an error score for each tree that we interpreted as a likelihood when weighting its solutions. PhyloWGS, unlike other methods, could not use a fixed mutation clustering. This led to the method incorrectly merging clusters, causing artificially high VAF loss and relationship error. More generally, all methods except Pairtree failed to produce output on some simulated datasets. These failures stemmed from methods terminating without producing output, crashing outright, or failing to finish within 24 hours (see Section 9.11 for details).

### 2.6 Pairtree outperforms existing methods on simulated data

Fig. 3 summarizes how the methods performed on simulated data, with a method’s scores reflecting its performance on only the datasets for which it produced output. Pairtree was the only method that produced results for all 576 simulations (Fig. 3a). Nevertheless, Pairtree fared better than comparison methods on trees with 30 or fewer subclones, succeeding on all datasets while achieving negative median VAF losses (Fig. 3b-c). In fact, Pairtree always produced lower error than other methods for every such dataset (Fig. S3), except for two datasets with three subclones and a single cancer sample where CALDER had negligibly better VAF losses (i.e., 0.002 bits lower or less). Pairtree also performed better than comparison methods with respect to relationship error. In general, for 30 subclones or fewer, relationship error was almost zero when the number of cancer samples exceeded the number of subclones (Fig. S5b). For these cases, only one clone tree fit the ground-truth subclonal frequencies (S14a) and Pairtree achieved low error by finding that tree or a close approximation thereof (S14b-c). When applied to datasets with 100 subclones, Pairtree had higher VAF losses (Fig. 3b) and relationship errors (Fig. 3c) than with fewer subclones. Pairtree outperformed other methods for 100-subclone trees with respect to VAF loss, except for 16 datasets (15%) where PhyloWGS performed better (Fig. S3) and 22 where CALDER was better.

**Figure 3:**
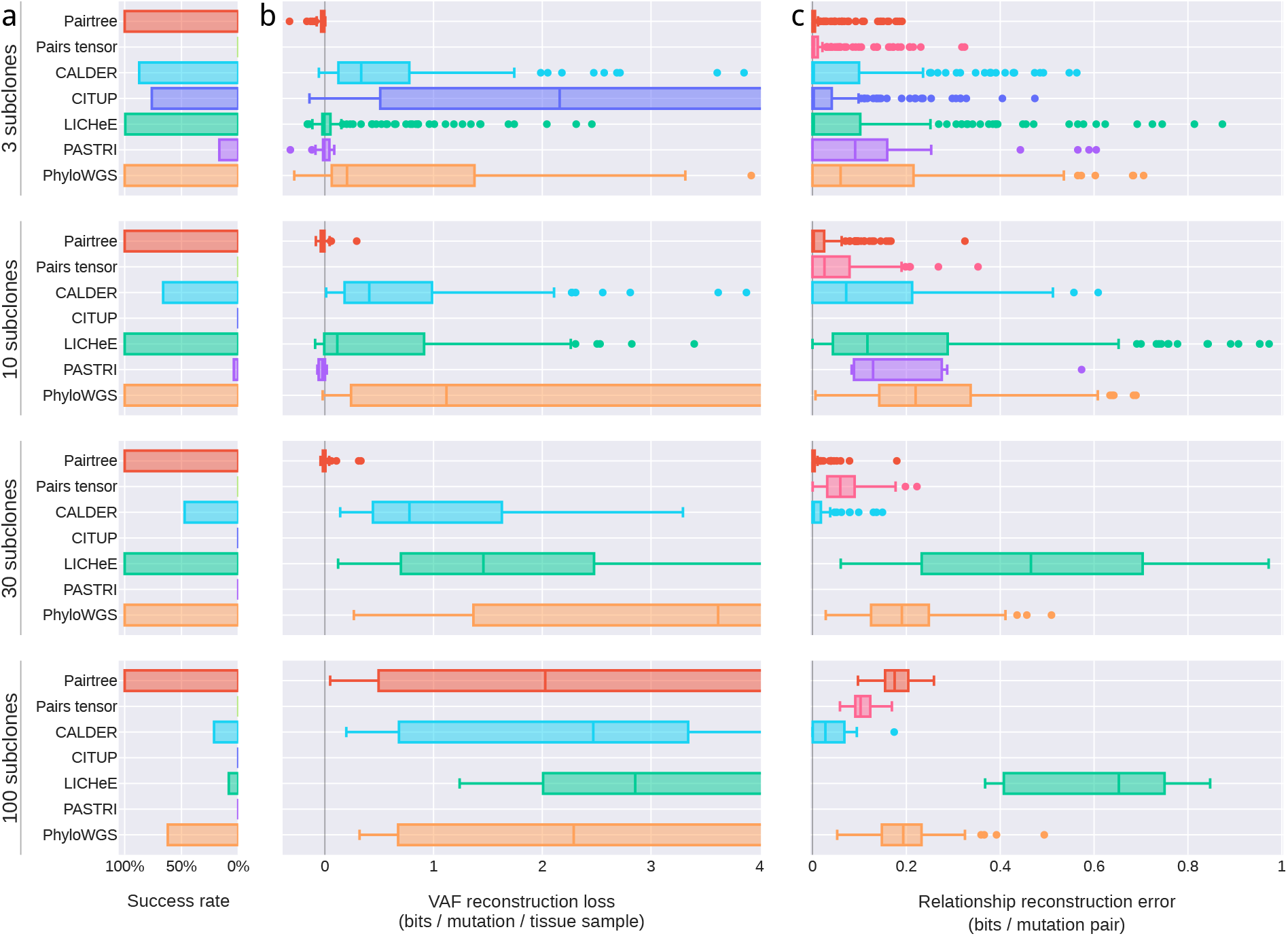
Benchmark performance on 576 simulated datasets. Simulations are grouped by number of subclones (rows). **a.** Bar plots show each method’s success rate in the group. Successes are reconstruction problems for which the method produced at least one tree in 24 hours (wall-clock time) and did not crash. **b.** Boxplots show distributions of VAF reconstruction losses for a method on a problem group. Scores reflect only datasets where a method ran successfully. VAF reconstruction loss is the decrease in average, per-mutation log likelihood of VAF data using subclonal frequencies assigned by the method, when compared to the true frequencies used to generate the data. Negative loss indicates better VAF reconstructions than true trees, while high loss indicates inaccurate tree structures. Mid-lines in box plots indicate medians. Plots are truncated at four bits. Fig. S1 shows untruncated distributions. **c.** Boxplots show distributions of relationship reconstruction error in each group for each method’s successful runs. Relationship reconstruction error is measured as the average Jensen-Shannon divergence per subclone pair between the true distributions over pairwise relations, and empirical distributions computed from the trees output by a method. Errors can range between zero bits (perfect match) and one bit (complete mismatch).

CITUP failed on all datasets with ten or more subclones, and on 32% of three-subclone cases (Fig. 3a). All failures on three-subclone datasets occurred because CITUP crashed (Section 9.11). On ten-subclone datasets, 29% of CITUP runs ran out of time, with the other 71% failing because CITUP crashed. On the three-subclone cases where it ran successfully, its VAF loss was poor (Fig. 3b), perhaps because of a mismatch between its sequencing error model and the model used for computing VAF loss. Conversely, the method exhibited better relationship error than other non-Pairtree methods (Fig. 3c), suggesting its tree structures were more accurate.

PASTRI, which cannot run on datasets with more than 15 subclones [35], failed for 83% of three-subclone cases and 96% of ten-subclone cases (Fig. 3). For datasets with three or ten subclones, PASTRI produced output on 10%, terminated without producing a result on 84%, and ran out of time on the remaining 6% (Section 9.11). When it produced solutions, PASTRI generally performed well, reaching negative median VAF losses for three- and ten-subclone datasets, and relatively low relationship errors.

LICHeE fared better, producing results on all cases with 3, 10, or 30 subclones (Fig. 3). However, the method ran out of time for 92% of 100-subclone datasets. After Pairtree, LICHeE was the next-best performing method, with low VAF losses and moderate relationship errors on datasets with three or ten subclones, beating PhyloWGS on both measures. LICHeE performed less well on 30-subclone cases, where it exhibited lower VAF losses than PhyloWGS but higher relationship errors.

PhyloWGS produced clone trees for all datasets with 30 or fewer subclones (Fig. 3). In these cases, PhyloWGS generally had worse VAF losses and relationship errors than Pairtree or LICHeE, except for the 30-subclone datasets, where it had better relationship error than LICHeE but worse VAF loss. PhyloWGS performed better than other non-Pairtree methods on 100-subclone cases, where it finished within 24 hours for 62% of such datasets, but usually had higher VAF losses than Pairtree (Fig. S3).

CALDER in its non-longitudinal mode failed on 13% of three-subclone cases, 34% of ten-subclone cases, 53% of 30-subclone cases, and 79% of 100-subclone cases (Fig. 3). On datasets with 30 or fewer subclones where it succeeded, CALDER generally produced VAF losses lower than PhyloWGS and on par with LICHeE, and relationship errors that were better than all non-Pairtree methods. On the 21% of 100-subclone cases where it produced a result, CALDER exhibited performance that was generally the best of all methods, achieving lower VAF loss than Pairtree on 22 of the 108 datasets with 100 subclones.

Relationship error can also be measured for the Pairs Tensor alone, without requiring trees. The Pairs Tensor estimates pairwise relationships well (Fig. 3c), requiring only a fraction of the computational resources of the full Pairtree method (Fig. S12). Although the Pairs Tensor does slightly worse than Pairtree on trees with 30 or fewer subclones, it has less relationship error than any other method. On datasets with 100 subclones, the Pairs Tensor was better able to delineate pairwise relationships between subclones than the full Pairtree method (Fig. 3c).

### 2.7 Pairtree improves with more cancer samples, but other methods worsen

After controlling for other variables, all methods except Pairtree performed worse when provided more cancer samples. CITUP and PASTRI’s failure rates increased with the number of cancer samples (Fig. 4a). Though LICHeE and PhyloWGS produced output for all cases with 30 subclones or fewer, they had higher VAF losses with more cancer samples (Fig. 4b). By contrast, Pairtree never failed and had nearly zero median VAF loss regardless of the number of simulated cancer samples on datasets with 30 subclones or fewer (Fig. 4a-b). Relationship errors decreased for both full Pairtree and the Pairs Tensor with more samples (Fig. 4c). LICHeE, conversely, exhibited rapidly increasing error with more samples, while PhyloWGS’ performance fluctuated. Because CALDER failed on subsets of all datasets when partitioned by number of subclones, we consider it separately. CALDER generally failed more frequently as the number of subclones or number of cancer samples increased (Fig. S8), while its VAF loss was largely independent of the number of subclones (Fig. S9).

**Figure 4:**
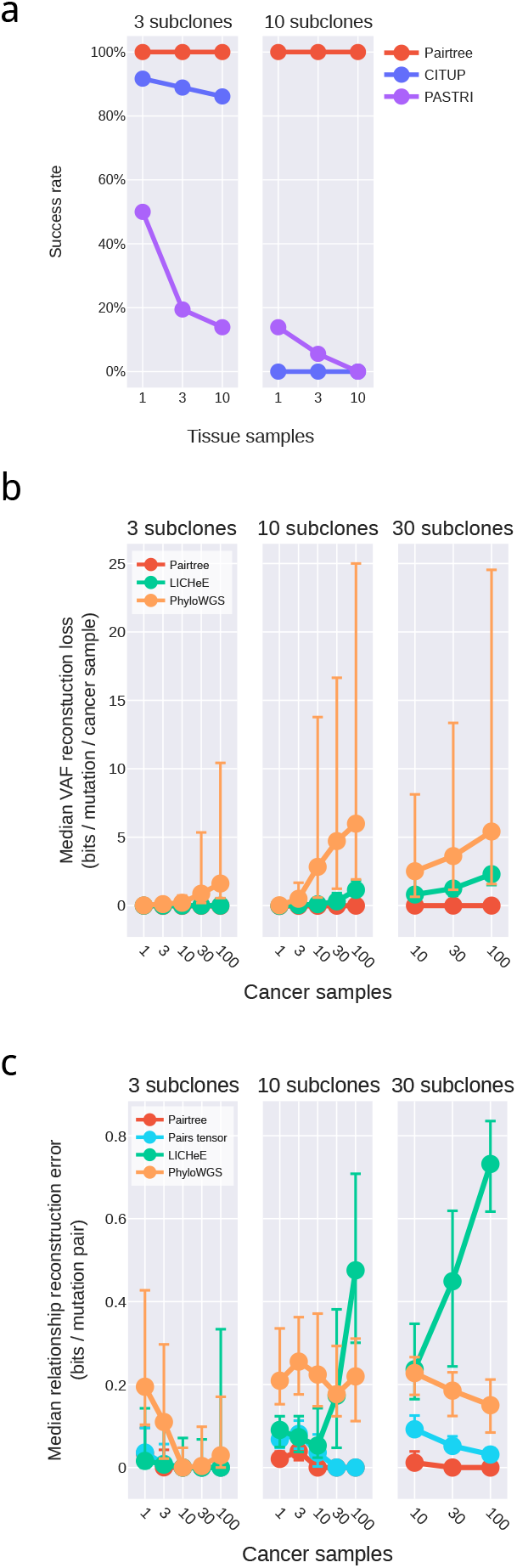
Performance on simulated datasets as a function of number of subclones and cancer samples. CALDER is not shown because it succeeded only on subsets of the different dataset groups shown, while the methods represented here succeeded on all datasets in the depicted groups. **a.** Method success rate. For CITUP and PASTRI, success rate depended on the number of subclones and/or cancer samples in datasets. Pairtree, LICHeE, and PhyloWGS succeeded on all datasets depicted. **b.** Median VAF reconstruction loss as a function of number of samples. For LICHeE and PhyloWGS, VAF loss increases with more cancer samples. **c.** Median relationship reconstruction error as a function of number of samples. LICHeE’s error generally increased with more cancer samples, while other methods showed the opposite effect. Error bars represent the first and third quartiles in (b-c).

### 2.8 Pairtree accurately detects mutations that violate the Infinite Sites Assumption

Pairtree’s Pairs Tensor can be used to identify mutations that violate the ISA (see Section 9.6). Figs. S19 to S21 show the performance of Pairtree’s ISA violation detection algorithm under four different scenarios: technical noise (i.e. sequencing artifacts), homoplasy, back mutation, and miscalled CNAs. This evaluation is performed under two different strengths of evidence, encompassing strong support (i.e., a 5% difference in VAF, implying a 10% difference in subclonal frequency) and weak support (i.e., a 0.025% difference in VAF, implying a 0.05% difference in subclonal frequency). We also assessed ISA-violation detection for clone trees with 10 and 30 subclones. Pairtree has 100% precision and recall for simulated sequencing artifacts in all scenarios, and its precision for finding ISA-violating mutations does not drop below 99% for the other three three cases. Its recall exceeds 97% for all cases within these three except for the weak-support case on 30-subclone trees, where its recall drops to 88% for homoplasy and back mutation. This demonstrates that Pairtree can detect ISA violations nearly perfectly in most scenarios, save for two where its detection is still excellent.

### 2.9 Pairtree performs better than human experts on complex real clone tree reconstructions

We applied Pairtree, CALDER, CITUP, LICHeE, PASTRI, and PhyloWGS to genomic data from 14 B-ALL patients [11]. Samples were obtained at diagnosis and relapse for each patient. In addition, each sample was transplanted into immunodeficient mice, generating multiple patient-derived xenografts (PDXs). The patient samples were profiled using WES, while the PDXs were used targeted sequencing based on leukemic variants found in the patient WES data. There were 16 to 509 mutations called per patient (median 40), clustered into 5 to 26 subclones per patient (median 8). By combining patient and PDX samples, we obtained between 13 and 90 tissue samples per cancer (median 42). Across cancers, the median read depth was 212 reads.

To define an expert-derived baseline for these datasets, we first built high-quality clone trees for each dataset manually, subjecting them to extensive review and refinement before evaluating them for biological plausibility [11]. Then we use the same technique as Pairtree to fit tree-constrained subclonal frequency estimates to the VAF data. The data fit of these estimates, as computed by likelihood, yields the *expert-derived baseline.* As with simulated data, methods that improve on the baseline achieve negative VAF losses.

CITUP and PASTRI failed on 13 of the 14 cancers, and so we excluded these methods from the comparison. Pairtree found trees that fit the sequencing data as well as, or slighter better than, the expert baseline for 12 of 14 cancers (Fig. 5), resulting in VAF losses between 0 and −0.05 bits. On two cancers, Pairtree inferred clone trees that fit the VAF data substantially better than the expert baseline, resulting in negative losses of −0.32 bits and −1.42 bits. LICHeE beat the baseline for one cancer, reaching a negative loss of −0.86 bits; (nearly) matched the baseline for four other patients, incurring between 0 and 0.11 bits of loss; and had substantially worse VAF losses for the remaining nine patients. PhyloWGS suffered at least 0.35 bits of loss on all patients, reaching a median VAF loss of 4.42 bits. As PhyloWGS could not adhere to the expert-derived clustering, unlike other methods, it often merged clusters incorrectly, causing high VAF loss. CALDER failed on 3 of the 14 cancers, was worse than Pairtree on all the other 11, and worse than LICHeE on 9 of the 11.

**Figure 5:**
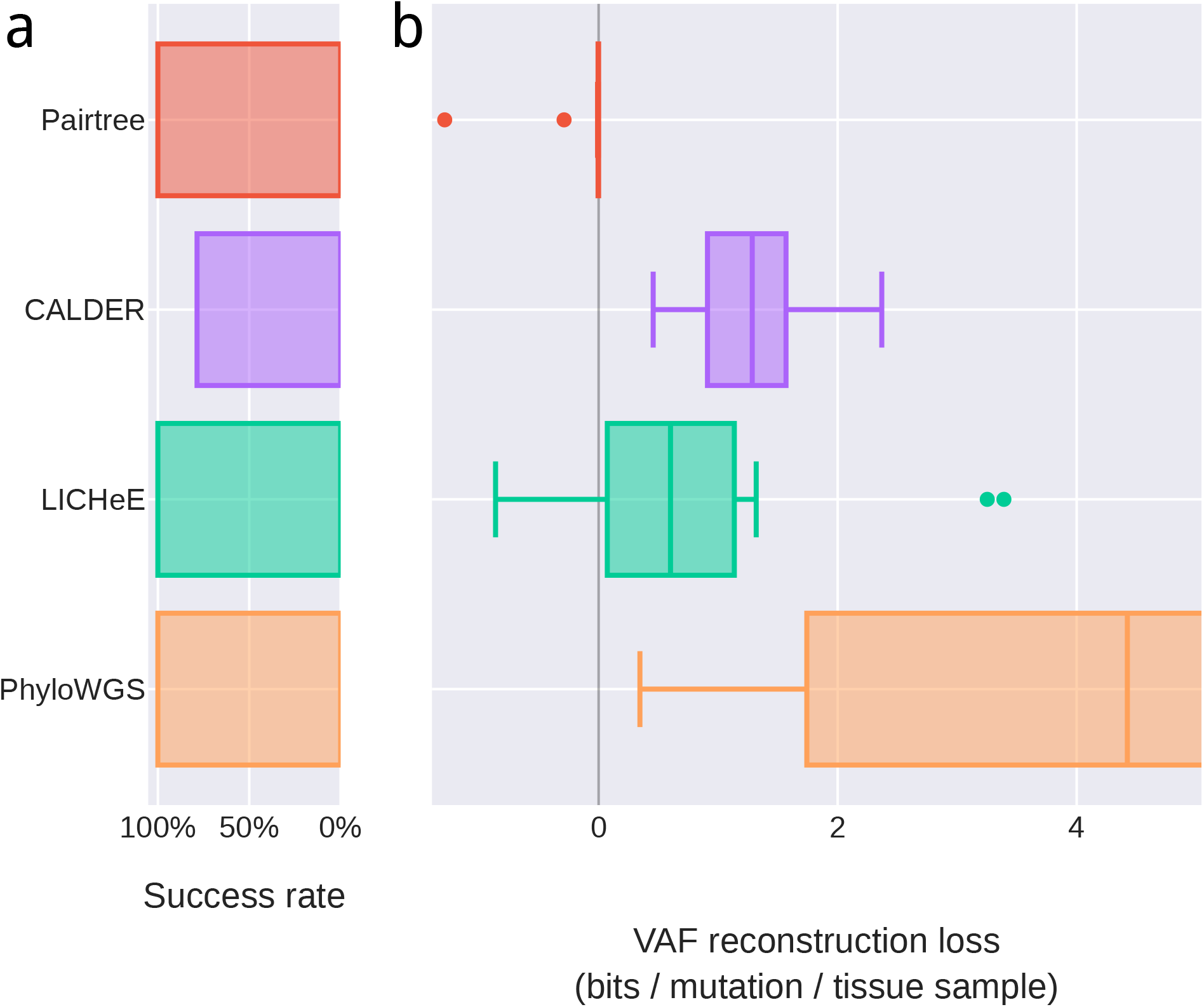
Method performance loss for 14 B-ALL patient datasets. The number of cancer samples for each dataset ranged from 13 to 90. **a.** Pairtree, LICHeE, and PhyloWGS succeeded on all 14 datasets. CALDER succeeded on only 11 of the 14 (79%). CITUP and PASTRI each failed on 13 of 14 datasets and so are not shown. **b.** VAF loss on the subset of datasets where each method succeeded. VAF reconstruction losses are reported as a negative log likelihood normalized to the number of mutations and cancer samples, relative to the MAP subclonal frequencies for expert-derived trees. Lower loss indicates better performance, while negative loss corresponds to performance better than human experts. Mid-lines in box plots indicate medians. The axis is truncated at 5 bits. Fig. S2 shows untruncated distributions.

### 2.10 Consensus graphs intuitively illustrate uncertainty in clone trees

Pairtree provides interactive visualizations to help navigate the multiple clone tree solutions that it produces for each dataset (Fig. 6). By using the likelihoods associated with each solution as weights, Pairtree produces a *weighted consensus graph*, in which the nodes represent subclones, and each directed edge is assigned a weight equal to the marginal probability that it appears in a clone tree drawn from the empirical clone tree distribution produced by Pairtree. Thus, the consensus graph summarizes the estimated posterior probability of each parental relationship between subclones. These summaries are useful for interpreting Pairtree’s results, as they provide a concise representation of the evolutionary relationships supported by the data, alongside the confidence underlying each. By taking the maximum-weight spanning tree of this graph, the user can generate a single consensus tree. To demonstrate the consensus graph’s utility, we ran Pairtree multiple times on one of the B-ALL cases from Fig. 5, using variable numbers of cancer samples (Fig. 6). As we provided more cancer samples, confidence in evolutionary relationships increased, until all parents were resolved with near certainty. Providing more samples can also correct erroneous inferences—with 30 samples, population 8 appeared to be the likely parent of population 15, but with 90 samples, it became clear that population 15’s parent is population 6.

**Figure 6:**
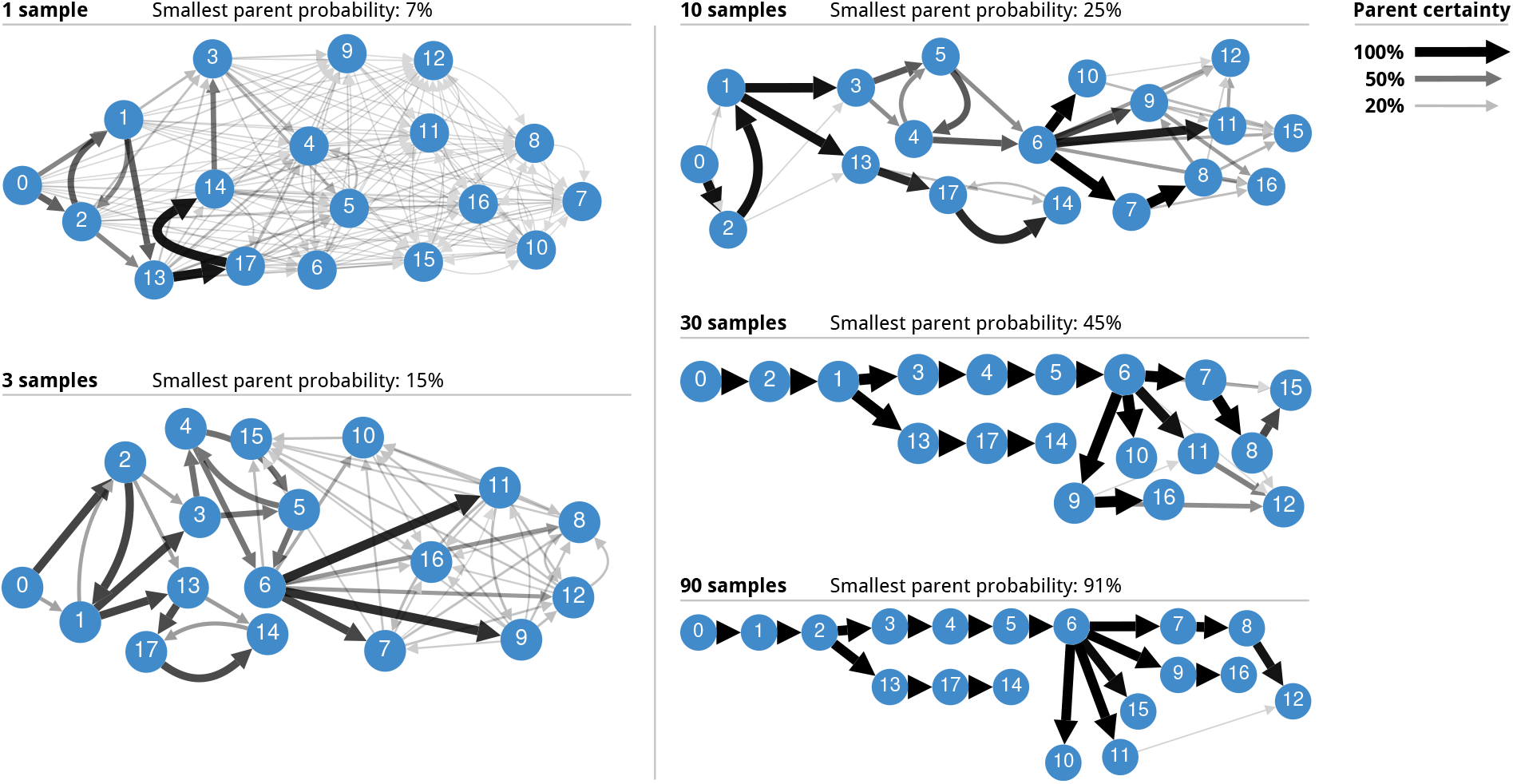
Consensus graph visualization of posterior tree distributions. These consensus graph visualizations are based on one of the 14 B-ALL cancers analyzed with Pairtree, for which 90 cancer samples were available. Consensus graphs are shown for variable numbers of samples, ranging from a single sample to all 90. All edges with less than 5% posterior probability are hidden. The minimum spanning tree certainty is the minimum of the maximum parent probabilities of each subclone.

## 3 Discussion

Pairtree is the first automated method that reliably recovers large, complex clone trees from bulk DNA sequencing data. For simulated clone trees with up to 30 subclones, Pairtree’s reconstructed clone trees almost always fit the VAF data as well as or better than the original clone trees used to generate the data. On 14 B-ALL cancers, with up to 26 subclones and 90 samples per cancer, Pairtree’s clone trees fit the VAF data as well as, or better than, those constructed by experts. No other tested method was consistently accurate on real or simulated benchmarks containing ten subclones or more. Pairtree was also the only method whose clone trees reliably became more accurate when more samples were used in the reconstructions. This is surprising—as each cancer sample provides additional information about evolutionary relationships between subpopulations, subclonal reconstruction problems should become easier with more cancer samples, not more difficult.

Identifying the correct clone tree for a given dataset may not be possible, and so Pairtree is specifically designed to identify and report ambiguities in the clone tree reconstruction. For example, the relationships among subclones with low VAFs in all samples may not be possible to resolve because, depending on the read coverage, the low VAFs might be consistent with multiple ancestral relationships between the subclone pair. In these circumstances, Pairtree is designed to capture this uncertainty in both its Pairs Tensor and through its MCMC-derived samples from the clone tree posterior. Additionally, due to incomplete genome coverage or the inherent sequencing limits of mutation detection, all clone trees provide an incomplete view of a cancer’s evolution. Accounting for uncertainty under these conditions becomes even more important because of the difficulty of resolving the true tree.

A key factor in Pairtree’s success is its efficient search through the space of clone trees. Beyond ten subclones, this tree space quickly becomes too large for exhaustive enumeration (CITUP) or unguided stochastic search (PhyloWGS). Even methods that reduce the search space by applying hard constraints to exclude some parent-child relationships (LICHeE, PASTRI, CALDER) can fail to recover clone trees with more subclones because as the number of samples increases, these hard constraints become more likely to be incorrect and thus exclude the correct solution (Section 9.14). By contrast, Pairtree’s stochastic tree search is guided by the Pairs Tensor, which provides soft constraints defined by a well-motivated probability model. Consequently, Pairtree’s constraints become more precise as more cancer samples are provided, without excluding the true clone tree.

As Pairtree’s performance degrades on the 100-subclone benchmarks, alternative search strategies may be necessary for very large clone trees. While Pairtree almost always fails to correctly resolve a subclone’s parent (Fig. S14c), it achieves relatively low relationship error (Fig. S14d), suggesting it may be capturing the coarse tree structure. If so, Pairtree may fare better using a tiered approach, in which it would group together subclones with similar pairwise relations to others, build subtrees for each group separately, and then connect the subtrees using the groups’ pairwise relations to compose the full clone tree. Given 100 subclones with 10 or more cancer samples, the Pairs Tensor is already better than Pairtree itself at capturing the correct evolutionary relationships between subclones (Fig. S5b-c). Future work should focus on understanding what conditions (e.g., high read depth or many cancer samples) under which the Pairs Tensor converges to a partial clone tree [42] that succinctly summarizes all clone trees with non-negligible posterior probability.

Future extensions of Pairtree could incorporate alternative models of cancer evolution by introducing non-uniform priors on clone tree structure or subclonal frequencies. Any evolutionary model that can assign a likelihood to a given clone tree structure and set of subclonal frequencies can be immediately incorporated in the Metropolis-Hastings scoring, though some subclonal frequency priors may make the subclonal frequency optimization non-convex (see Section 9.4.3) or reduce the accuracy of the MAP approximation to the marginal likelihood of the tree (see Section 9.4.4). Additionally, non-uniform priors may decrease the value of the Pairs Tensor as a proposal distribution for tree inference, but, encouragingly, any prior that permits tractable computation of their marginal distributions over subclone pairs can also be incorporated into the Pairs Tensor. For example, CALDER’s longitudinal constraints (i.e., once a subclone goes extinct at a given timepoint, it never returns) [43] can be incorporated as time-dependent priors on subclonal frequencies for individual subclones and subclone pairs where one is the ancestor of the other. Section 9.18 describes some possible alternative evolutionary models in detail.

Throughout this work, we have stressed performance metrics that recognize there are often many solutions consistent with observed data (Section 9.16), extending previous ones that compare single clone trees to one another (see [23, 43] and others). We developed new metrics that extend ones we previously developed [24] to score multiple candidate solutions from a method against a single ground-truth tree. Our new metrics permit the ground truth to be uncertain, with multiple potential truths equally consistent with noise-free observations. In general, characterizing uncertainty in clone tree reconstructions is critical. Even when methods produce multiple solutions, users typically want a single answer, and so select the highest-scoring tree while neglecting other credible candidates that fit their data nearly as well. Consequently, they lose information about which evolutionary relationships between subclones are well-defined by the data, and which are uncertain because they have multiple equally likely possibilities. If users are to benefit from a method’s ability to produce multiple solutions, the method must provide tools for interpreting this uncertainty. Pairtree’s weighted consensus graph characterizes the uncertainty present in each evolutionary relationship, depicting all credible possibilities and the confidence underlying each (Fig. 6). This allows users to make informed conclusions about their data.

In summary, Pairtree can reconstruct highly accurate trees representing the evolutionary relationships among up to 30 subclones based on sequencing data from up to 100 samples from a cancer. Using pairwise mutation relationships, Pairtree can detect mutations that violate the ISA (Section 9.6) or have technical issues corrupting their observed data. By scaling to many more subclones and cancer samples than past approaches, and by illustrating the uncertainty present in solutions, Pairtree can address questions in many cancer research domains. These include understanding the origin and progression of tumours, measuring tumour age and heterogeneity, mapping out mechanisms of tumour adaptation to therapy, and understanding the relationship between primaries and metastases. Pairtree also has applications beyond cancer, where it can be used to examine somatic evolution in non-cancerous tissues for any asexually-dividing cell population. In the future, the Pairtree framework can be extended to scale to even more complex trees, integrate single-cell sequencing data (Section 9.7), and permit violations of the infinite sites assumption (Section 9.6.1).

## 5 Methods

### 5.1 Structure

A succinct summary of Pairtree is provided here. Section 9.2 through Section 9.7 provide an expanded version of these concise methods.

### 5.2 Pairtree input

Pairtree requires (*V_js_, R_js_*), the variant and reference read counts, respectively, and *ω_js_*, the subclonal frequency to VAF conversion factor, for each mutation *j* ∈ {1, 2, …, *J*} in each sample *s* ∈ {1, 2, …, *S*}. Pairtree also can take as input a grouping of mutations into subclones. For each subclone *k*, the associated set of mutations *S_k_* ⊂ {1, 2, …, *M*} is used to define a “supervariant” representing the set (see below). When asked to cluster mutations itself, Pairtree also computes a supervariant for each subclone. The VAF conversion factor is defined as *ω_js_* = *A_js_/N_js_*, where *A_js_* is the average number of alleles containing mutation *j* in cells that contain *j* in sample *s*. Typically *A_js_* = 1, unless there is a subclonal copy number aberration (CNA) that causes a gain in copy number (or a loss) of the allele containing *j* [40]. In this circumstance, unless *A_js_* can be accurately estimated, we suggest removing mutation *j* from the input. In contrast, the value of *N_js_* is the average copy number per cell of the locus containing *j*, and it can often be estimated directly from relative read depth, or if CNA reconstruction is available, then in autosomal regions, *N_js_* = 2 + *ρ*(*κ* – 2), where *ρ* is the proportion of cells with the CNA and *κ* is the new CN. In areas of normal CNA, 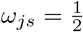 if autosomal, and *ω_js_* = 1 if haploid (e.g., male sex chromosomes).

A supervariant *k* is an artificial mutation representing subclone *k*. The supervariant is associated with a set of values (*V_ks_, R_ks_, ω_ks_*) whose likelihood, as a function of the subclonal frequency *φ_ks_*, is proportional to the product of the likelihoods of all the mutations *j* ∈ *S_k_*, provided those mutations have the same *ω_js_* values. The supervariant thus allows us replace all calculations considering the set of mutations in *S_k_* with a single calculation on the supervariant *k*. Assuming that the values of *ω_js_* are equal for all *j* ∈ *S_k_*, we can set *ω_ks_* to that shared value and then set *V_ks_* ∑_*j*∈*S_k_*_ *V_js_* and *R_ks_* ∑_*j*∈_*S_k_*__ *R_js_*. Otherwise, if not all the *ω_js_*’s value are equal, we must compute a *j*-dependent correction for each mutation as part of computing *V_ks_* and *R_ks_* (see Section 9.3.8). Henceforth, we will assume that all subclonal clusters have been replaced with supervariants and thus that all computations are on single mutations or between pairs of mutations. We also assume that any ISA-violating mutations have been filtered out before the supervariants are defined (Section 9.6 describes Pairtree’s mutation filtering algorithm).

### 5.3 Computing the Pairs Tensor

The Pairs Tensor consists of a set of posterior probability distributions, for each mutation pair *i* and *j*, over their possible pairwise evolutionary relationships, represented by *θ_ij_*. In addition to the three relationships defined in Section 2.2 (*θ_ij_* ∈ {*ancestor, descendant, branched*}), Pairtree’s ISA-violation detection requires a fourth relationship, *garbage*. Additionally, one of Pairtree’s clustering algorithms makes use of a fifth relationship, *coincident* (see Section 9.5.3). To compute these posteriors, we use a uniform prior 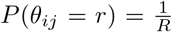 for each relationship *r* given *R* total relationships, and a data likelihood of all data, *x_i_* and *x_j_*, associated with mutations *i* and *j*, respectively. As the samples are exchangeable, we can write

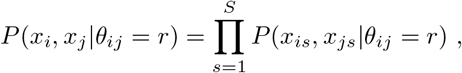

where *x_is_* (*x_js_*) represents the data associated with mutation *i* (*j*) in sample *s*. To compute the persample data likelihoods, we integrate out the subclonal frequencies *ϕ_is_* and *ϕ_js_* of *i* and *j*, respectively, i.e.,

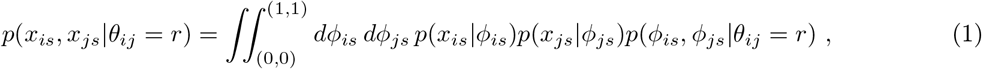

where *p*(*x_is_*|*ϕ_is_*) = Binom(*V_is_*|*T_is_*, *ω_is_ ϕ_is_*), for *T_is_* = *V_is_* + *R_is_*, and Binom(*V*|*N,p*) is the binomial likelihood of *V* given *N* trials with a success probability of *p*.

For each relationship r, the prior *P*(*ϕ_is_, ϕ_js_*|*θ_ij_* = *r*) enforces constraints on the values of *ϕ_is_* and *ϕ_js_*:

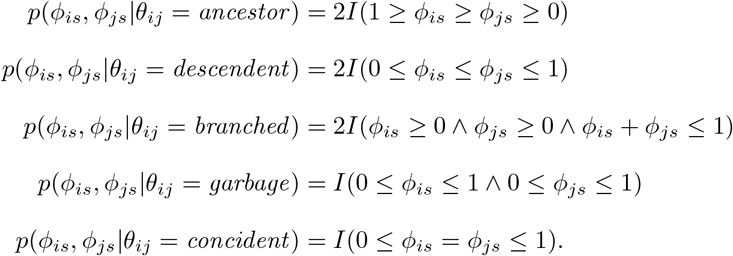

Here *I*(*B*) is the indicator function which equals 1 if statement *B* is true and 0 otherwise. Through algebraic manipulation, the 2-D integral in Eq. (1) can be converted into a 1-D integral that Pairtree computes numerically using quadrature (Section 9.2.6). The Pairs Tensor is used not only for building clone trees, but also for clustering mutations (Section 9.5.3) and detecting ISA violations (Section 9.6).

### 5.4 Constructing clone trees

Pairtree samples from a posterior distribution over clone trees using the the Metropolis-Hastings MCMC algorithm. After initializing tree search (Section 9.3.7), Pairtree uses the Pairs Tensor to propose a new tree by moving the location of one of the nodes in the current tree. These tree modifications are computed in two steps.

First, the algorithm samples a node (i.e., mutation) *b* to move from a probability distribution over nodes *q*(*b*|*t*), based on how unlikely *b*’s pairwise relationships in the current tree *t* are according to the Pairs Tensor. Let *p*(*θ_kb_*|*x_k_,x_b_*) be the posterior probability denoted in the Pairs Tensor of mutations *k* and *b* having pairwise relation *θ_kb_*. Then we define the pairwise relationship error to be

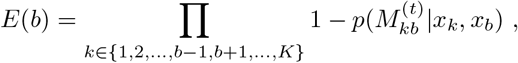

where 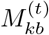 is the pairwise relationship of *k* and *b* in *t*. Then we create *q*(*b*|*t*) by transforming the vector *z* = (log *E*(1), log *E*(2), …, log *E*(*K*)) using a scaled softmax function ssmax(*z*) ≡ softmax(*w* * *z*), where *w* is a scalar chosen so that 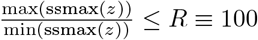. The *w* scalar is set to 1 if 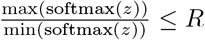, or otherwise to whatever value greater than 1 is necessary to make 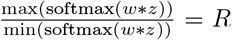. This ensures that every node has a non-negligible probability of being selected for modification.

The algorithm then chooses a destination for mutation *b* by sampling another node *a* from a probability distribution *q*(*a*|*b,t*) defined over all trees nodes except *b* and *b*’s current parent in tree *t*, denoted 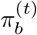. If *a* is not a descendant of *b*, the new tree *t*^(*b,a*)^ is generated by moving the subtree with *b* as its root so that *b* becomes a child of *a*. Otherwise, if *a* is a descendant of *b*, the positions of *a* and *b* are switched without altering other nodes. Like *q*(*b*|*t*), the distribution *q*(*a*|*b,t*) is defined using a vector *y* whose elements 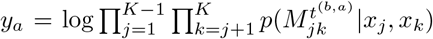, represent the “tree score” for tree *t^b,a^* under the Pairs Tensor. As with *q*(*b*|*t*), we use the scaled softmax with *R* = 100 to define *q*(*a*|*b,t*), so that *q*(*a*|*b,t*) = ssmax(*y*).

With the proposal tree *t*^(*b,a*)^ generated, we use the Metropolis-Hastings algorithm to accept the proposal with probability

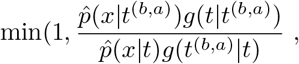

where the transition probability for the Metropolis-Hasting decision rule is *g*(*t*^(*b,a*)^|*t*) = *q*(*b*|*t*)*q*(*a*|*b,t*).

Note that if *t*′ = *t*^(*b,a*)^, then *t* = *t*′^(*a,b*)^, implying *g*(*t*|*t*^(*b,a*)^) = *q*(*a*|*t*^(*b,a*^))*q*(*b*|*a*, *t*^(*b,a*)^). Here, we approximate the likelihood *p*(*x*|*t*) for data *x* using the maximum likelihood estimate of Φ = {*ϕ_ks_*, ∀*k*, *s*} that satisfies the tree constraints of *t*. We define

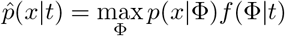

where *f*(Φ|*t*) = 1 if Φ satisfies the constraints imposed by *t* and 0 otherwise, and

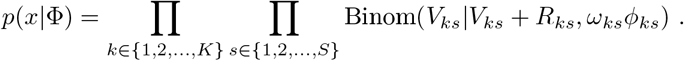

The maximum likelihood subclonal frequencies Φ are computed either exactly using gradient-based optimization, or it can be rapidly approximated based on a Gaussian approximation to the likelihood (Section 9.4). These frequencies must satisfy the following constraints:

1. *ϕ_ks_* ∈ [0,1] for all *k* and *s*.
2. *ϕ*_0*s*_ = 1 for all *s*, where 0 indexes the non-cancerous node that is the root of any clone tree.
3. The subclonal frequency for *k* must be at least as great as the sum of its childrens’ frequencies in *t*, i.e., 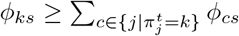.

### 5.5 Generating simulated data

We generated simulated data with four parameters:

- *K*: number of subpopulations
- *S*: number of cancer samples
- *M*: number of variants
- *T*: number of total reads per variant

We created simulated datasets with the following parameter combinations.

**Table 1:**
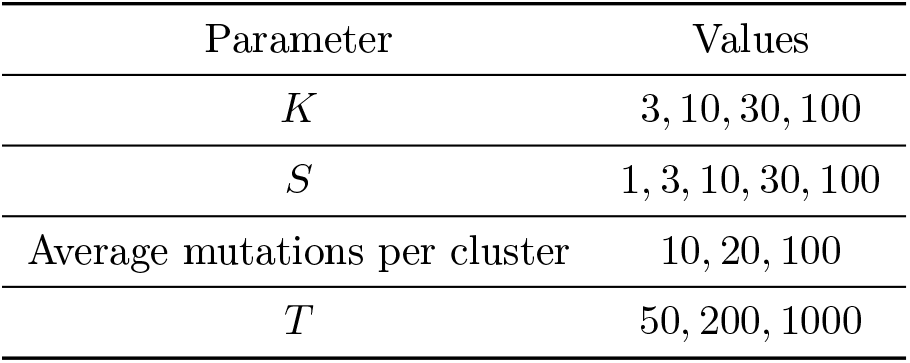
Simulated data parameters.

All combinations of these parameter values were used to generate simulated data, except cases when *K* ∈ {30, 100} and *S* ∈ {1, 3}. This provided 144 parameter combinations, with four datasets generated from each, yielding 576 simulated datasets. Using these parameters, we generated simulated datasets with the following procedure.

1. Generate the tree structure, *t.* For each subclone *k* ∈ {1, 2, …, *K*1}, sample a parent 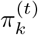. We selected as parent the previous subpopulation (i.e., 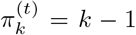) with probability *μ* = 0.75, and otherwise sample 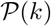 from the discrete Uniform (0, *k* – 1) distribution. This extension probability created “linear chains” of successive subpopulations, with each member of the chain taking only a single child, interrupted sporadically by the creation of new tree branches.
2. Generate the subpopulation frequencies *η_ks_* for each subpopulation *k* in each cancer sample *s*, with *s* ∈ 1, 2, …, *S*. These values were sampled separately for each *s*, with [*η*_0*s*_, *η*_1*s*_, …, *η_Ks_*] ~ Dirichlet(*α***1**), using *α* = 0.1, where **1** is a vector of 1’s
3. Compute the subclonal frequencies *ϕ_ks_* for each subclone *k* in each cancer sample *s* based on *t* and *η_ks_* values, such that

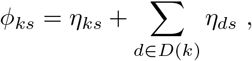

where *D*(*k*) denotes the set of descendants of *k* according to the tree structure.
4. Assign the *M* mutations to subclones. To ensure every subclones has at least one mutation, set the subclones of the first *K* mutations to 1, 2, …, *K*. To assign the remaining *M* – *K* mutations, sample subclone weights from the *K*-dimensional Dirichlet(1, 1, …, 1), then sample assignments from the *K*-dimensional categorical distribution using these weights.
5. Sample read counts for the variants. Let *A*(*m*) ∈ {1, 2, …, *K*} represent the subclone to which variant *m* was assigned. Let 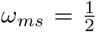 represent the probability of observing a variant read when sampling reads from the variant’s locus, for all subpopulations contained within *m*’s subclone, reflecting a diploid variant not subject to any CNAs. Then, for each cancer sample *s*, given the fixed total read count *T* used for all variants in a dataset, we sample the number of variant reads *V_ms_* ~ Binomial(*T,ω*_*ms*_*ϕ*_*A*(*m*),*s*_).

### 5.6 VAF reconstruction loss

The VAF reconstruction loss is computed from the solution set produced by each clone-tree reconstruction method. This solution set Ω consists of three elements:

1. A set of trees {*t*_1_, *t*_2_, …, *t_U_*}
2. A probability distribution *p*(*t_u_*) over this set, with 0 ≤ *p*(*t_u_*) ≤ 1 and 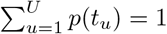
3. A set of subclonal frequencies 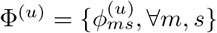 for each tree *t_u_*.

The loss is defined for each tree *t_u_* over the mutation read count data *x*, with mutations *m* and cancer samples *s*. We use 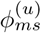 to indicate the subclonal frequency in *t_u_* for sample *s* associated with the subpopulation containing mutation *m*. For mutation *m* in sample *s*, we define the likelihood

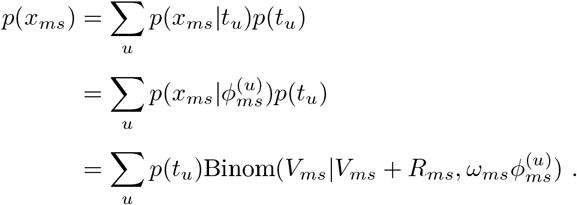

Now, to compute the VAF reconstruction loss *ϵ*_Ω_, we calculate the mean negative log-likelihood across all *M* mutations and *S* cancer samples, i.e.,

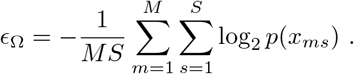

Note that because *p*(*x_ms_*) is a discrete distribution, *ϵ*_Ω_ ≥ 0.

We report VAF reconstruction loss relative to a baseline 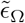. For simulated data, we use as the baseline a solution set Ω^base^ consisting of a single tree and the true subclonal frequencies Φ^true^ that generated the data. For real data, we use as the baseline the subclonal frequencies computed by Pairtree (Section 9.4) for our expert-derived trees. This yields the relative VAF loss 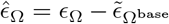. The relative VAF loss 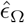 can be negative, indicating that a method has found a better solution than the baseline.

#### 5.6.1 Relationship reconstruction error

In determining relationship reconstruction error (Section 2.4), we wish to compare the distribution over pairwise mutation relationships imposed by a method’s set of candidate solutions relative to the simulated truth. Suppose a dataset consists of *M* mutations. Every clone tree built for this dataset by a method places each mutation pair (*i, j*) unambiguously into one of the four pairwise relationships. We use *θ_ij_* to delineate the pairwise model for the mutation pair induced by a given clone tree. Assume the method provides a distribution over different clone trees *t_u_*, with the posterior probability of *t_u_* represented as *p*(*t_u_*), such that ∑_*u*_ *p*(*t_u_*) = 1. In this case, we can compute the posterior probability of the *θ_ij_* relation as *p*(*θ_ij_*) = ∑_*u*_ *p*(*θ_ij_*|*t_u_*)*p*(*t_u_*), where

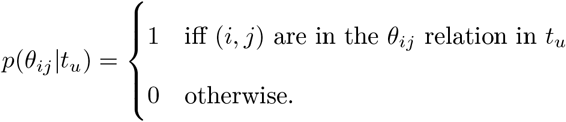

Using the set of true trees (Section 9.9.5), we will define 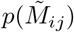 as the distribution over different relations for all *N* trees consistent with the true subclonal frequencies. For the true tree set, we will establish a uniform prior 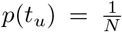, since no true tree should be privileged over another. For the mutation pair (*i, j*), we can now compute the Jensen-Shannon divergence (JSD) between a clone-tree-construction method’s *p*(*θ_ij_*) and the true 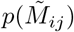, which we denote as 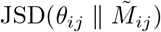. We use the base-two logarithm in computing JSD, yielding a measurement in bits.

Given *M* mutations in a dataset, there are 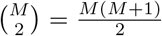 mutation pairs (*i, j*). We thus define the relationship reconstruction error *ϵ_R_* for a solution set as the mean JSD between pairs, such that

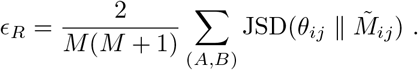

Using the mean allows us to compare *ϵ_R_* values for datasets with different numbers of mutations, so that we can understand which result sets have more or less error.

## 6 Acknowledgements

J.A.W. was supported by a Canada Graduate Scholarship from the National Sciences and Engineering Research Council of Canada, a Sir James Lougheed Award of Distinction from the Government of Alberta, and additional awards and funding from the University of Toronto Department of Computer Science and School of Graduate Studies, the Ontario Institute for Cancer Research, and the Vector Institute for Artificial Intelligence. Experiments were run using computational resources provided by SciNet and Compute Canada. The authors gratefully acknowledge Bei Jia and José Bento for extending their method for computing subclonal frequencies.

## 7 Author contributions

Q.D.M. conceived of and supervised the project. Q.D.M. and J.A.W. designed the project with input from S.M.D., and J.A.W. implemented Pairtree and ran the experiments. J.A.W. and Q.D.M. drafted the manuscript, and L.D.S. provided extensive edits and feedback. S.M.D. and J.E.D. designed the project and collected the data that motivated Pairtree’s development, and provided feedback throughout the project that guided the design of how Pairtree reports and visualizes its results. All authors reviewed and approved the final manuscript.

## 9 Supplementary information

### 9.1 Structure

The supplementary information is divided into two sections:

- Section 9.2 through Section 9.7 describe the methods composing Pairtree in greater detail and evaluate Pairtree’s ability to detect violations of the ISA.
- Section 9.8 through Section 9.17 examine characteristics of the simulated data and illustrate considerations that arose in benchmarking Pairtree relative to existing methods.

### 9.2 Computing pairwise relations

#### 9.2.1 Defining the data likelihood for pairwise relations

Let *A* and *B* represent two distinct mutations. We denote their observed read counts, encompassing both variant and reference reads, as *x_A_* and *x_B_*. Assuming both mutations obey the ISA, the pair (*A, B*) must fall in one of four pairwise relationships, denoted by *θ_AB_*.

1. *θ_AB_* = *coincident*, meaning *A* and *B* are co-occurring. That is, *A* and *B* occur in precisely the same cell subpopulations, implying *A* is never present without *B* and vice versa. This relationship indicates we cannot distinguish the order in which *A* and *B* were acquired because the data do not distinguish an intermediate subpopulation that occurred between them.
2. *θ_AB_* = *ancestor*, meaning *A* is ancestral to *B*. That is, *A* occurred in a population ancestral to *B*, such that some cells possess *A* without *B*, but no cell has *B* without *A*. This implies that *A* preceded *B*.
3. *θ_AB_* = *descendent*, meaning *B* is ancestral to *A*. This relationship implies *θ_BA_* = *ancestor*.
4. *θ_AB_* = *branched*, meaning *A* and *B* occurred on different branches of the clone tree, such that they never occur in the same set of cells. This relationship confers no information about the respective timing of *A* and *B*.

To the four possible relationships above, we add a fifth, termed the *garbage relation* and denoted by *θ_AB_* = *garbage*. This represents mutation pairs that do not fit into any of the four different relationships already defined, while also providing a baseline against which the other four relationships can be compared. This catch-all category places no constraints on the pairwise subclonal frequencies of the two mutations across cancer samples, so it is the only relationship that can include ISA violations identified by the four-gamete test [45]. This garbage relation could also model unreported CNAs that appear to be ISA violations, or highly inaccurate VAFs for one of the two mutations that arose from some artifact of the assay.

The likelihood of the pair’s relationship is written as *p*(*x_A_*, *x_B_*|*θ_AB_*). First, we note that every cancer sample *s* can be considered independently of others, i.e., they are exchangeable, so the likelihood factors as

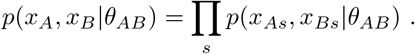

To compute the pairwise-relationship data likelihood for one cancer sample *s*, we integrate over the possible subclonal frequencies *ϕ_As_* and *ϕ_Bs_* associated with the subclones that gave rise to mutations *A* and *B*, respectively. This yields the likelihood

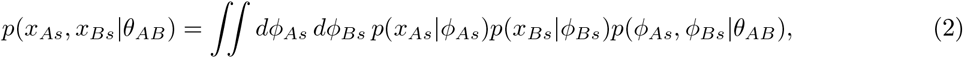

where the likelihoods of the observed read counts *x_As_* and *x_Bs_* are conditionally independent of all other variables given their corresponding subclonal frequencies *ϕ_As_* and *ϕ_Bs_*, respectively. In the following two section, we provide concrete definitions for each factor in Eq. (2).

#### 9.2.2 Observation model for read count data

For mutation *j* ∈ {*A, B*} from cancer sample *s*, whose observed read count data are represented by *x_js_*, we define *p*(*x_js_*|*ϕ_js_*) using the following variables:

- *V_js_*: number of genomic reads of *j*’s locus where the variant allele was observed
- *R_js_:* number of genomic reads of *j*’s locus where the reference allele was observed
- *ω_js_*: subclonal frequency to VAF conversation factor for *j*’s locus in sample *s*

Here *ω_js_* is usually used to correct for how, in autosomal regions of normal copy number in diploid cells, only half the alleles from *j*’s locus in a *j*-containing cell actually contain the variant *j*. In this case, *j*’s VAF is on average equal to half of its subclonal frequency, so 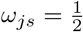. On sex chromosomes in males, which are haploid, *ω_js_* = 1.

In general, *ω_js_* = *Q_js_/W_js_* where *Q_js_* is the average number of *j*-variant-containing alleles present in a *j*-containing cell in sample *s*. Likewise, *W_js_* is the population-average copy number of the locus containing *j* (either the normal and variant allele) in sample *s*. By the ISA, in most cases *Q_js_* = 1, because the number of *j* variant alleles per cell can change only is if there is a copy number change (whether loss or gain) that affects the *j*-containing allele in cells within *j*’s subclone [40]. Unless there is strong evidence otherwise, we propose always setting *Q_js_* = 1. If there is evidence that *Q_js_* ≠ 1, unless *Q_js_* can be well-estimated in some other way, we propose not including that mutation in the input to Pairtree. Though Pairtree’s garbage detection may identify an uncorrected subclonal change in the number of mutant alleles per cell, not all such changes lead to ISA violations and so may not be detectable. In contrast, the value of *W_js_* should be set to account for any CNAs affecting any of the cells in sample *s* at *j*’s locus. Specifically, if there is a CNA at *j*’s locus that gives rise to a copy number *κ* in a fraction of cells equal to *ρ* in sample *s*, then *W_js_* = 2 + (*κ* – 2) * *ρ*, assuming that the normal copy number is 2. If this CNA is clonal, then *ρ* is the purity of the sample *s*. Often, the average copy number of a locus *W_js_* can be estimated directly from the relative read depth without solving for *κ* and *ρ*, in which case the direct estimate of *W_js_* can be used.

We use a binomial model Binom(*V_js_*|*N, p*) with parameters *N* and *p* to represent the likelihood of observing *V_js_* variant reads for mutation *j* in cancer sample *s*, given a subclonal frequency *ϕ_js_*. We set *N* = *V_js_* + *R_js_* to indicate the number of reads mapping to *j*’s genomic locus in sample *s*, and *p* = *ω_js_ϕ_js_* to represent the proportion of these reads that carry the variant. This yields *p*(*x_js_*|*ϕ_js_*) = Binom(*V_js_*|*V_js_* + *R_js_, ω_js_ϕ_js_*).

#### 9.2.3 Constraints on subclonal frequencies imposed by pairwise relationships

Now Eq. (2) only requires *p*(*ϕ_As_, ϕ_Bs_*|*θ_AB_*) to be defined. We use this prior to force *ϕ_As_* and *ϕ_Bs_* to be consistent with the relationship *θ_AB_*, as the ancestor, descendent, and branched relationships all place constraints on the subclonal frequencies *ϕ_As_* and *ϕ_Bs_*. We thus define

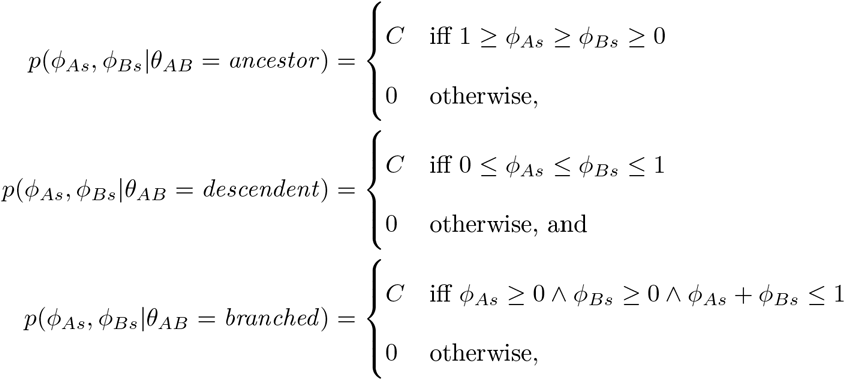

where *ϕ_As_, ϕ_Bs_* ∈ [0,1]. Note that for *θ_AB_* ∈ {*ancestor, descendent, branched*}, the prior *p*(*ϕ_As_, ϕ_Bs_*|*θ_AB_*) is non-zero only inside a right triangle lying within the unit square on the Cartesian plane with corners at {(0,0), (0,1), (1,0), (1,1)}. Specifically, each of the three different densities is non-zero only within a (different) triangle whose vertices correspond to three of the four unit square’s corners. As these triangles triangle have area 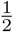, we set 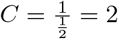 to ensure that all three prior densities integrate to 1.

We must still define the priors for the two remaining relationships, *θ_AB_* ∈ {*coincident, garbage*}. The garbage relationship permits all combinations of *ϕ_As_* and *ϕ_Bs_*, so we set

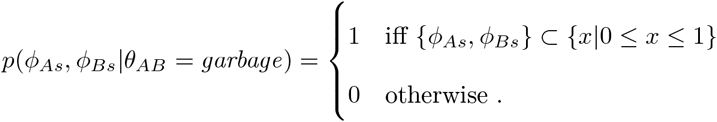

The coincident relationship requires the two mutations to arise from the same subclone, and so they are constrained to share the same subclonal frequency.

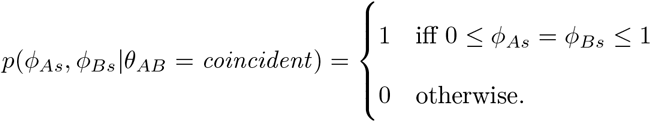

The garbage relationship establishes a baseline against which evidence for the non-garbage relationships can be evaluated. Observe that, in Eq. (2), *p*(*x_As_*|*ϕ_As_*)*p*(*x_Bs_*|*ϕ_Bs_*) is integrated over the unit square when *θ_AB_* = *garbage*. Conversely, when *θ_AB_* ∈ {*ancestor, descendent, branched*}, we integrate *p*(*x_As_*|*ϕ_As_*)*p*(*x_Bs_*|*ϕ_Bs_*) over a triangle covering half the square. Consequently, 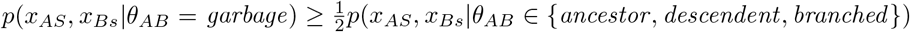. This arises because *p*(*ϕ_As_, ϕ_Bs_*|*θ_AB_*) = 2 for subclonal frequencies consistent with *θ_AB_* ∈ {*ancestor, descendent, branched*}, while *p*(*ϕ_As_, ϕ_Bs_*|*θ_AB_*) = 1 for subclonal frequencies consistent with *θ_AB_* = *garbage*. When the read counts for the mutations *A* and *B* clearly permit one of the three non-garbage relationships, most of the probability mass of the two associated binomials will reside within the simplex permitted by the relationship, and so contribution of the binomial likelihoods to the non-garbage relationship will be nearly the same as for the garbage relationship. Thus, the data likelihood (also known as evidence) for the non-garbage relationship will be nearly double that for the garbage one. Conversely, when the read counts push most of the binomial mass outside the permitted simplex, the non-garbage evidence will be substantially lower than the baseline provided by garbage. Although any one sample will always favour at least one non-garbage model over garbage, by accumulating evidence across many cancer samples, we can detect ISA violations. If different cancer samples favour different relationship types, the steady accumulation of the garbage evidence will outweigh the evidence for any of the other three relations, meaning garbage will be declared as the most likely relationship for the mutation pair. In Section 9.6.2, we describe how to use the pairwise evidence for garbage relationships to determine which mutations are the likely source of ISA violations.

#### 9.2.4 Efficiently computing relationship data likelihoods

We now consider how to compute the pairwise likelihood given in Eq. (2) for *θ_AB_* ∈ {*ancestor, descendent, branched*}.

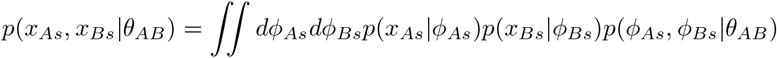

We can rearrange the integral to move the factor corresponding to the mutation *A* observations outside the inner integral.

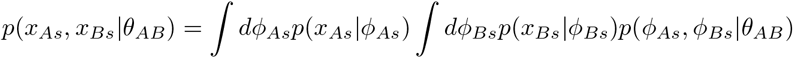

Now, because *p*(*ϕ_As_, ϕ_Bs_*|*θ_AB_*) is piecewise-constant when *θ_AB_* ∈ {*ancestor, descendent, branched*}, we can, for these relationships, impose this factor’s effect by changing the integration limits. Let *L*(*ϕ_As_, θ_AB_*) and *U*(*ϕ_As_, θ_AB_*)) represent functions whose outputs are the lower and upper integration limits, respectively, for the inner integral whose differential is *dϕ_Bs_*, as a function of *ϕ_As_* and the relationship *θ_AB_*. These functions are defined thusly:

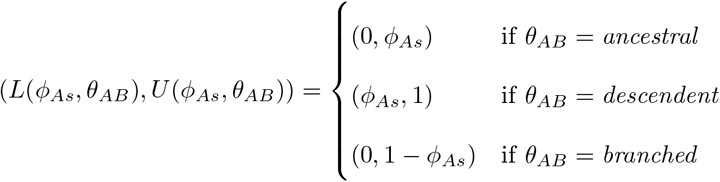

By writing the inner integral using these integration limits, and limiting the outer integral to the [0, 1] interval permitted for *ϕ_As_*, the factor *p*(*ϕ_As_, ϕ_Bs_*|*θ_AB_*) can be replaced by 2, as it is constant over the interval of integration.

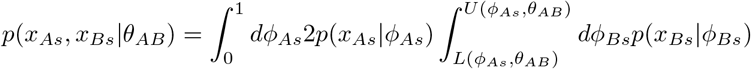

To render the inner integral more computationally convenient, rather than integrate over *ϕ_Bs_*, we would prefer to integrate over *q_Bs_* ≡ *ω_Bs_ϕ_Bs_.* Thus, we will integrate by substitution, using 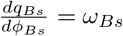.

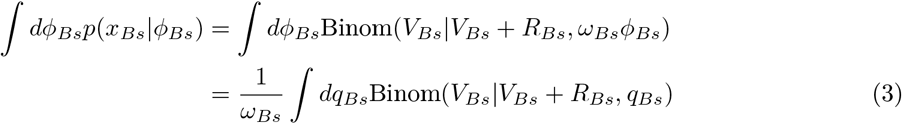

Observe that the inner integral is now simply integrating the binomial PMF over its parameter *q_Bs_.* To compute this integral, we rely on the following equivalence between this integral and the incomplete beta function *β*:

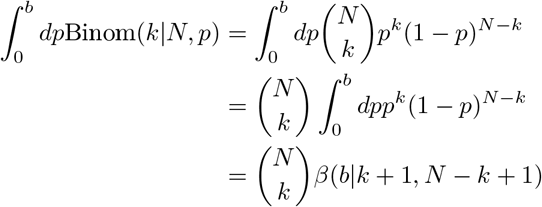

Now we can compute the integral over an arbitrary limit by the fundamental theorem of calculus.

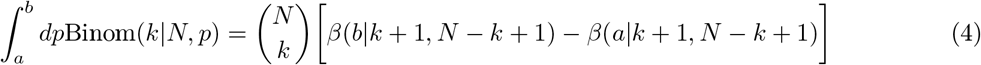

Finally, we combine the above results, allowing us to compute the pairwise relationship likelihood when *θ_AB_* ∈ {*ancestor, descendent, branched*} as a one-dimensional integral.

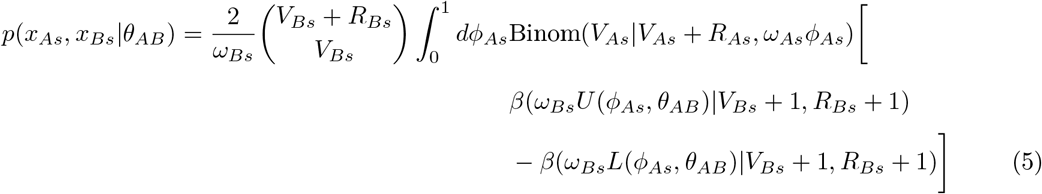

To compute this numerically, we use the one-dimensional quadrature algorithm from `scipy.integrate.quad`.

#### 9.2.5 Efficiently computing evidence for garbage and coincident pairwise relationships

We now examine how to compute the pairwise relationship likelihood for *θ_AB_* = *garbage* using the general likelihood given in Eq. (2). First, observe that we are integrating over *ϕ_As_* ∈ [0,1] and *ϕ_Bs_* ∈ [0,1], meaning there is no constraint placed on *ϕ_Bs_* by *ϕ_As_*. By removing the dependence of *ϕ_Bs_* on *ϕ_As_*, the likelihood can be broken into the product of two one-dimensional integrals, each taken over the interval [0,1]. Then, by drawing on results Eq. (3) and Eq. (4), we can compute an analytic solution to each integral.

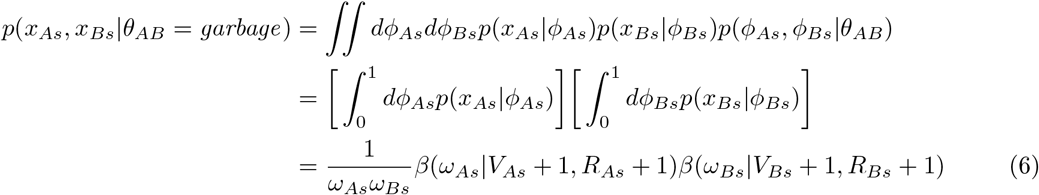

Finally, we compute the likelihood for *θ_AB_* = *coincident*. As our coincident constraint requires *ϕ_As_* = *ϕ_Bs_*, we are integrating along the diagonal line *ϕ_As_* = *ϕ_Bs_* that cuts through the unit square formed by *ϕ_As_* ∈ [0,1] and *ϕ_Bs_* ∈ [0,1]. This can be evaluated as a line integral along the curve *r*(*ϕ*) ≡ 〈*ϕ*, *ϕ*〉 for *ϕ* ∈ [0,1], with the Euclidean norm 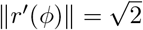.

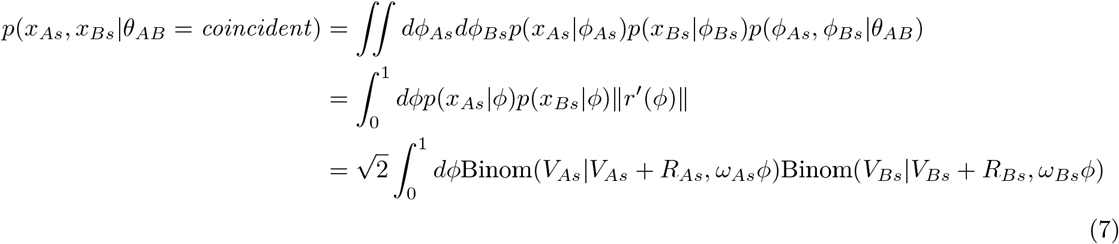

As with the ancestral, descendent, and branched relationships, we use the one-dimensional quadrature algorithm from `scipy.integrate.quad` to compute this.

#### 9.2.6 Computing the posterior probability for pairwise relationships

In Eq. (5), Eq. (6), and Eq. (7), we established how to compute the evidence for each of the five possible relations between mutation pairs, which takes the general form *p*(*x_A_, x_B_*|*θ_AB_*).. By combining these evidences with a prior probability *p*(*θ_AB_*) over relationships for mutation pair (*A, B*), we can compute the posterior probability *p*(*θ_AB_*|*x_A_, x_B_*) of each relationship.

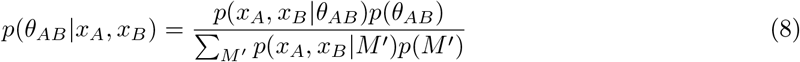

As we discuss in Section 9.3.8, we assume that, when Pairtree is run, mutations have already been clustered into subpopulations and “garbage” mutations have already been discarded. Consequently, we are computing pairwise relations between groups of mutations comprising subclones, and so we assign zero prior mass to the *coincident* and *garbage* relationships, ensuring these relationships also have zero posterior mass. The other three relationships are assigned the same prior probability, as we have no reason to believe one is more likely than the others.

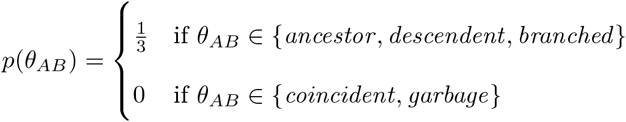

### 9.3 Performing tree search

#### 9.3.1 Representing cancer evolutionary histories with trees

Most clone tree reconstruction algorithms group mutations into subclones, with mutations that share the same subclonal frequency across cancer samples placed together. While thousands of mutations are typically observed using whole-genome sequencing, the mutations can typically be grouped into a much smaller number of subclones, simplifying the cancer’s evolutionary history. This grouping is valid because, as a cell population expands within a cancer, the frequencies of all mutations shared by cells in that population will increase in lockstep. Although Pairtree does not explicitly require that mutations be grouped into subclones, it can take these groupings as input. In this case, it replaces each mutation group with a single mutation, termed a *super-variant*, that represents the subclone.

When provided with *K* mutation clusters as input, each consisting of one or more mutations, Pairtree will produce a distribution over trees with *K* + 1 nodes. Node 0 corresponds to the non-cancerous cell lineage that gave rise to the cancer, while node *k* ∈ {1, 2, …, *K*} corresponds to the subclone associated with mutation cluster *k*. Node 0 always serves as the tree root, representing that the patient’s cancer developed from non-cancerous cells, and thus has no assigned mutations and a subclonal frequency of *ϕ*_0*s*_ = 1 in every cancer sample *s*.

An edge from node *A* to node *B* indicates that subclone *B* evolved from subclone *A*, acquiring the mutations associated with cluster *B* while also inheriting all mutations present in *A* and *A*’s ancestral nodes. The children of node 0 are termed the *clonal cancer populations.* Typically, there is only one clonal cancer population, but the algorithm allows multiple such populations when the data imply them. Multiple clonal cancer populations indicate that multiple cancers developed independently in the patient, such that they shared no common cancerous ancestor.

An edge from node *A* to node *B* means that, at the resolution permitted by the data, we cannot discern any intermediate cell subpopulations that occurred between these two evolutionary points. Nevertheless, such subpopulations may well have existed in the cancer.

#### 9.3.2 Tree likelihood

To describe the tree likelihood, we develop the following notation:

- *K*: number of cancerous subpopulations (and mutation clusters), with individual populations indexed as *k* ∈ {1, 2, …, *K*}
- *S*: number of cancer samples, with individual samples indexed as *s* ∈ {1, 2, …, *S*}
- *M_k_*: set of mutations associated with subclone *k*
- *V_ms_*: observed variant read count for mutation *m* in cancer sample *s*
- *R_ms_*: observed reference read count for mutation *m* in cancer sample *s*
- *ω_ms_:* probability of observing a variant read at mutation *m*’s locus within a subclone possessing *m*, in cancer sample *s*
- *ϕ_ks_*: subclonal frequency of subclone *k* in cancer sample *s*
- Φ: set of *ϕ_ks_* values for all *K* and *S*

The data *x* consists of the set of all *V_ms_, R_ms_*, and *ω_ms_* mutation values, as well as the *M_k_* clustering of those mutations into subclones. Given the tree *t*, consisting of a tree structure and associated subclonal frequencies Φ = {*ϕ_ks_*}, Pairtree uses the likelihood *p*(*x*|*t*, Φ) to score the tree. We describe how to compute the subclonal frequencies in Section 9.4. Below we use *x_ks_* to represent all data in sample *s* for the mutations associated with subclone *k*, while *x_ms_* refers to the data for an individual mutation *m*.

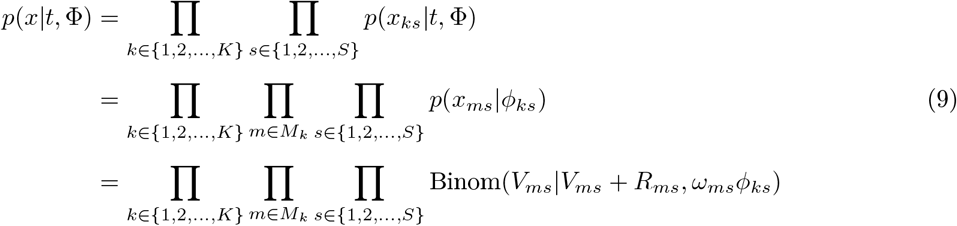

The likelihood Eq. (9) demonstrates that tree structure is not explicitly considered in the tree likelihood. Instead, we assess tree likelihood by how well the observed mutation data are fit by the tree-constrained subclonal frequencies accompanying the tree. Typically, we obtain a tree’s subclonal frequencies by making a maximum a posteriori (MAP) estimate, as described in Section 9.4.

Though Eq. (9) is ultimately the likelihood used by Pairtree for tree search, examining another perspective can help us understand what this likelihood represents. If we wished to directly assess the quality of a tree structure independent of its subclonal frequencies, thereby obtaining the likelihood *p*(*x*|*t*) rather than *p*(*x*|*t*, Φ), we would integrate over the range of tree-constrained subclonal frequencies permitted by the tree structure.

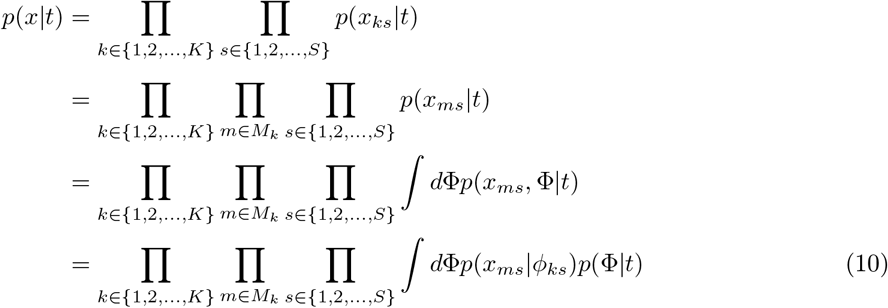

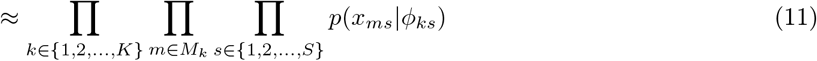

In Eq. (10), the factor *p*(Φ|*t*) is an indicator function representing whether the set of subclonal frequencies Φ obeys the constraints imposed by the tree structure *t*:

1. All subclonal frequencies exist within the unit interval, such that *ϕ_ks_* ∈ [0,1] for all *k* and *s.*
2. The non-cancerous node 0 is an ancestor of all subpopulations, such that *ϕ_ks_* = 1 for all *k* and *s*.
3. Let *C*(*k*) represent the children of population *k* in the tree. The subclonal frequency for *k* must be at least as great as the sum of its childrens’ frequencies, such that *ϕ_ks_* ≥ ∑_*c*∈*C*(*k*)_ϕ_*cs*_.

To obtain Eq. (10), we assume that only a narrow range of subclonal frequencies are permitted by the tree structure, and so we can use the MAP subclonal frequencies to approximate the integral and obtain Eq. (11), which is the likelihood function that Pairtree uses, as per Eq. (9). Consequently, we use Pairtree’s likelihood *p*(*x*|*t*, Φ) of the tree *t* and subclonal frequencies Φ as an approximation of the marginal likelihood of the tree *p*(*x*|*t*).

As an aside, note that a set of subclonal frequencies Φ obeying the three constraints enumerated above may be consistent with multiple tree structures—i.e., we may have *p*(Φ|*t*) ≠ 0 for a fixed Φ and different tree structures *t*. This shows how ambiguity may exist: a tree’s subclonal frequencies may permit multiple possible tree structures, all of which would be assigned the same likelihood. Each cancer sample’s subclonal frequencies typically impose additional constraints on possible tree structures, reducing this ambiguity.

#### 9.3.3 Using Metropolis-Hastings to search for trees

Pairtree uses the Metropolis-Hastings algorithm [37], a Markov Chain Monte Carlo method, to search for trees that best fit the observed read count data *x*. For notational convenience, our references to a tree *t* should be understood to implicitly include a set of subclonal frequencies Φ that have been computed for t, such that the likelihood denoted *p*(*x*|*t*) actually represents the likelihood *p*(*x*|*t*, Φ) described in Section 9.3.2.

According to the Metropolis-Hastings algorithm, to obtain samples from the posterior distribution over trees *p*(*t*|*x*), we must modify an existing tree *t* to create a new proposal tree *t*′. The *t*′ tree is accepted or rejected as a valid sample from the posterior according to how its likelihood *p*(*x*|*t*′) compares to the existing tree’s *p*(*x*|*t*), as well as the probabilities *p*(*t* → *t*′) of transitioning from the *t* tree to the *t*′ tree, and *p*(*t*′ → *t*) of returning from *t*′ to *t*. By Metropolis-Hastings, we assume that, given enough samples generated in this manner, we are eventually obtaining samples from the posterior distribution over trees 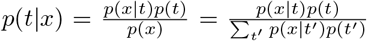. To establish our tree prior *p*(*t*), we denote the number of possible tree topologies for *K* subclones as *T*(*K*), which is a large but finite number that is exponential as a function of *K* [23]. Thus, we define our tree prior as a uniform distribution 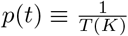, as we have no reason to prefer one tree structure to another a priori. Consequently, in computing the posterior ratio 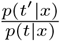 required for Metropolis-Hastings, all factors except the likelihoods *p*(*x*|*t*) and *p*(*x*|*t*′) cancel.

Pairtree can run multiple MCMC chains in parallel, with each starting from a different initialization (Section 9.3.7). By default, Pairtree runs a total of *C* chains, with *C* set to the number of CPU cores present on the system by default, and *P* = *C* executing in parallel. Both *P* and *C* can be customized by the user. From each chain, *S* = 3000 samples are drawn by default. The first *B* ∈ [0,1] proportion of trees are assumed to be early attempts by the sampling procedure to migrate toward high-probability regions of tree space, and so are discarded as burn-in samples because they are assumed to poorly reflect the true posterior. By default, we set 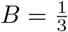. To reduce correlation between successive samples, Pairtree supports thinning, by which only a fraction *T* ∈ [0,1] of non-burn-in samples are retained. By default, Pairtree does not thin samples, so *T* = 1. Pairtree uses *T* to calculate a parameter 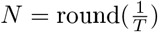, such that the algorithm records every *N*th sample. Thus, the actual number of trees recorded from a chain is 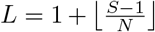. Only after thinning the chain are the burn-in samples discarded, resulting in round(BL) trees being returned as posterior samples from the chain. The *C, P, S, B*, and *T* parameters can all be changed by the user.

Once all chains finish sampling, Pairtree combines their results to provide an estimate of the posterior tree distribution. Given the uniform tree prior *p*(*t*), the posterior tree probability simplifies to 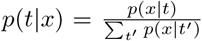. If the same tree *t* appears multiple times in this multiset—as it will, for instance, if proposal trees are rejected in Metropolis-Hastings and the last accepted tree is sampled again—each instance will appear as a separate term in the sum over *t*′, reflecting that each is a distinct sample from the posterior estimate.

#### 9.3.4 Modifying trees via tree proposals

To generate a new proposal tree *t*′ from an existing tree *t*, Pairtree relies on tree updates similar to those established in [18, 38]. The algorithm modifies *t* by moving an entire sub-tree under a new parent, or by swapping the position of two nodes. Specifically, Pairtree generates a pair (*A, B*), where *B* denotes a tree node to be moved, and *A* represents its destination. This pair is subject to the constraints {*A, B*} ⊂ {0,1, …, *K*}, such that *A* ≠ *B, A* is not the current parent of *B*, and *B* is not the root node 0. Two possible cases result. If *A* is a descendant of *B*, then the positions of *A* and *B* are swapped, without modifying any other tree nodes. This implies that the previous descendants of *B* (excluding *A* itself) become the descendants of *A*, while the previous descendants of *A* become the (only) descendants of *B*. Otherwise, *A* is not a descendant of *B* (i.e., *A* is an ancestor of *B*, or *A* is on a different tree branch), and so the sub-tree with *B* at its head is moved so that *A* becomes its parent. Observe that both moves can be reversed, which is a necessary condition for the Markov chain to satisfy detailed balance. In the first case, if *A* was descendent of *B* in *t*, then the pair (*B, A*) applied to the tree *t*′ will restore *t*. In the second case, if *A* was not descendent of *B* in *t*, and *b*’s parent in *t* was node *P*, then the pair (*P,B*) applied to tree *t*′ will restore *t*.

Pairtree provides two means of choosing the pair (*A, B*). The first mode uses the pairs tensor to inform tree proposals (Section 9.3.5). The second mode proposes tree updates blindly without reference to the data (Section 9.3.6), and is helpful for escaping pathologies associated with the first mode. Pairtree randomly selects between these modes for each update (Section 9.3.6).

#### 9.3.5 Using the pairs tensor to generate tree proposals

One of Pairtree’s key contributions is to recognize that the pairs tensor provides an effective guide for tree search, conferring insight into what portions of an existing tree suffer from the most error, and how those portions should be modified to reduce error. To create the proposal (*A, B*) for modifying the tree *t*, as described in Section 9.3.3, Pairtree generates discrete probability distributions *W^(A,B)^* and *W^(B)^*, corresponding to distributions over 0, 1, …, *K* that are used to sample *A* and *B*, respectively. The choice of *B* depends only on the current tree state *t*, and so we denote the corresponding probability distribution as *W*^(*B*)^. The choice of *A*, conversely, depends both on the current tree state *t* and whatever choice we made for *B*, and so we denote the corresponding probability distribution as *W*^(*A,B*)^. Every *W*^(*A,B*)^ and *W*^(*B*)^ depends solely on the tree state, such that the Markov chain used for Metropolis-Hastings is time-invariant.

The algorithm generates the probability distribution *W*^(*B*)^ such that the most probability mass is placed on elements corresponding to tree nodes with the highest pairwise error. First, observe that a tree induces a pairwise relationship between every pair of mutations—i.e., a tree places every mutation pair in a coincident, ancestral, descendent, or branched relationship. In Section 9.2, we described how to use mutation read counts to compute a probability distribution over these four relationships for every pair. For a given mutation *B*, we can thus compute the joint probability of the pairwise relationships between *B* and every other mutation induced by the tree *t* to determine how well-placed *B* is within the tree. Consider the mutation pair (k,B). If *p*(*θ_kB_*|*x_k_, x_B_*) represents the probability of the pair taking pairwise relation *θ_kB_*, then the probability of the pair taking one of the three other possible relationships is 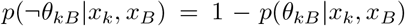, which we can think of as the pairwise relationship error. Then, the joint pairwise relationship error for all *K* – 1 pairs that include *B* is 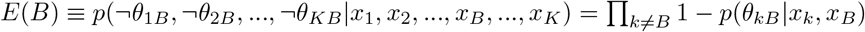.

We compute the probability distribution *W*^(*B*)^, whose elements represent the probability of selecting the node *B* to be moved within the tree, in accordance with the pairwise relationship error *E*(B). To accomplish this, we treat log *E*(*B*) as the logarithms of elements in an unnormalized probability distribution. To normalize the tuple (*E*(1), *E*(2), …, *E*(*K*)) to create a probability distribution, we use the scaled softmax function ssmax(*x*) ≡ softmax(*Sx*), where the *S* scalar is chosen such that 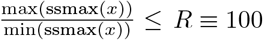. Specifically, the *S* scalar is set to 1 if 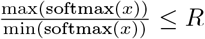, or otherwise to whatever value greater than 1 is necessary to make 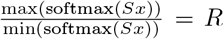. The scaled softmax can be understood as a “softer softmax,” ensuring no element in *W*^(*B*)^ ≡ ssmax((log *E*(1), log *E*(2), …, log *E*(*K*))) has more than 100 times the probability mass of any other. In practice, this results in every tree node having a non-trivial probability of being selected for modification.

With the probability distribution *W*^(*B*)^ established, we sample *B* ~ *W*^(*B*)^. We now need to establish how to compute the probability distribution *W*^(*A,B*)^, whose elements represent the probability of selecting the destination *A* for the node *B*. Critically, pairwise relations provide a computationally efficient means of evaluating hypothetical trees that modify *B*’s position—we can, in fact, test every possible proposal for *A* ∈ {0, 1, …, *K*} – {*B, P_B_*}, where *P_B_* denotes the current parent of *B*. With the choice of *B* already made, let *D_B_*(*A*) ≡ Π_(*j k*)_ *p*(*θ_jk_*|*x_j_, x_k_*) represent the joint probability of choosing *A* as the destination for *B*. By this formulation, (*j,k*) ranges over all 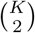 pairs within the set {1, 2, …, *K*}, and *D_B_*(*A*) represents the joint probability of all pairwise relations induced by the tree *t*^(*A,B*)^, which results from making the modification to tree *t* denoted by (*A, B*). Similar to *W*^(*B*)^, we apply the scaled softmax to the log *D_B_*(*A*) elements to create *W*^(*A,B*)^, with *W*^(*A,B*)^ ≡ ssmax((log *D_B_*(1), log *D_B_*(2), …, log *D_B_*(*K*))). We then sample *A* ~ *W*^(*A,B*)^.

We now have a concrete realization of the (*A, B*) pair that we can apply to tree *t*, yielding a modified tree *t*′. By using the pairwise relations as a guide, we selected a node (or subtree) *B* to modify, whose selection probability was dictated by the pairwise errors induced by its position in the tree. Then, we selected a destination *A*, which we swapped with the node *B* if *A* was already a descendant of *B*, or otherwise made the parent of the *B* subtree. In choosing *B*, we considered only the joint pairwise error of the *K* – 1 pairs including *B*; however, in choosing *A*, we considered the pairwise probabilities of all 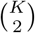 pairs that would result from the modified tree. Considering all pairs is necessary because moving the whole subtree rooted by *B* changed the position of all *b*’s descendants, potentially affecting many pairs that did not include *A* or *B*.

Thus, we selected a modification (*A, B*) to *t* that should, on average, yield a *t*′ tree with less error in pairwise relations. Ultimately, however, the question of whether to accept *t*′ as a posterior tree sample is decided by the Metropolis-Hastings decision rule that requires computing new subclonal frequencies Φ′ for *t*′, then considering the likelihood of the previous tree *p*(*x*|*t*, Φ) relative to the new likelihood *p*(*x*|*t*′, Φ′). Intuitively, once *B* is chosen, considering the change in pairwise relations induced by every possible choice of *A* captures substantial information about the quality of the tree that would be created by the (*A, B*) modification, while incurring only a modest computational cost. To fully evaluate the new tree *t*′, we must, however, use the full likelihood, which captures more subtle information about higher-order relations beyond pairwise. Though this is a more reliable indicator of the new tree’s quality, it requires the computationally expensive step of computing Φ′, which is why Pairtree does not do this when evaluating potential tree modification proposals.

#### 9.3.6 Escaping local maxima in tree space by allowing uniformly sampled tree proposals

Sampling the (*A, B*) tree modifications solely using the pairs tensor sometimes results in Pairtree becoming stuck in local maxima that exist in the tree space whose likelihood is defined with respect to the pairs tensor, but that have low likelihood in the tree space defined by the tree likelihood. Consequently, the tree-proposal algorithm may repeatedly propose tree modifications that improve consistency with pairwise relationships while worsening the overall tree, leading to many successive proposals being rejected. That is, some tree nodes may have high pairwise error, such that they are often sampled as the *B* subtree to modify. These nodes may in addition have destinations *A* within the tree that substantially reduce this pairwise error, resulting in the (*A, B*) modification being sampled with high probability. When the tree *t*′ induced by this modification is evaluated using the tree likelihood *p*(*x*|*t*′, Φ′), however, it may have poor likelihood, resulting in the modified tree being rejected by Metropolis-Hastings. This pathology occurs because *t*′ may appear to be a good candidate when only pairwise relations are considered, but when higher-degree relationships, such as those between mutation triplets, are captured in the subclonal frequency-based likelihood *p*(*x*|*t*′, Φ′), the tree is revealed to be poor.

Were the tree proposals (*A, B*) generated solely using the pairwise relations, Pairtree would repeatedly propose the same modification only to have it rejected, resulting in the algorithm becoming stuck at a sub-optimal point in tree space. To overcome this, we added two decision points in the tree generation process that permit uniformly sampled modifications. Firstly, when sampling the node *B* to move within the tree, Pairtree will use the pairwise relation-informed choice only *γ* = 70% of the time. In the other 1 – *γ* = 30% of cases, Pairtree will sample *B* from the discrete uniform distribution over {1, 2, …, *K*}. Secondly, in *ζ* = 70% of cases, Pairtree will sample the destination node *A* from the discrete uniform distribution over {0, 1, …, *K*} – {*B,P_B_*}, where *P_B_* denotes the current parent of *B*. Both decisions are made independently and at random when generating the tree proposal, such that a proposal using pairwise relations for both *A* and *B* is generated for only *γζ* = 49% of tree modifications. Conversely, (1 – *γ*)(1 – *ζ*) = 9% of tree modifications are generated without considering the pairwise relations in any capacity. Both *γ* and *ζ* can be modified at runtime by the user. Their default values were chosen to ensure that approximately half of tree modification proposals are fully informed by pairwise relations, while the remaining half ignore the pairwise relations for at least part of the proposal generation, allowing the algorithm to explore regions of tree space that might otherwise be rendered difficult to reach.

#### 9.3.7 Tree initialization

‘ To sample trees via Metropolis-Hastings, the MCMC chain must be initialized with a tree structure. Similar to the tree-sampling process, which can generate proposals using the pairs tensor (described in Section 9.3.5) or without it (Section 9.3.6), the initialization algorithm can use the pairs tensor to infer reasonable relationships between subclones, or can ignore the pairs tensor and thereby avoid potential biases that would inhibit tree search.

We first describe tree initialization using the pairs tensor. In this mode, Pairtree constructs the tree in a top-down fashion, selecting subclones to add to the tree with a sampling probability based on which appear to have the most ancestral relationships relative to subclones not yet added. Once the algorithm determines which subclone to add, it considers all possible parents from amongst the nodes already added, sampling a choice based on which induces the least pairwise relation error for all subclones. This algorithm uses the scaled softmax described in Section 9.3.5.

**Listing 1:**
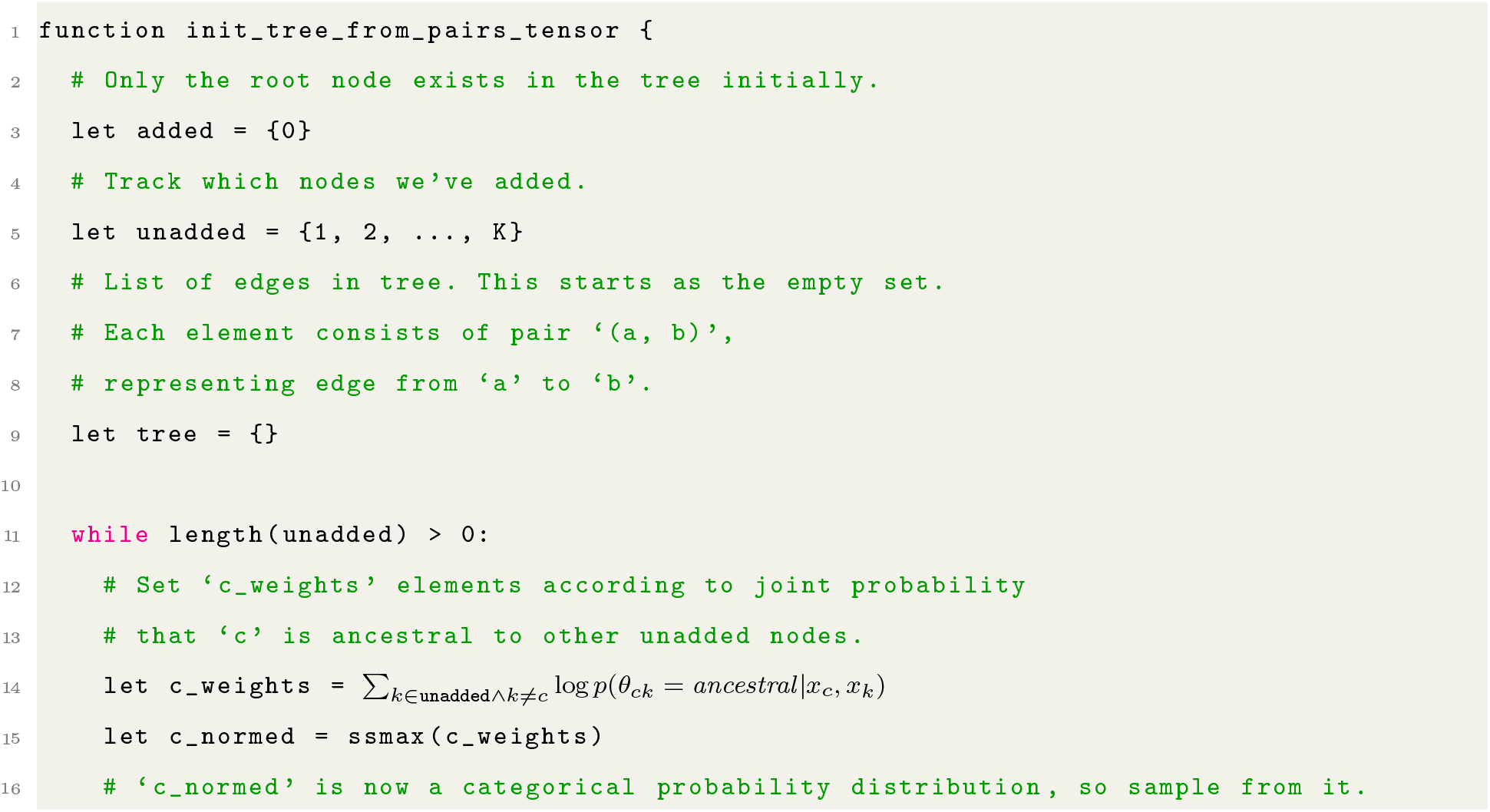
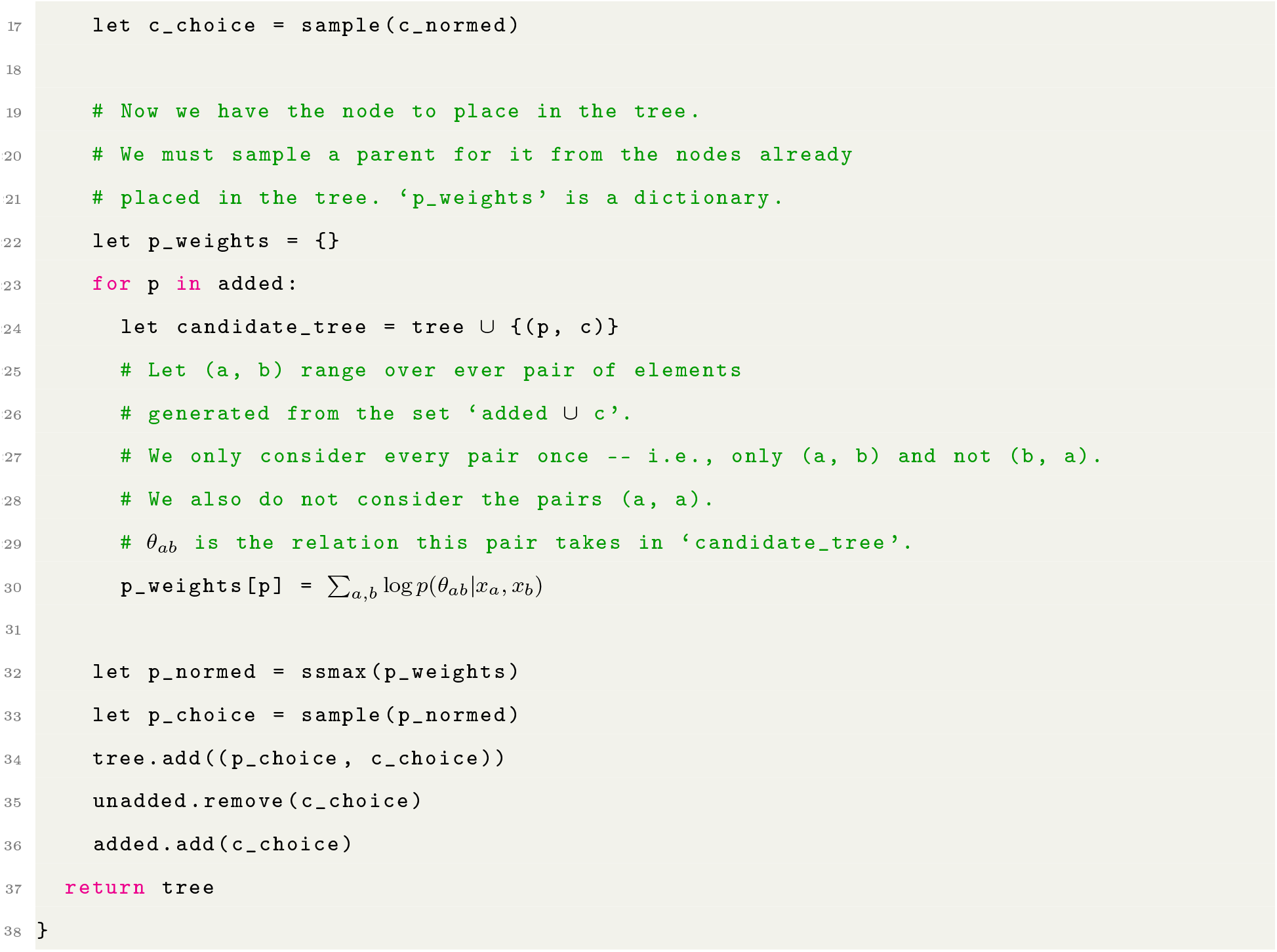
Tree initialization algorithm using pairs tensor.

In the second mode, Pairtree initializes a tree without reference to the pairwise relations, by placing every subclone as an immediate child of the root. This initialization is unbiased insofar as it imposes no ancestral or descendent relationships amongst subclones, assuming instead that the Metropolis-Hastings update scheme can rapidly alter this initial tree to produce a reasonable solution.

When initializing an MCMC chain, Pairtree selects between the two initialization modes at random, with probability ι = 70% of selecting the pairwise-relation-based mode, and 1 – *ι* = 30% probability of the unbiased mode. The *ι* parameter can be specified by the user, with the default value chosen under the assumption that Pairtree will typically be used in multi-chain mode, such that different chains will benefit from different initializations that allow the algorithm to more fully explore tree space.

#### 9.3.8 Reducing Pairtree’s computational burden using supervariants

Pairtree assumes that mutations have been clustered into subpopulations, with “garbage” variants discarded, before the tree-construction algorithm begins. As a result, all mutations within a subpopulation are rendered *coincident* relative to one another. Mutations within a subclone also share the same evolutionary relationships to all mutations outside the subclone. Thus, to reduce the computational burden imposed by the method, rather than working with individual mutations, we can instead represent each subpopulation with a single *supervariant*, then compute pairwise relations between these rather than their constituent mutations.

Conceptually, relative to the individual mutations that compose it, a supervariant should provide a more precise estimate of the subclonal frequency of its corresponding subclone. Specifically, a mutation *m* in a cancer sample *s* has *V_ms_* variant reads and *R_ms_* reference reads, yielding total reads *T_ms_ ≡ V_ms_* + *R_ms_* and a 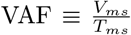. Given a probability of observing the variant allele *ω_ms_*, we conclude that *ω_ms_T_ms_* reads originated from the variant allele, and so we can estimate the corresponding subclone’s subclonal frequency by 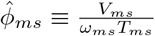. Each mutation’s 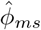 should thus serve as a noisy estimate of its subclone’s true *ϕ_m_s*.

Let *x_ms_* represent the data associated with mutation *m* in sample *s*, such that *x_ms_* ≡ {*V_mS_, R_ms_, ω_ms_*}. Under a binomial observation model (Section 9.3.2), given subclonal frequency *ϕ_ks_* for the subclone *k* harboring mutation *m* in sample *s*, we have the mutation likelihood *p*(*x_ms_*|*ϕ_ks_*) ≡ Binom(*V_ms_*|*V_ms_* + *R_ms_, ω_ms_ ϕ_ks_*). Let *M_k_* be the set of mutations associated with subclone *k*. Then, from all *j* ∈ *M_k_*, we get the following joint likelihood for cancer sample *s*:

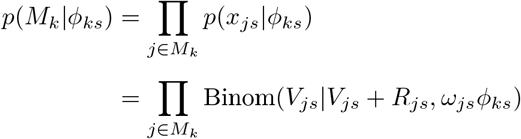

Assuming *ω_js_* takes the same value *ω_ks_* for all *j* ∈ *M_k_*, the joint likelihood takes the following form:

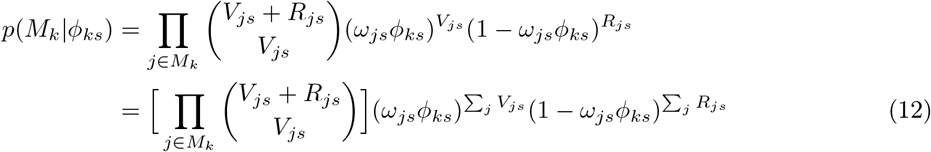

We want the likelihood for the supervariant *k* representing the variants in *M_k_* to take the same functional form. Thus, we set *V_ks_* ≡ ∑_*j*∈*M_k_*_ *V_js_* and *R_ks_ ≡ ∑_*j*∈M_k__ R_js_*, yielding the following supervariant likelihood.

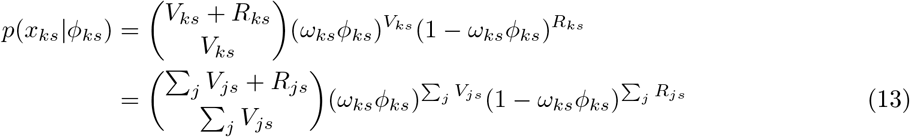

Observe that Eq. (13) takes the same functional form as Eq. (12), such that they differ only by a constant of proportionality *C* that does not depend on *ϕ_ks_*.

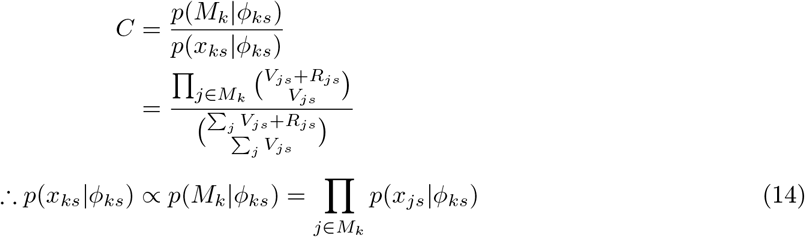

Consequently, the supervariant′s likelihood accurately reflects the joint likelihood of the subclone’s constituent variants, while reducing the algorithm’s computational burden. In practice, the constant factor *C* by which the two differ does not matter, as the Metropolis-Hastings scheme (Section 9.3.3) that uses the likelihood (Section 9.3.2) requires only the ratio of two likelihoods to navigate tree space, such that *C* cancels.

Of course, Eq. (14) holds only when *ω_ks_* = *ω_js_* for all *j* ∈ *M_k_*. Most often, we are given diploid variants with 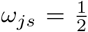, and so we fix 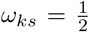 for all supervariants. Thus, supervariants are assured to accurately represent their constituent variants when those variants are from diploid genomic regions. For non-diploid variants with 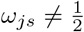, we must rescale the provided data *x_js_* to use a fixed 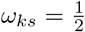, allowing us to use an approximation of the correct likelihood. To achieve this, we establish the following:

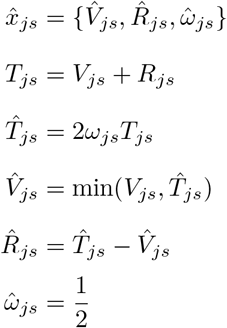

This representation ensures the corrected variant read count 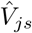 cannot exceed the corrected total read count 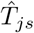, which could otherwise occur because of binomial sampling noise inherent to the genomic sequencing process, or an erroneous *ω_js_* value that does not correctly reflect a copy number change. Note that both 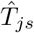 and 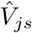 can take non-integer values. If the original 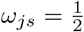, then the corrected read counts are unchanged from their original values. From this point, for all mutations *j* ∈ *M_k_* associated with subclone *k*, we compute corrected supervariant read counts 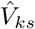 and 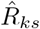:

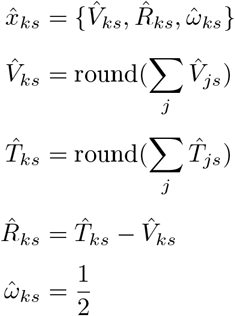

Based on Eq. (14), if the mutations *j* ∈ *M_k_* all had 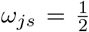, the *ϕ_ks_* value we obtain in maximizing the supervariant likelihood 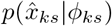 is also optimal for the full joint likelihood over the individual mutations *p*(*M_k_*|*ϕ_ks_*) = Π_*j*∈*M*_*k*__ *p*(*x_js_*|*ϕ_ks_*), since the two likelihoods differ only by a constant of proportionality. If some mutations *j* had 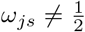, the supervariant likelihood 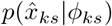 approximates the full joint likelihood, and so the obtained *ϕ_ks_* value is only approximately optimal for the latter. To overcome this, Pairtree’s implementation of the rprop optimization algorithm could be easily modified to optimize *ϕ_ks_* with respect to the individual variants *j*, each with its own *ω_js_*, rather than the combined supervariant representation that requires a single *ω_ks_*. Equivalently, rprop could use multiple supervariants per subclone, with a single supervariant representing all constituent mutations possessing the same *ω_js_*. The projection algorithm, however, necessitates using a single supervariant, which in turn requires a single *ω_ks_*. Though the adjusted supervariant read counts yield only an approximation of the likelihood for non-diploid mutations, this is not a critical flaw, as projection is already computing a Gaussian approximation of the likelihood, rather than the exact binomial likelihood used by rprop.

### 9.4 Fitting subclonal frequencies to trees

Pairtree provides two algorithms for computing subclonal frequencies for a tree structure. Both attempt to maximize the data likelihood (Section 9.3.2), fitting the observed read count data as well as possible while fulfilling all constraints imposed by the tree structure. The first algorithm, named *rprop*, is based on gradient descent (Section 9.4.2), and directly maximizes the tree’s binomial likelihood. The second algorithm, named *projection*, uses techniques from convex optimization to compute subclonal frequencies maximizing the likelihood of a Gaussian approximation to the binomial [41]. While rprop typically produces higher-likelihood subclonal frequencies than projection, particularly for subclones where the Gaussian approximation to the binomial is poor, the projection algorithm runs substantially faster with many subclones (e.g., for 30 subclones or more). By default, Pairtree uses the projection algorithm, but the user can select rprop at runtime.

#### 9.4.1 Converting between subclonal frequencies and subpopulation frequencies

To permit a more convenient representation, both rprop and projection work with subpopulation frequencies *H* = {*η_ks_*}, rather than the subclonal frequencies Φ = {*ϕ_ks_*}, where *k* and *s* are indices over subclones and cancer samples, respectively. Given a tree structure *t*, we can readily convert from one representation to the other. Let *D*(*k*) represent the set of descendants of subclone *k* in tree structure *t*, and *C*(*k*) represent the set of direct children of subclone *k*. Then, in cancer sample *s*, we have

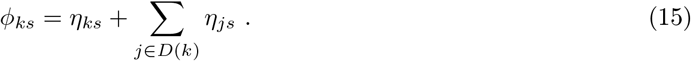

Equivalently, we obtain

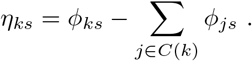

From the subclonal frequency constraints described in Section 9.3.2, we see that, because the root node takes *ϕ*_0*s*_ = 1, we must have the constraint

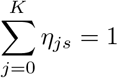

across all *K* subclones, and that each individual *η_js_* ∈ [0,1]. As each cancer sample *s* is independent from every other, both rprop and projection optimize the set {*η_ks_*} separately for each fixed *s*.

#### 9.4.2 Fitting subclonal frequencies using rprop

The rprop algorithm is a simpler version of RMSprop [46, 47], intended for use with full data batches rather than mini-batches. To perform unconstrained optimization on the parameters *H_s_* = {*η_ks_*} for a fixed cancer sample *s*, the algorithm first reparameterizes to *H_s_* = softmax({*ψ_ks_*}), so that we need not enforce constraints on {*ψ_ks_*} but can ensure *H_s_* ⊂ [0,1] and ∑_*k*_ *η_ks_* = 1.

On each iteration, given a tree structure *t* and existing subclonal frequencies Φ, rprop converts Φ to population frequencies *H*, then computes the derivatives

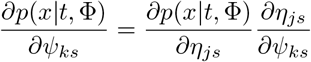

for all subclone combinations *j* and *k*, using the tree likelihood (Section 9.3.2). The algorithm then uses the sign of the gradient to update the *ψ_ks_* values, ignoring the gradient′s magnitude. For each value of *k*, rprop maintains a step-size parameter *λ_k_*, which is limited to lie within the interval [10^-6^,50], preventing excessively small or large step sizes. The algorithm also maintains a step-size multiplier *P_ki_* for subclone *k* on iteration *i*, with *P_ki_* = 1.2 if 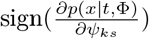 agrees with the sign from the previous iteration *i* – 1, and 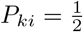 otherwise. Using these values, rprop performs the gradient update

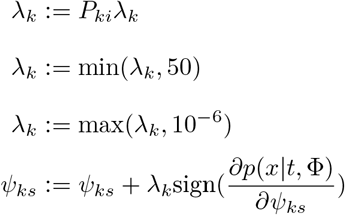

The rprop algorithm continues this process until none of the 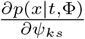 values exceed 10^-5^ in a particular iteration, or until *I* = 10000 iterations elapse, with *I* being customizable by the user.

To initialize the *H_s_* = {*η_ks_*} values, we generate initial values 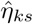 with the following algorithm. *C*(*k*) represents the set of direct children of *k* in the tree.

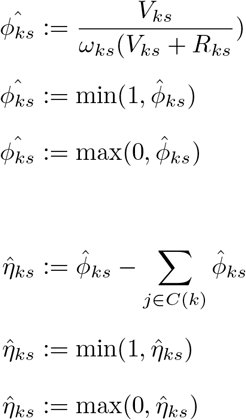

Observe that the constraint 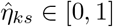 is satisfied. To ensure 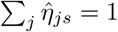, we finally set 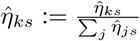. This initialization reflects that, if the provided tree structure *t* is consistent with the data and there is minimal noise in the data, the 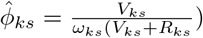 subclonal frequencies should be close to the maximum likelihood estimate for Φ in *p*(*x*|*t*, Φ).

#### 9.4.3 Proof that the subclonal frequency objective is convex

From Eq. (9), we have

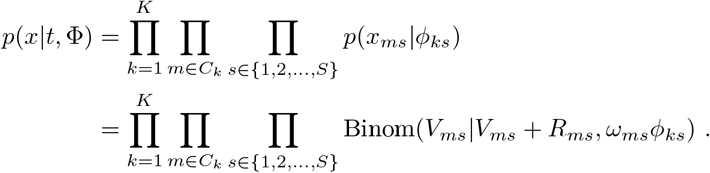

Since we fit the subclonal frequencies Φ^(*s*)^ for each cancer sample *s* independently, it suffices to consider

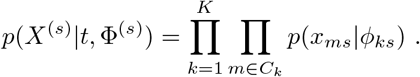

We will examine

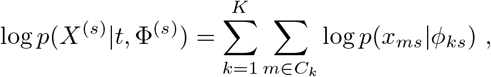

which is the objective optimized by the rprop algorithm (Section 9.4.2).

By Eq. (15), we have *ϕ_ks_* = *η_ks_* + ∑_*j*∈*δ_k_*_ *η_js_*, where *δ_k_* represent the set of descendants of subclone *k* within a given tree. Thus

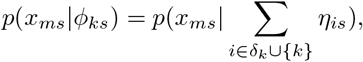

which implies

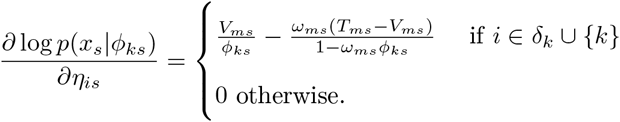

Note that the partial derivative with respect to *η_is_* can be written purely with reference to *ϕ_ks_*. This yields

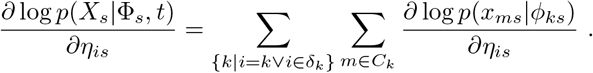

Next, we will define the Hessian via the second-order derivatives

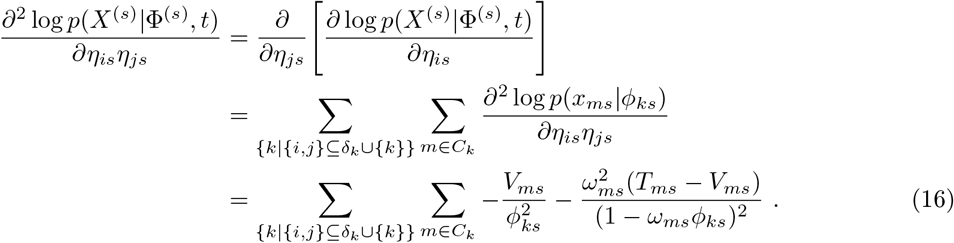

Let **A**^(s)^ correspond to the Hessian for the negative log-likelihood for sample *s*, such that

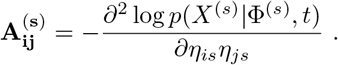

Observe that

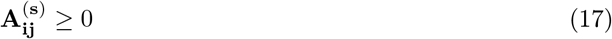

because, ∀*k*∀*m*∀*s*, we have *ϕ_ks_* ∈ (0,1], *ω_ms_* ∈ (0,1], *T_ms_* ∈ {1, 2,…}, and *V_ms_* ∈ {0,1,…}. Using similar logic, we have

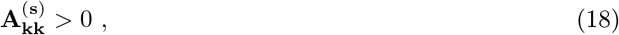

since *k* ∈ *δ_k_* ∪ {*k*} and every subclone *k* will have one or more mutations *m* ∈ *C_k_* that are associated with it.

Now we will demonstrate that **A**^(s)^ is symmetric positive-definite. By definition, the Hessian is symmetric. We can express the Hessian as a matrix decomposition with matrices *B* and *D*^(*s*)^ such that **A**^(s)^ = *BD*^(*s*)^ *B*^*T*^. Let *D*^(*s*)^ be the *K* × *K* diagonal matrix where

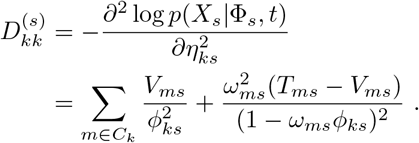

Let *B* be the *K* × *K* ancestral matrix such that

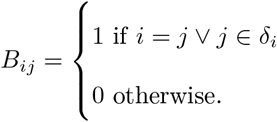

These matrices define the decomposition **A**^(s)^ = *BD*^(*s*)^*B^T^*. Without loss of generality, we can index the clone tree nodes according to a topological sort, thereby ensuring that *B* will be upper (or lower) triangular. Under these conditions, as *B*’s diagonal consists entirely of 1, we have 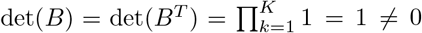, and so *B* and *B^T^* are both invertible. Since (*B^T^*)^*T*^ *D*^(*s*)*T*^ *B*^*T*^ = *BD*^(*s*)^ *B^T^* = **A**^(s)^, we have demonstrated that **A**^(s)^ is congruent with the diagonal matrix *D*^(*s*)^, and thus **A**^(s)^ is positive-definite. Because the Hessian is positive-definite for the entire domain of the negative log-likelihood – log *p*(*X*^(*s*)^|Φ^(*s*)^,*t*), the function is convex.

Finally, we will show that the likelihood’s domain is convex. We can consider the domain of the negative log-likelihood to consist of valid subpopulation frequency vectors *H*^(*s*)^ = [*η*_0*s*_, *η*_1*s*_, *η*_2*s*_,… *η_Ks_*], where each element must be non-negative and together they must sum to unity. This is valid because such a vector *H*^(*s*)^ in conjunction with the tree structure *t* uniquely determine the subclonal frequencies Φ^(*s*)^ for use in – log *p*(*X*^(*s*)^|Φ^(*s*)^,*t*). If we take a scalar 0 ≤ *j* ≤ 1 and any two subpopulation frequency vectors 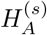 and 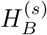, then 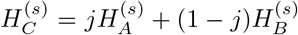 will also lie within the domain, as all elements of 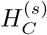 will also be non-negative and sum to unity. Consequently, as a convex combination of any two members of the domain is itself convex, the domain is convex. Thus, as both the function domain and the function being optimized are convex, the problem as a whole is convex, and rprop should converge on a globally optimal solution.

#### 9.4.4 The posterior mass for subclonal frequencies is highly concentrated around the MAP estimate

When computing the subclonal frequencies Φ using the read-count data *x* and a tree structure *t*, we first obtain the subpopulation frequencies *H*, which we then transform to Φ using *t* (Section 9.4.1). Conceptually, we wish to understand our uncertainty in *H* stemming from the distribution *p*(*H*|*x, t*). As we optimize each cancer sample *s* independently, we only need to characterize *p*(*H*^(*s*)^|*X*^(*s*)^, *t*), yielding the subpopulation frequency vector *H*^(*s*)^ for sample *s* using the corresponding data *X*^(*s*)^ and tree structure *t*. Properties of this distribution imply that we can use the MAP estimate of this distribution as a good representation for it because minimal uncertainty exists.

As we previously demonstrated, the distribution itself and the overall optimization problem are convex (Section 9.4.3). Now, note that

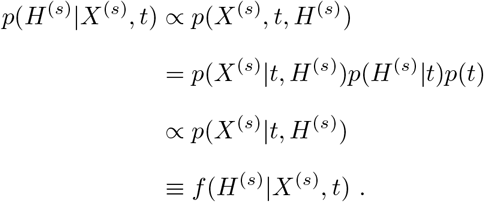

We do not consider *p*(*H*^(*s*)^|*t*), as the distribution over subpopulation frequencies is a uniform Dirichlet given the tree structure. Specifically, *p*(*H*^(*s*)^|*t*) = *K*! for a tree containing *K* cancerous subclones and thus *K* + 1 nodes. Likewise, we use a uniform prior over trees *p*(*t*). As in Section 9.4.3, we will define **A**^(s)^ as the Hessian for the negative log-likelihood in sample *s*, such that

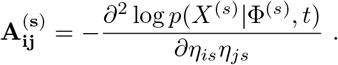

We will use Laplace’s method to approximate *f*(*H*^(*s*)^ *X*^(*s*)^,*t*) = *p*(*X*^(*s*)^|*H*^(*s*)^,*t*), which we have shown is proportional to the desired distribution *p*(*H*^(*s*)^ *X*^(*s*)^,*t*). Using the Hessian **A**^(s)^, we have

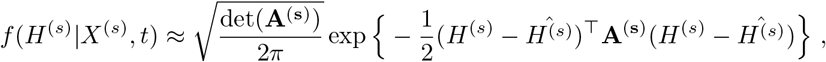

where 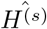 indicates the MAP estimate for our subpopulation frequency vector. In the Gaussian approximation produced by Laplace’s method, **A**^(s)^ serves as the precision matrix. By Eq. (16), we have

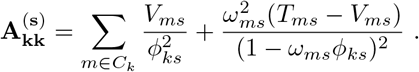

Observe that the diagonal elements of *A*^(*s*)^ will all have magnitude

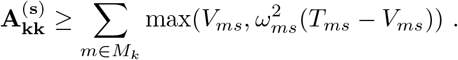

This occurs because *ϕ_ks_* ∈ (0,1], *ω_ms_* ∈ (0,1], *T* ∈ {1, 2,…}, and *V* ∈ {0,1,…}. At even moderate read depths, the variance of each dimension of our Gaussian approximation, given by 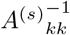, will be extremely low—by summing the read counts, whether in low-depth settings with relatively many variants or high-depth settings with relatively few variants, we obtain a lower bound on precision that reveals a Gaussian with extremely low variance. For instance, consider the case of just one mutation *m* at a typical read depth of *T_ms_* = 50 in a diploid setting with 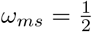. If this is the only variant associated with subclone *k*, then regardless of the variant read count *V_ms_*, we have 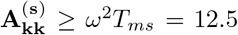. Most subclones will, of course, have higher read depths and/or more assigned mutations, driving this bound much higher. Consequently, most probability mass will thus be concentrated around the MAP estimate, meaning the MAP can serve as a reasonable proxy for the distribution as a whole.

Finally, note that the *Z*(Φ^(MAP)^) factor used in the Laplace approximation to the marginal likelihood provided by Eq. (11) depends on det(**A**^(s)^). Because **A**^(s)^ = *BD*^(*s*)^*B^T^* and det(*B*) = det(*B^T^*) = 1, we have

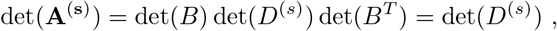

which is independent of the tree structure represented by *B*.

#### 9.4.5 Fitting subclonal frequencies using projection

The projection algorithm draws on the approach provided in [41]. The authors describe a method to efficiently enumerate mutation trees, in which individual nodes correspond to genomic mutations. To make this enumeration feasible, they developed an algorithm to rapidly compute tree-constrained subclonal frequencies. Using our supervariant representation, we can apply their approach to computing subclonal frequencies for clone trees by representing our binomial likelihood with a Gaussian approximation. First, we review the authors’ notation and map it to the equivalent notation in Pairtree.

- : *q*: number of mutations, equivalent to our number of subclones *K*
- : *p*: number of cancer samples, equivalent to our *S*
- : *F* ∈ ℝ^*q*×*p*^: equivalent to our subclonal frequencies Φ, with *F_vs_* equivalent to our *ϕ_ks_*
- : *U* ∈ {0,1}^*q*×*q*^: ancestry matrix created from tree structure *t*, such that *U_j,k_* = 1 iff subclone *j* is an ancestor of subclone *k* or *j = k*
- *M* ∈ ℝ^*q*×*p*^: equivalent to our population frequencies *H* = {*η_ks_*}, with *M_vs_* equivalent to our *η_ks_*

With 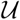 representing the set of all ancestral matrices consistent with the perfect phylogeny problem (Section 9.6.1), the authors solve the optimization problem 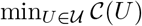, such that

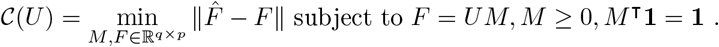

Here, || · || is the Frobenius norm, and 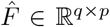 is the noisy estimate of the subclonal frequencies obtained from the data. Observe there is a one-to-one correspondence between *U* and *t*, as changing the structure of *t* will necessarily change ancestral relations described in *U*, and vice versa. Thus, the authors attempt to find the optimal ancestry matrix *U*, corresponding to an optimal tree *t*, that allows tree-constrained subclonal frequencies *F* best matching the noisy subclonal frequencies 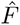 observed in the data. Ultimately, the authors solve this problem through enumeration. While this scales better than previous enumerative approaches because of the authors’ efficient computation of the optimal *M* for a given ancestry matrix *U*, the approach is still rendered infeasible for the large trees that Pairtree works with using a search-based method.

Useful for Pairtree is the authors’ extremely efficient means of projecting the observed frequencies 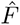 on to the space of valid perfect-phylogeny models using Moreau’s decomposition for proximal operators and a tree reduction scheme [41]. We utilize this to quickly compute subclonal frequencies Φ for a given tree *t* that corresponds to an ancestry matrix *U*. To allow us to use a Gaussian estimate of our binomial likelihood, the authors developed an extended version of their algorithm [48], in which they additionally take as input a scaling matrix *D* ∈ ℝ^*q*×*p*^ with all *D_ks_* > 0. Using the element-wise multiplication operator ⊙, the modified algorithm computes

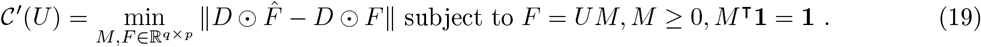

We will refer to the algorithm as the “projection optimization algorithm,” and to Eq. (19) as the “projection objective.” We now show how to use the projection objective to compute the MAP for a Gaussian approximation of our original binomial likelihood. First, observe that our goal is to maximize the binomial likelihood defined in Section 9.3.2 by finding optimal subclonal frequencies Φ for a given tree *t*. Thus, we wish to find

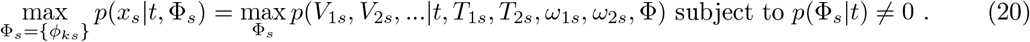

Here, *t* represents the provided tree structure, while Φ_*s*_ refers to a set of scalar *ϕ_ks_* values that obey the tree constraints described in Section 9.3.2, with *p*(Φ_*s*_|*t*) ≠ 0 indicating that the set obeys the constraints. The *s* index represents the cancer sample, with each sample optimized independently. Our data *x_s_* consists of, for subclone *k*, a count of variant reads *V_ks_* and reference reads *R_ks_*, yielding total reads *T_ks_* = *V_ks_* + *R_ks_*. We define this as a binomial likelihood, in which we are optimizing the *ϕ_ks_* values.

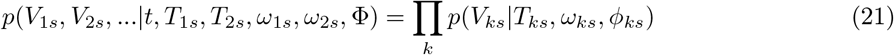

To approximate this using the Gaussian, we perform the following operations.

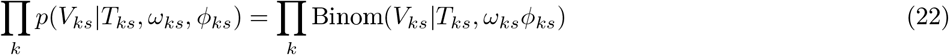

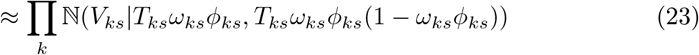

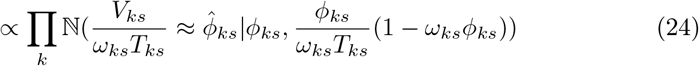

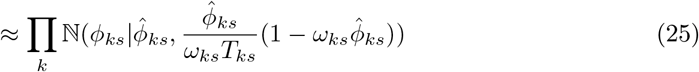

We relied on the following operations to achieve the above:

- Eq. (22) defined Eq. (21) with respect to the binomial distribution.
- Eq. (23) approximated Eq. (22) with the Gaussian distribution. We represent the Gaussian PDF for a random variable *x* drawn from a Gaussian with mean *μ* and variance *σ*^2^ as 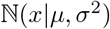.
- Eq. (24) divided the Gaussian random variable by the scalar *ω_ks_T_ks_*, yielding another Gaussian proportional to the preceding. The new Gaussian random variable is 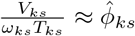, our MAP of the subclonal frequency *ϕ_ks_* for Binom(*V_ks_*|*T_ks_*, *ω_ks_ ϕ_ks_*). As *ϕ_ks_* ∈ [0,1], we set 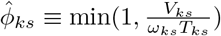.
- To achieve a distribution over the unknown *ϕ_ks_*, Eq. (25) swaps the Gaussian’s random variable 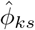 and mean *ϕ_ks_*, yielding the same Gaussian PDF. Additionally, it approximates the variance of the Gaussian in *Eq*. (24) by replacing *ϕ_ks_* with its MAP in the variance definition.

Let the variance of each Gaussian be represented with 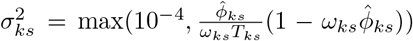. We set a minimum variance of 10^-4^ to prevent our *ϕ_ks_* estimates from being too precise to permit effective optimization. To transform Eq. (25) into the form required by the projection objective Eq. (19), observe

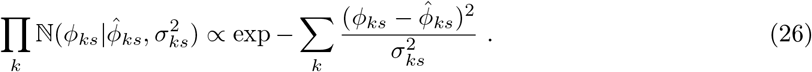

Thus, maximizing Eq. (26) is equivalent to optimizing the objective

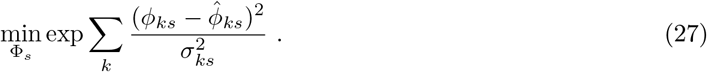

As both exp *x* and 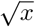 are monotonic functions, this is equivalent in turn to

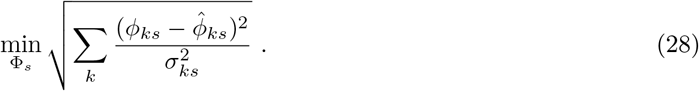

To complete the transformation of Eq. (28) to the projection objective Eq. (19), we establish the following notation.

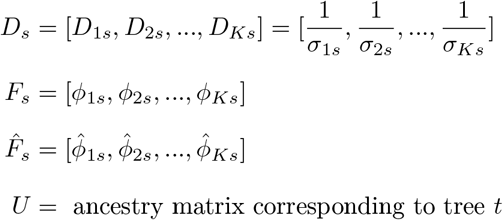

Now, Eq. (28) can be rewritten using the Frobenius norm:

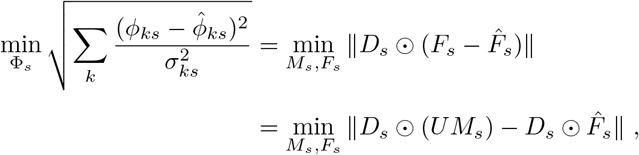

Thus, we can now call the projection optimization algorithm to compute *F_S_* and *M_S_*, which are *K*-length vectors representing tree-constrained subclonal frequencies and subpopulation frequencies, respectively. Both obey the constraints inherent to the tree *t* that are expressed through the ancestry matrix *U*. The *F_s_* values are the MAP under the Gaussian approximation Eq. (25) of binomial likelihood Eq. (22), ultimately achieving a near-optimal solution to the original optimization objective Eq. (20).

### 9.5 Clustering mutations into subclones

#### 9.5.1 Clustering overview

Pairtree takes as input a clustering of mutations into subclones. Pairtree provides two mutation clustering algorithms for grouping mutations into subclones. Mutation clusters may also be generated by other methods. Alternatively, Pairtree may be run on the mutations directly without first clustering them into subclones, yielding a *mutation tree* instead of a clone tree. A mutation tree is equivalent to a clone tree in which each clone bears only a single distinct mutation, such that every tree node corresponds to a unique mutation.

Both of Pairtree’s mutation-clustering algorithms use a Dirichlet process mixture model (DPMM) and perform inference via Gibbs sampling (see also [32, 39]). The algorithms differ in how they define their probabilistic clustering models. Let Π = {*π*_1_, *π*_2_, …, *π_M_*} represent a clustering of *M* mutations into *K* clusters, with *π_i_* indicating the assignment to a cluster of mutation *i*, such that *π_i_* ∈ {1, 2, …, *K*}. Each cluster corresponds to a genetically distinct subclone. By virtue of using a DPMM, *K* is not fixed, but instead inferred from the data.

Let *x* represent the mutation read count data. From these, we will define the posterior distribution over clusterings

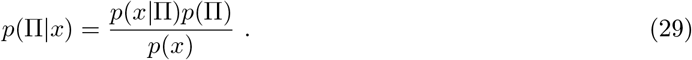

Each clustering model defines its own likelihood *p*(*x*|Π), but uses the same clustering prior *p*(Π). The clustering prior draws on the DPMM concentration hyperparameter *α*, representing the cost of placing a mutation in a new cluster relative to adding it to an existing cluster. For *K* clusters over *M* mutations, with *n_k_* mutations in cluster *k*, we define

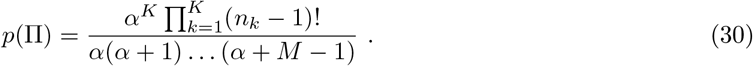

Both clustering models use Gibbs sampling, such that each clustering iteration resamples the cluster assignment of each mutation individually, conditioned upon the assignments of all other mutations. Thus, we wish to compute 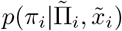, where *π_i_* indicates the cluster assignment of mutation *i*, Π is the cluster assignments of all mutations including *i*, and 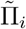 represents the cluster assignments of all mutations excluding *i*, such that 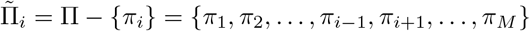.

By representing the data associated with all mutations except *i* with 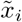, we get

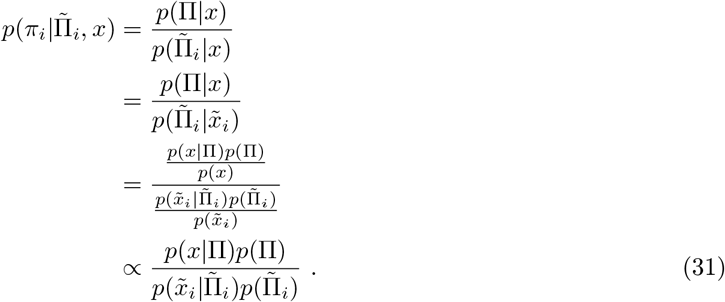

In Eq. (31), we use Eq. (30) to establish

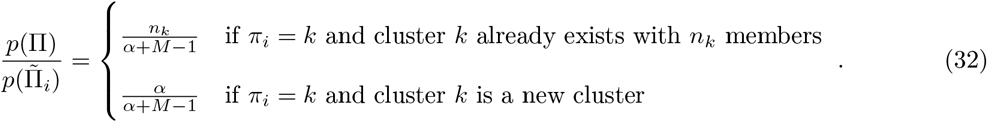

To complete Eq. (31), we need only define 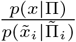. We leave this definition to the clustering models described in Section 9.5.2 and Section 9.5.3. Once this factor is defined, we can compute 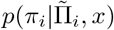 because we have in Eq. (31) a quantity proportional to it.

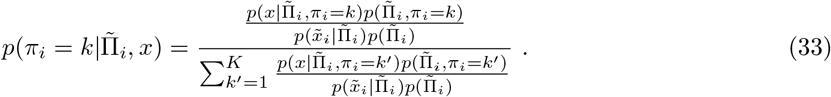

We use this definition to perform Gibbs sampling, as described in Section 9.5.4.

#### 9.5.2 Clustering mutations using subclonal frequencies

For each mutation *i* in each cancer sample *s*, we have a variant read count *V_is_*, reference read count *R_is_*, total read count *T_is_* = *V_is_* + *R_is_*, and probability of observing the variant allele *ω_is_*. To cluster mutations using subclonal frequencies, we first define for each mutation *m* in each cancer sample *s* an adjusted total read count 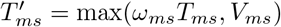. Thus, 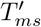 represents the (potentially fractional) number of reads originating from the variant allele across all cells, regardless of whether the reads include mutation *m* on that allele. The complete data likelihood is then defined using the following notation:

- *S*: number of cancer samples
- *K*: number of clusters
- *M*: number of mutations
- *ϕ_ks_*: subclonal frequency of cluster *k* in sample *s*
- *C_k_* ⊆ {1, 2, …, *M*}: set of mutations assigned to cluster *k*, with 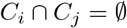 for all *i* and *j*

This yields the complete data likelihood

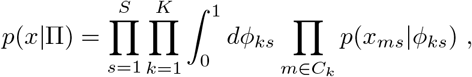

with 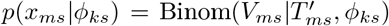. Strictly speaking, as 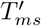 may take a fractional value, it may not be a valid parameter choice for the binomial. Nevertheless, for computational convenience, we compute the integral over the binomial using the beta function, which allows for continuous 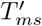 values. Consequently, we have

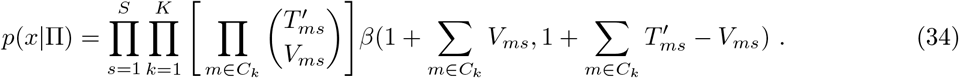

By Eq. (33), we need only define 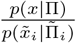 to complete the definitions required for Gibbs sampling. This follows easily from Eq. (34), yielding

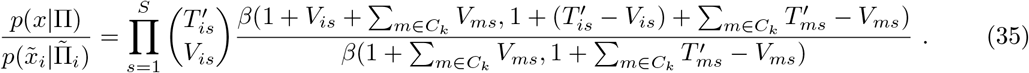

This allows us to proceed with Gibbs sampling, as described in Section 9.5.4.

#### 9.5.3 Clustering mutations using pairwise relations

As an alternative to clustering with subclonal frequencies, we can cluster mutations using the pairwise relations described in Section 9.2. To do so, we compute the posterior distributions over pairwise relations for every pair of individual variants *A* and *B*, rather than the supervariants defined from an established clustering that are used for tree search. Computing the pairwise posterior distributions over relationships *θ_AB_* necessitates that we first redefine the pairwise prior described in Section 9.2.6 to permit non-zero mass on the *coincident* relationship. For this, we allow the user to set a constant *P* representing the prior probability that mutations *A* and *B* are coincident, with 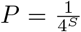 for *S* cancer samples by default, yielding

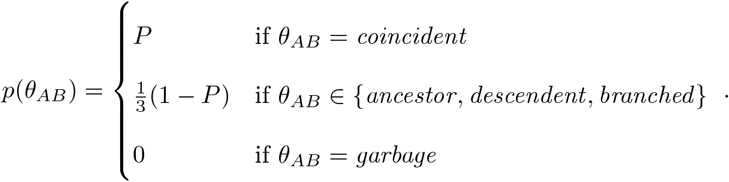

We define *p*(*θ_ab_* ≠ *coincident*|*x*) = 1 – *p*(*θ_ab_* = *coincident*|*x*). After computing these pairwise relation posteriors for every mutation pair (*a, b*) ∈ {1, 2, …, *M*} × {1, 2, …, *M*} with *a* > *b*, we can define the clustering data likelihood as

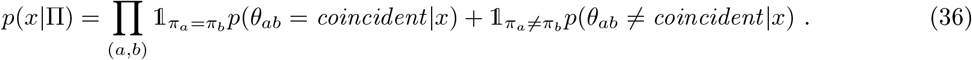

As we consider every pair (*a, b*) without also including the pair (*b, a*), there are 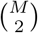 factors in the product for *M* mutations. This notation relies on the indicator function

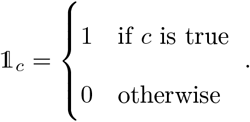

From this, we can define 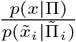, completing the definitions required for Gibbs sampling.

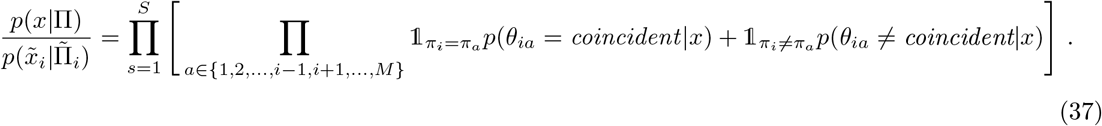

Thus, 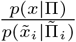 is a product over the *S* cancer samples and *M* – 1 pairs that include mutation *i*. This allows us to proceed with Gibbs sampling, as described in Section 9.5.4.

#### 9.5.4 Performing Gibbs sampling

Pairtree clusters mutations using Gibbs sampling, drawing on the probabilistic framework given in Eq. (33), and the subclonal frequency likelihood Eq. (35) or pairwise relationship likelihood Eq. (37). The primary advantage of the subclonal frequency model is that, unlike the pairwise model, it does not require the time-intensive computation of the pairs tensor before clustering can begin. The pairwise model, conversely, can be easily applied to data types other than bulk sequencing that can be represented within the pairwise relation framework, such as single-cell sequencing.

By default, the algorithm runs a total of *C* chains, with *C* set to the number of CPU cores present on the system by default, and *P* = *C* executing in parallel. Both *P* and *C* can be customized by the user. Each chain takes 1000 samples by default, which can also be changed by the user. Unlike the tree search algorithm, the clustering algorithm makes no attempt to discard burn-in samples from each chain. As tree search relies on a single clustering common to all trees, we select the clustering result with the highest posterior probability as the algorithm’s output. Nevertheless, the user could easily adapt the implementation to represent different possible clusterings alongside their posterior probabilities, conferring insight into multiple possible solutions.

The subclonal frequency and pairwise relationship clustering models use different clustering initializations, purely as an implementation artifact. The subclonal frequency models simply assigns all variants to a single cluster. Conversely, the pairwise relationship model places each variant in a separate cluster. Alternative, the pairwise model also permits the user to specify an initial clustering to use for initialization. In this case, user-specified clusters can be merged, but will never be split, such that the user can force multiple variants to always remain in the same cluster.

Two hyperparameters affect clustering results. The first, *α*, is used in Eq. (30), with higher values corresponding to an increased number of clusters. Let 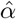 be the value provided by the user as input to the algorithm. Given a dataset with *S* cancer samples, The *α* value used in Eq. (30) is computed from this as 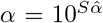, with 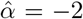 by default. Representing *α* on a logarithmic scale via 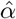 makes representing especially large or small values of *α* more convenient for the user, while scaling it with *S* ensures that the algorithm’s preference for placing data points in new clusters is unaffected by the magnitude of posterior weight contributed by data likelihood factors—i.e., each cancer sample-specific likelihood is effectively weighted by its own 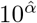 prior factor in computing the posterior. Finally, to prevent numerical issues, we force *α* ∈ [exp(−600),exp(600)].

The second clustering hyperparameter is *P*, the prior probability of two mutations being coincident (Section 9.5.3). Similar to how the *α* parameter is specified, the algorithm ensures that the number of cancer samples *S* does not affect the algorithm’s preference for starting new clusters by taking as input 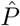, with 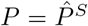. By default, we take 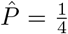, such that we enforce a uniform distribution over the four possible pairwise relations for each cancer sample.

### 9.6 Detecting garbage mutations

#### 9.6.1 Perfect phylogenies, the infinite sites assumption, and the four-gamete test

To simplify subclonal reconstruction, algorithms make the ISA, which posits that the genome is so large as to be effectively infinite in size, meaning that each genomic site is mutated at most once during the cancer’s evolution. This implies that the same site can never be mutated twice by separate events, and that it can never return to the wildtype. Moreover, two cells bearing the same mutation are assumed to share a common ancestor in which that mutation occurred. Critically, the ISA allows us to characterize more subclones than we have cancer samples. In addition, the ISA is necessary to infer the pairwise relationships between mutations from their frequencies (Section 9.2).

Most clone tree reconstruction algorithms make the ISA so that cancer phylogenies become *perfect phylogenies*, such that descendant subclones inherit all the mutations of their ancestors. Given complete genomes for each cancer cell, a perfect phylogeny can be constructed in linear time [49]. However, the bulk-tissue DNA sequencing data commonly used today do not provide complete genomes. Instead, the samples consist of mixtures of different subclones, rendering NP-complete the construction of a perfect phylogeny consistent with the exact subclonal frequencies of mutations across multiple samples [50]. Nevertheless, the ISA implies relationships between mutation frequencies that can assist subclonal reconstruction. Firstly, mutations in ancestral subclones must always have subclonal frequencies at least as high as those in descendent subclones, across every observed cancer sample. Secondly, two mutations on different tree branches can never have frequencies that sum to greater than one in any sample. This information is critical to building clone trees, as it hints at evolutionary constraints that provide a partial order over mutations (i.e., a mutation *B* cannot be the ancestor of another mutation *A* if the subclonal frequency of *B* is lower than *A* in any sample from the cancer), and that imply that some mutations must occur on the same evolutionary lineage (i.e., they cannot be on different tree branches).

#### 9.6.2 Detecting ISA violations and other erroneous mutations

Though the ISA is valid for most mutations [27], violations of the ISA occur sporadically in cancer [26]. As Pairtree relies on the ISA when converting mutation allele frequencies to subclonal frequencies and building clone trees, we wish to detect and remove such mutations before building the clone tree. Additionally, other factors such as missed CNA calls or technical noise can skew this conversion and result in mutations that should be excluded from the tree. Pairwise relationships between mutations can reveal both ISA violations and other types of erroneous mutations, which we refer to collectively as *garbage mutations*. Detecting ISA violations using these relations is similar to applying the four-gamete test (FGT) [45].

Given a mutation pair (*A, B*), we can view the noise-free subclonal frequencies *ϕ_As_* and *ϕ_Bs_* for this pair in a cancer sample *s* as eliminating possible relationship types. In Section 9.2.3, we established the constraints that the subclonal frequencies for a mutation pair (*A, B*) must obey to fulfill the ancestor, descendent, or branched relations if those mutations obey the ISA. Equivalently, we can use these constraints to rule out possible relationships in sample *s*–for instance, if *ϕ_As_* < *ϕ_Bs_*, we can eliminate the possibility that *A* is ancestral to *B*. Provided both mutations obey the ISA, at least one relationship type will be consistent with the subclonal frequencies of the mutations across all cancer samples. However, if no relationship is consistent across samples, we can deduce either that one or both mutations do not obey the ISA, or some other factor has led to inaccurate observations of their subclonal frequencies.

We consider four possible causes of garbage mutations, the first two of which stem from ISA violations.

- *Back-mutation* to wildtype occurs when a mutation is lost and reverts to wildtype.
- *Homoplasy* occurs when a mutation is acquired multiple times on different tree branches, rather than being inherited from a common ancestor.

The second two types are not ISA violations, but nevertheless result in erroneous observations of subclonal frequencies that would compromise the accuracy of clone tree reconstruction.

- *Incorrect ploidy* occurs when the wrong variant read probability *ω_vs_* is provided for variant *v* in sample *s*. These may arise, for instance, from an uncalled instance of loss of heterozygosity (LOH) or other missed CNA calls.
- *Technical noise* is a catch-all category for other erroneous mutations. While the other three categories allow in principle their mutations to be placed in the clone tree once the errors are corrected, mutations arising from this category cannot.

Pairtree does not attempt to differentiate between these categories. Instead, Pairtree tries to flag garbage mutations arising from any of these causes so they can be removed from dataset—otherwise, forcing them into the tree could skew the tree’s structure and subclonal frequencies. Potentially, the frequencies of such mutations could be corrected after tree construction, allowing the mutations to be added to the tree post-hoc.

Pairwise relations between mutations allow us to identify when the subclonal frequency estimates in different samples do not support a consistent evolutionary relationship between the pair (Section 9.2). Multiple cancer samples are necessary to identify such failures. To use pairwise relationships to detect and discard garbage mutations, we use an iterative greedy algorithm. Notably, pairwise relationships are symmetric, such that if the mutation pair (*A, B*) is declared garbage, it is impossible to decide from this alone whether *A* or *B* is the garbage mutation. However, when we consider the pairwise relationships between a putative garbage mutation and non-garbage mutations, the garbage mutation should have high posterior probability of the garbage relationship between itself and many legitimate mutations. This will be true particularly when, given a high-posterior-garbage mutation pair (*A, g*) in which *A* is legitimate and *g* is garbage, the mutation *A* arises from a subclone possessing many legitimate mutations with nearly equal subclonal frequencies. All mutations in *A*’s subclone should be deemed garbage with respect to *g* but not-garbage with respect to most other mutations, making *g*’s garbage nature apparent. Consequently, in each iteration of our garbage detection algorithm, our intent is to nominate as garbage whatever mutation has the highest probability of being garbage with respect to the pairs it forms to all other mutations.

The algorithm uses two hyperparameters denoted *γ* and *ρ*. *γ* is the prior probability assigned to the garbage relationship for each mutation pair (Eq. (8)), while *ρ* is the maximum permitted pairwise garbage probability amongst all non-garbage mutations. Let *N_i_* represent the number of non-garbage mutations at iteration *i* of the algorithm. For each mutation pair (*A, B*), we have the probability *p*(*θ_AB_* = garbage|*x_A_, x_b_, γ*) that the corresponding pairwise relationship is garbage. At each iteration *i*, we first check whether all mutation pairs (*j, k*) have *p*(*θ_jk_* = garbage |*x_j_, x_k_, γ*) ≤ *ρ*. If so, the algorithm terminates and returns the current set of *N_i_* mutations as non-garbage, rendering all others garbage. Otherwise, some pairwise garbage probability exceeds *ρ*, so we compute for each mutation *j* the joint probability that it is garbage with respect to every other mutation

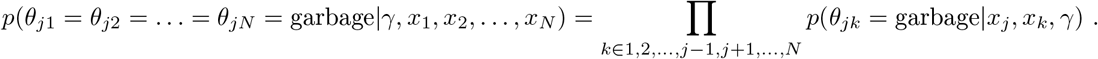

Then, we select the specific mutation *j* with

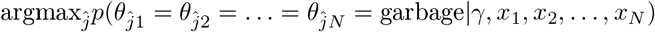

as the most likely garbage mutation. We mark this mutation as garbage to reduce the non-garbage et to size *N_i_* – 1, then continue with iteration *i* + 1 of the algorithm.

#### 9.6.3 Generating simulated datasets that include garbage mutations

To generate simulated data that includes garbage mutations, we first generate a clone tree with legitmate mutations using the same procedure as for the other Pairtree simulations (Section 9.8.2). We enerate simulated garbage mutations for the homoplasy, back-mutation, and incorrect-ploidy categories Section 9.6.2) by proposing subclonal frequencies for a putative garbage mutation based on the noise-ree frequencies of existing non-garbage mutations in the tree. Each proposed garbage mutation is then ccepted or rejected according to whether it fulfills several garbage criteria described subsequently. The nethod for generating the proposed garbage mutation *g*’s subclonal frequencies depends on its category.

1. For *back-mutations*, we uniformly sample a mutation pair (*A, B*) that has *B* as a descendant of *A*. In every cancer sample *s*, we set *ϕ_gs_* = *ϕ_As_* – *ϕ_Bs_*. We are assured that *ϕ_gs_* ≥ 0 for all cancer samples *s* because *A* is ancestral to *B*. The frequencies of *g* thus incorporate two mutations events, with *A* reflecting the acquisition of a mutation and *B* reflecting its reversion to wildtype.
2. For *homoplasy*, we uniformly sample a mutation pair (*A, B*) such that *A* and *B* occur on different tree branches. In every cancer sample *s*, we set *ϕ_gs_* = *ϕ_As_* + *ϕ_Bs_*. We are assured that *ϕ_gs_* ≤ 1 for all cancer samples *s* because *A* and *B* were branched with respect to each other and shared a common ancestor. The frequencies of *g* thus incorporate two mutations events, with *A* and *B* representing two independent events that produced the same mutation.
3. For *incorrect ploidy*, we simulate the effect of being provided the incorrect variant read probability. To do so, we uniformly sample a mutation *A*, then set *ϕ_gs_* = *ϕ_As_* for all cancer samples *s*. In our simulated event, we suppose that the locus of the true mutation *g* suffered an LOH event before *g* occurred, such that every cell carrying *g* does not contribute copies of the wildtype allele. Thus, the true variant read probability is *ω_gs_* = 1 for all samples *s*. However, we suppose that Pairtree is not given knowledge of this LOH event, but is instead provided an incorrect variant read probability 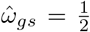, as would occur for diploid cells that were not subject to LOH. As the observed allele fraction in each sample *s* will be *ω_gs_ϕ_gs_*, the apparent subclonal frequency of *g* will be 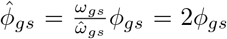. As no subclonal frequency may exceed 1, we set the apparent subclonal frequency in every sample *s* to the constrained value 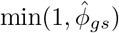.
4. For *technical noise*, we sample the subclonal frequency *ϕ_gs_* for each cancer sample *s* from the Uniform(0, 1) distribution.

Once a potential garbage mutation *g* has been proposed using the above procedure, we accept it only if it satisfies several garbage criteria with respect to legitimate mutations in the tree. This ensures that the simulated data confer enough information to detect *g* as garbage using pairwise relations relative to legitimate mutations. As the garbage detection algorithm will have access only to noisy observations of the subclonal frequencies, we set a scalar parameter Δ lying on the unit interval that ensures the conflicting relationships across samples that render the pair garbage should be clearly visible. By drawing on pairwise constraints (Section 9.2.3), we declare the pair (*m, g*) to be apparent garbage if the following three criteria are satisfied.

1. There exist a cancer sample *s* such that *ϕ_gs_* – *ϕ_ms_* > Δ. As this implies *ϕ_gs_* > *ϕ_ms_*, mutation *g* cannot be a descendant of *m*.
2. There exist a cancer sample *s*″ such that *ϕ_ms′_* – *ϕ_gs′_* > Δ. As this implies *ϕ_ms′_* > *ϕ_gs′_*, mutation *g* cannot be an ancestor of *m*.
3. There exist a cancer sample *s*′ such that *ϕ_ms″_* + *ϕ_gs″_* > 1 + Δ. As this implies *ϕ_ms″_* + *ϕ_gs″_* > 1, mutations *g* and *m* cannot be on different tree branches.

A minimum of two cancer samples can satisfy the three criteria, as we can have *s* = *s*′ or *s* = *s*″. To ensure the simulated data confer enough information to detect garbage, we require there are at least three samples that satisfy the criteria for each of *s, s′*, and *s″*. Given the potential overlap between *s* and *s*′, or between *s* and *s*″, this requires a minimum of six simulated cancer samples. Additionally, we require that, for putative garbage mutation *g*, there exist at least three distinct legitimate mutations that take the role of mutation *m* in the garbage criteria. This criterion is easy to satisfy—as described in Section 9.6.2, because all legitimate mutations within a subclone share the same noise-free subclonal frequencies, if any one of them satisfy the garbage criteria with respect to *g*, then all the subclone’s mutations will.

Given the above framework, we simulated garbage-containing clone trees using the following settings.

- Number of subclones: *K* ∈ {10, 30}
- Total number of legitimate mutations: 20*K*
- Total number of garbage mutations: 2*K*
- Number of cancer samples: *S* = 30
- Read depth: *T* = 1000
- Minimum garbage subclonal frequency difference: Δ ∈ {0.0005, 0.1}
- Number of datasets for each of four types of garbage mutation, for each number of subclones: 20

Thus, we generated 20 × 4 × 2 × 2 = 320 simulated datasets. In generating garbage datasets, some simulated tree structures could not support garbage mutations in all categories. For instance, a linear tree without any branches could not provide branched mutations to serve as the basis for simulated homoplasy. Thus, if several thousand attempts to generate the requisite number of garbage mutations proved unsuccessful for a given tree structure, that tree was discarded and a new tree was generated.

#### 9.6.4 Evaluating Pairtree’s ability to detect garbage mutations

Pairtree exhibits the ability to accurately differentiate between garbage and legitimate mutations using pairwise relations (Fig. S19). In each mutation category, for each parameter setting, we summed the counts of true positives, false negatives, and false positives to produce single values for precision and recall. Each bar in Fig. S19 thus represents performance on 20 different trees. For these experiments, we used garbage detection hyperparameters *γ* = 0.2 and *ρ* = 0.01. The garbage detection algorithm achieved a precision of 98% or higher and a recall of 88% or higher across conditions. Recall is generally lower than precision, with the worst recall of 88% occurring for ISA-violating mutations (i.e., homoplasy or wildtype back-mutations) on 30-subclone datasets simulated with Δ = 0.0005.

Three trends are notable.

1. Similar performance is achieved across the first three types of garbage mutations, including homoplasy, wildtype back-mutations, and incorrect ploidy. This is unsurprising, given that all garbage mutations were generated using a simulation process that required them to fulfill the same pairwise garbage criteria that rules out non-garbage relationships to legitimate mutations. The algorithm does best with technical noise mutations, where it achieves precision and recall of 100% for both 10- and 30-subclone trees using both settings of Δ. Such mutations clearly do not belong in the tree, since their subclonal frequencies are sampled in every cancer sample without regard for tree structure. The other three classes of garbage mutation are more subtle, given that they are based on legitimate mutations that were placed in the tree.
2. Performance on 30-subclone trees is worse than 10-subclone trees. This is expected because, within a single cancer sample, the 30-subclone trees have smaller differences in the subclonal frequencies for different subclones than the 10-subclone trees. Given that both settings used a read depth of 1000, it should be harder to infer these subclone frequency differences for the 30-subclone trees, and to discern whether those differences arise from one or both mutations being garbage, or from legitimate differences for mutations that belong in the tree.
3. We see worse performance for Δ = 0.0005 than Δ = 0.1. The subclonal frequency differences implying garbage become smaller for the lower value of Δ, and so this behaviour is anticipated. More interesting is how good performance remains at Δ = 0.0005. Using this difference in subclonal frequency, we can compute the expected number of reads this value corresponds to. Given a legitimate mutation *m* and garbage mutation *g*, with *V_ms_* and *V_gs_* variant reads in cancer sample *s*, we can use our binomial observation model to calculate this expectation. Suppose both mutations have the same variant read probability 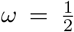 and number of total reads *T* = 1000, matching the values used for our simulations. By the linearity of expectation, we have 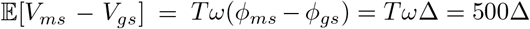. This gives an expected difference of 0.25 reads for Δ = 0.0005 and 50 reads for Δ = 0.1. The high precision and recall, even with an expected difference of less than one read between garbage and legitimate variants, suggests that garbage mutations become apparent even for small differences in implied subclonal frequencies when compiled over many samples and with respect to many mutations pairs.

In considering Pairtree’s performance for different choices of the garbage detection hyperparameters *γ* and *ρ*, respectively representing the garbage prior and maximum pairwise garbage, we see that garbage detection is most sensitive to *γ* and less so to *ρ* (Fig. S20). These experiments were conducted with Δ = 0.01. Generally, increasing *γ* results in a greater propensity to declare mutations garbage, leading to higher recall and lower precision. The best balance of precision and recall is achieved for *γ* = 0.2 for both 10 and 30 subclones, corresponding to a uniform prior over the four legitimate pairwise relationship categories considered alongside garbage. *ρ* has less effect on algorithm performance, with different *ρ* values affecting performance substantially only when *γ* = 0.5. Thus, for the experiments depicted in Fig. S19, we selected *γ* = 0.2 and *ρ* = 0.01 as the best choices.

#### 9.6.5 Detecting garbage mutations resulting from uncalled LOH

Some cases of incorrect ploidy cannot be detected using the pairwise framework. Specifically, these occur when a mutation *g* in a sample *s* has a variant allele frequency VAF_*gs*_ and variant read probability *ω_gs_* such that the implied subclonal frequency 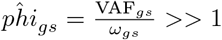. This problem often arises in the case of missed CNA calls, particularly when an (uncalled) LOH event occurred to remove the wildtype allele in a lineage and leave only the variant allele. As the MAP estimate for *g*’s subclonal frequency will be very nearly 1 in affected samples, the distributions over pairwise relationships will render mutation *g* as an almost-certain ancestor of every legitimate mutation in affected samples. If most or all cancer samples are affected by this uncalled LOH, the mutation *g* will have a high posterior probability of being an ancestor to most legitimate mutations, while having only a small probability of being garbage. Though existing approaches can build cancer phylogenies from single-cell data while accounting for LOH events [31], the method described here can detect LOH-affected mutations in bulk DNA-sequencing data.

To address this, we take an orthogonal approach using Bayes factors. Suppose mutation *g* in sample *s* has *V_gs_* variant reads and *R_gs_* reference reads. We establish a quantity proportional to the posterior probability of variant read probability *ω_gs_* as

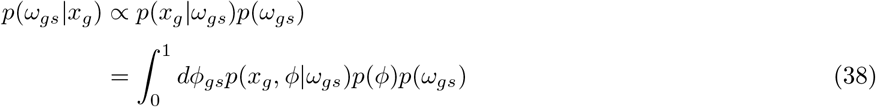

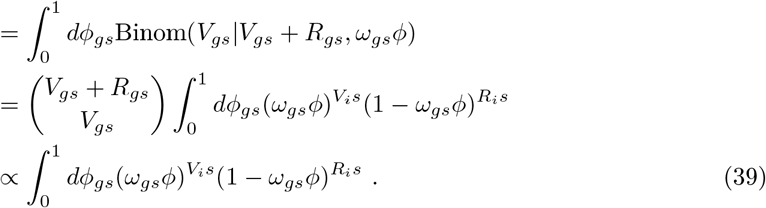

In Eq. (38), we set a uniform prior over *ϕ* and *ω_gs_* so that *p*(*ϕ*) = *p*(*ω_gs_*) = 1. Let *ψ_gs_* = *ω_gs_ϕ* so that we can integrate Eq. (39) by substitution. Using *β* to represent the incomplete beta function, this gives us

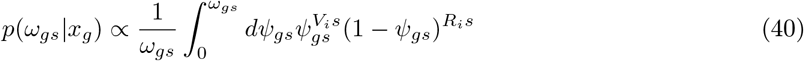

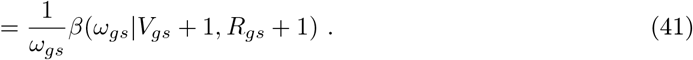

Now, suppose we are given as input the variant read probability *ω_gs_*. We wish to determine if this provided *ω_gs_* is incorrect, with a corrected 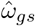 used for comparison. Here we will use 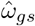 to represent the case of uncalled LOH. We compute

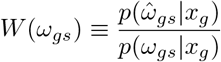

as the Bayes factor representing how much more likely the model using the corrected 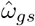 is relative to the provided *ω_gs_*. Using a configurable threshold *T*, we ask if there exists any cancer sample *s* with 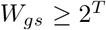, setting *T* = 10 by default. Thus, we determine if the model using the corrected variant read probability is approximately 1000 times as likely as the model using the provided probability. If 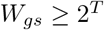 for any cancer sample *s*, we label the mutation *g* as garbage.

For simple scenarios such as an uncalled LOH event, the user can correct the provided *ω_gs_* to reflect the event. Such errors are particularly obvious when the cancer sample’s purity (i.e., what proportion of sequenced cells originate from the cancer) is high, as the implied subclonal frequencies 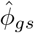 will be nearly equal to 2. Regardless, this test is often worth using before running the full Pairtree algorithm, as test failures point to likely erroneous data. Because the test does not require computing pairwise relationships, it is computationally cheap to execute.

#### 9.6.6 Generating simulated datasets with mutations subject to uncalled LOH

To evaluate Pairtree’s ability to detect mutations whose implied subclonal frequencies are skewed by uncalled CNA, we generated another set of trees. As in Section 9.6.3, we simulated the effect of being provided the incorrect variant read probability. From an existing tree with multiple non-garbage mutations, we uniformly sample a mutation A, then set *ϕ_gs_* = *ϕ_As_* for all cancer samples *s*. In our simulated event, we suppose that the locus of the true mutation *g* suffered an LOH event before *g* occurred, such that every cell carrying *g* does not contribute copies of the wildtype allele. Thus, the true variant read probability is *ω_gs_* = 1 for all samples *s*. However, we suppose that Pairtree is not given knowledge of this LOH event, but is instead provided an incorrect variant read probability 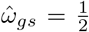, as would occur for diploid cells that were not subject to LOH. As the observed allele fraction in each sample *s* will be *ω_gs_ϕ_gs_*, the apparent subclonal frequency of *g* will be 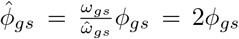. We then accept *g* as a legitimate garbage mutation only if there exists at least one cancer sample where 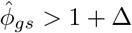, where Δ is a parameter lying on the unit interval. Unlike the “missed CNA” variants generated in Section 9.6.3, however, we do not require that the pairwise constraints between *g* and legitimate mutations appear to suggest the garbage relationship.

Given the above framework, we simulated garbage-containing clone trees using the following settings.

- Number of subclones: *K* ∈ {10, 30}
- Total number of legitimate mutations: 20*K*
- Total number of garbage mutations: 2*K*
- Number of cancer samples: *S* = 30
- Read depth: *T* = 1000
- Minimum invalid subclonal frequency difference: Δ ∈ {0.0005, 0.01}
- Number of datasets for each number of subclones: 100

Thus, we generated 100 × 2 = 200 simulated datasets. In generating garbage datasets, some simulated tree structures could not support garbage mutations in all categories. For instance, a tree with an initial normal-tissue population frequency of 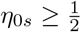 in every sample *s* would be unable to generate any garbage mutations satisfying the subclonal frequency criterion, as every mutation subclonal frequency would be less than 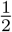. Thus, if several thousand attempts to generate the requisite number of garbage mutations proved unsuccessful for a given tree structure, that tree was discarded and a new tree was generated.

#### 9.6.7 Evaluating Pairtree’s ability to detect uncalled LOH

The missed-CNA detection algorithm produced high precision and recall for simulated LOH events (Fig. S21). We used the default Bayes factor threshold of *T* = 10 for these experiments. As in the pairwise relationship garbage detection experiments (Section 9.6.4), the number of true positives, false negatives, and false positives were summed across datasets, such that every bar represents results on 100 different trees. For most settings, precision and recall were 99% or higher (Fig. S21). The only exception occurred for recall on 30 subclones with Δ = 0.0005, where it fell to 91%. The pairwise garbage detection algorithm fared slightly better for missed CNAs, where it reached 96% (Fig. S21). Thus, performance for detecting missed CNAs without using pairwise relationships was on par with or slightly lower than the algorithm version using pairwise relationships. We must note, however, that this performance is evaluated on different datasets—unlike in the datasets used for the pairwise experiments, mutations here did not need to satisfy pairwise garbage criteria with respect to legitimate mutations, but did, however, need to exhibit at least one cancer sample with an implied noise-free subclonal frequency exceeding 1 + Δ.

These dataset differences were intended to accurately reflect the algorithms’ performance in the different settings for which they were designed. We expect that some missed-LOH mutations can be detected by the pairwise algorithm but not by the non-pairwise algorithm. These mutations would consist of ones whose implied subclonal frequencies do not exceed 1, but nevertheless violate pairwise constraints relative to legitimate mutations across multiple cancer samples. Likewise, some missed-LOH mutations can likely be detected by the non-pairwise algorithm but not the pairwise algorithm. Such mutations would have implied subclonal frequencies exceeding one in at least one cancer sample. As such high frequencies would imply the mutation is ancestral to all legitimate mutations, it would be difficult to detect such mutations using the pairwise framework, necessitating another approach. The relatively low computational cost of the non-pairwise algorithm renders it easy to run before using the pairwise garbage detection, variant-clustering, and tree-building aspects of Pairtree.

### 9.7 Using single-cell DNA sequencing data for building clone trees

Single-cell DNA sequencing (scDNA-seq) is used to study cancer evolution [51, 52]. In principle, scDNA-seq directly measures each cancer cell’s genotype, avoiding the need to deconvolve the signal from many cell subpopulations that is inherent to bulk sequencing. However, in practice, scDNA-seq data is noisy, with amplification biases giving rise to false positive and frequent false negative mutation calls (i.e., dropouts) [53]. For this reason, and the substantial additional expense of current scDNA-seq protocols relative to bulk sequencing, the latter will likely remain widely used for the foreseeable future.

Nevertheless, it is possible in principle to extend Pairtree to construct clone trees from single-cell DNA sequencing (scDNA-seq) data. Doing so would require modifying Pairtree’s pairwise relation framework to incorporate binary mutation observation processes inherent to scDNA-seq, replacing the current processes based on mutation’s estimated subclonal frequencies in bulk sequencing, and changing the tree likelihood function used during the MCMC tree inference procedure. Similar extensions could be used to infer clone trees from mixtures of scDNA and bulk data (see, e.g., [20]).

We have demonstrated that Pairtree can accurately recover clone trees with more subclones than cancer samples by deconvolving bulk samples. This deconvolution ability could also be useful in detecting and resolving cell doublets. Another potential extension of Pairtree to scDNA-seq would be to combine multiple cells to generate quasi-bulk data.

### 9.8 Creating simulated data

#### 9.8.1 Parameters for simulating data

We first define parameters characterizing the different simulated cancers.

We created simulated datasets with the following parameter combinations.

**Table 2:** Simulated data parameters. All combinations of these parameter values were used to generate simulated data, excepting cases when *K* ∈ {30,100} and *S* ∈ {1, 3}. This provided 144 parameter combinations, with four datasets generated from each, yielding 576 simulated datasets.

| Parameter | Values |
| --- | --- |
| $K$ | 3, 10, 30, 100 |
| $S$ | 1, 3, 10, 30, 100 |
| Mutations per cluster | 10, 20, 100 |
| $T$ | 50, 200, 1000 |

Observe there are 4×5×3×3 = 180 parameter combinations. When *K* ∈ {30,100}, we did not simulate datasets with *S* ∈ {1, 3} samples, as trees with so many subpopulations and so few cancer samples are unrealistic—to resolve a large number of distinct mutation clusters, a large number of cancer samples is typically needed. Simulated datasets with *K* ≥ 30 and *S* < 10 would thus correspond to complex trees with few cancer samples, posing a highly underconstrained computational problem that would not reflect how methods perform on realistic datasets. Thus, as there are 2 × 2 × 3 × 3 = 36 parameter sets yielding under-constrained simulations, we used the remaining 180 – 36 = 144 sets to generate simulations. For each valid parameter set, we generated four distinct datasets, yielding 144 × 4 = 576 simulated datasets.

Above, rather than setting the number of mutations per dataset *M* directly, we instead specified the average number of mutations per cluster. This reflects that, because each subclone is distinguished by one or more unique mutations, trees with more subclones should have more mutations. Consequently, the number of mutations generated per dataset was *M* = *K*(mutations per cluster). Nevertheless, as described in Section 9.8.2, mutations are assigned to subclones in a non-uniform probabilistic fashion, such that the number of mutations in each subclone is only rarely equal to the parameter value for number of mutations per cluster used when generating the data.

#### 9.8.2 Algorithm to generate simulated data

We generated simulated data using the following algorithm. Python code implementing this algorithm is available at https://github.com/morrislab/pearsim.

1. Generate the tree structure. For each subclone *k*, with *k* ∈ {1, 2, …, *K* – 1}, sample a parent 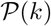. We extended the previous subpopulation (i.e., 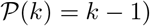) with probability *μ* = 0.75, and otherwise sample 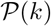 from the discrete Uniform(0, *k* – 1) distribution. This extension probability created “linear chains” of successive subpopulations, with each member of the chain taking only a single child, interrupted sporadically by the creation of new tree branches. As the normal tree root, denoted as node 0, exists at the outset, node 1 will always take it as a parent. Note that this scheme allows for the creation of “polyprimary” trees, in which the root 0 takes multiple clonal cancerous children. Such polyprimary cases are created for approximately 1 – *μ* = 25% of datasets.
2. Generate the subpopulation frequencies *η_ks_* for each subpopulation *k* in each cancer sample *s*, with *s* ∈ 1, 2, …, *S*. These values were sampled separately for each *s*, with [*η*_0*s*_, *η*_1*s*_, …, *η_Ks_*] ~ Dirichlet(*α*, …, *α*) = Dirichlet(0.1, …, 0.1). We use the symmetric Dirichlet distribution with a single α parameter because we have no reason to desire that any population frequency tend to be greater or less than others a priori. The choice of α has important implications for the structure of the simulated data (Section 9.17). As the η vector is drawn from the Dirichlet, we have 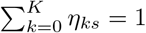 for each sample *s*.
3. Compute the subclonal frequencies *ϕ_ks_* for each subclone *k* in each cancer sample *s* using the tree structure and *η_ks_* values. Let *D*(*k*) represent the set of *k*’s descendants in the tree. Then, we have

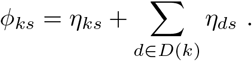
4. Assign the *M* variants to subclones. To ensure every subclones has at least one variant, set the subclones of the first *K* SNVs to 1, 2, …, *K*. To assign the remaining *M* – *K* SNVs, sample subclone weights from the *K*-dimensional Dirichlet(1,1,*…*, 1), then sample assignments from the *K*-dimensional categorical distribution using these weights.
5. Sample read counts for the variants. Let *A*(*m*) ∈ {1, 2, …, *K*} represent the subclone to which variant *m* was assigned. Let 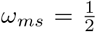 represent the probability of observing a variant read when sampling reads from the variant′s locus, for all subpopulations contained within m’s subclone, reflecting a diploid variant not subject to any CNAs. Then, for each cancer sample *s*, given the fixed total read count *T* used for all variants in a dataset, we sample the number of variant reads *V_ms_* ~ Binomial(*T,ω_ms_ϕ*_*A*(*m*),*s*_).

### 9.9 Evaluation metrics for method comparisons

#### 9.9.1 Intuitive explanation of metrics

We developed two metrics for evaluating clone-tree reconstruction algorithms that are suitable for use with multiple cancer samples. The first, termed *VAF reconstruction loss* (henceforth “VAF loss”), measures how well a tree’s subclonal frequencies match the allele frequency for each mutation implied by its CNA-corrected VAF. Each tree structure permits a range of subclonal frequencies, with the best subclonal frequencies matching the data as well as possible while also satisfying the tree constraints. Thus, the VAF loss evaluates a tree by determining how closely its subclonal frequencies match the observed data. VAF loss is reported in bits per mutation per cancer sample, representing the number of bits required to describe the data using the tree, normalized to the number of mutations and cancer samples. Lower values reflect better trees. As LICHeE could not compute subclonal frequencies itself, producing only tree structures, we used Pairtree to compute the MAP subclonal frequencies for its trees.

All evaluated methods report multiple solutions for each dataset, scored by a method-specific likelihood or error measure. To determine a single VAF loss for each method on each dataset, we used the method-specific solution scores to compute the expectation over VAF loss (equivalent to the weighted-mean VAF loss). VAF loss is always reported relative to a baseline. For simulated data, the baseline is the VAF loss achieved using the true subclonal frequencies that generated the data. For real data, the baseline is expert-constructed, manually-built trees that were subjected to extensive refinement, with Pairtree used to compute the MAP subclonal frequencies. Thus, VAF loss indicates the average extra number of bits necessary to describe the data using a method’s solutions rather than the baseline solution. Methods can find solutions that fit the data better than the baseline, yielding a negative VAF loss.

The second evaluation metric we developed, termed *relationship reconstruction error* (henceforth “re-lationship error”), recognizes that a clone tree defines pairwise relations between its constituent mutations, placing every pair in one of the four relationships discussed earlier. Using the set of trees reported by a method for a given dataset, we computed the empirical categorical distributions over pairwise mutation relations, with each tree’s relationships weighted by the likelihood or error measure reported by the method. We then compared these distributions to the distributions imposed by all tree structures permitted by the true subclonal frequencies, computing the Jensen-Shannon divergence (JSD) between distributions for each pair. This yields a relationship error ranging between 0 bits and 1 bit. Using these, we report the joint JSD across all mutation pairs to summarize the quality of the solution set, normalized to the number of pairs. Thus, the relationship error for a given dataset ranges between 0 bits and 1 bit, with smaller values indicating that a method better recovered the full set of trees consistent with the data. We did not apply this metric to real data, whose true subclonal frequencies, and thus true possible tree structures, are unavailable.

As a complement to relationship error, we also use tree error, which indicates what proportion of pairwise relationships imposed by a method’s best-scored tree match the true relations imposed by the tree structure used to generate the data. This metric does not reflect the ambiguity inherent to the problem, whereby multiple tree structures may be equally consistent with the noise-free lineage frequencies used to generate the data. Nevertheless, we provide results using this metric because it has been used in other studies [23, 43], meaning the evaluations are not performed solely using newly defined metrics.

#### 9.9.2 VAF reconstruction loss

The VAF reconstruction loss represents how closely the subclonal frequencies associated with a method’s clone tree solution set match the simulated data’s VAFs (Section 2.4). The constraints imposed by good solution trees should permit subclonal frequencies that closely match the data. In Section 9.3.2, we described the tree likelihood Eq. (9), which we also use to define the VAF reconstruction loss.

Assume the method provides a distribution over different clone trees *t*, with the posterior probability of *t* represented as *p*(*t*), such that ∑_*t*_ *p*(*t*) = 1. The loss is defined for each tree *t* over the mutation read count data *x*, with mutations *m* and cancer samples *s*. We use *ϕ_ms_* to indicate the subclonal frequency in *t* for sample *s* associated with the subpopulation containing mutation *m*. For mutation *m* in sample *s*, we define the likelihood

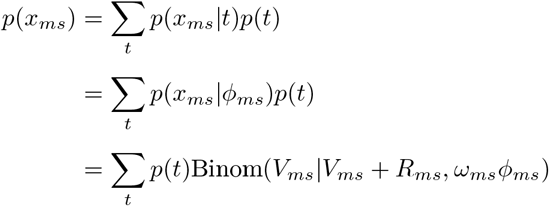

To compute the VAF reconstruction loss *ϵ*_Φ_, we calculate the mean negative log-likelihood across all *M* mutations and *S* cancer samples, with

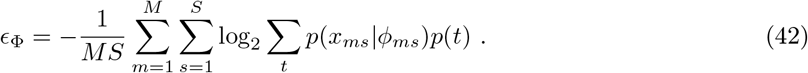

As *p*(*x_ms_*|*ϕ_ms_*) ≤ 1 and *p*(*t*) ≤ 1, given that both are discrete distributions, we have *ϵ*_Φ_ ≥ 0. We report VAF reconstruction loss relative to a baseline, though this is not necessary—the absolute metric is still useful for quantifying the error in the tree-constrained subclonal frequencies that are part of a solution set. Nevertheless, by reporting error relative to a baseline, we can more easily see how well a method is faring, given that some datasets will necessarily yield higher absolute VAF losses than others. For simulated data, we use as the baseline the true subclonal frequencies that generated the data. For real data, we use as the baseline the subclonal frequencies computed by Pairtree (Section 9.4) for our expert-derived trees. In both cases, we use Eq. (42) to compute the baseline VAF loss 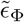, with the distribution over trees *p*(*t*) consisting of a single tree, for which *p*(*t*) = 1. This yields the relative VAF loss

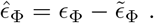

These are the values reported in this study for VAF loss. The relative VAF loss 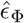 can be negative, indicating that a method has found a better solution than the baseline. On simulated data, for instance, this can occur if there is only one tree consistent with the simulated subclonal frequencies, and the clone-tree-reconstruction method finds only that tree, to which it then fits the MAP subclonal frequencies. These will necessarily fit the observed data better than the true frequencies, yielding a negative relative VAF loss.

#### 9.9.3 Relationship reconstruction error

In determining relationship reconstruction error (Section 2.4), we wish to compare the distribution over pairwise mutation relationships imposed by a method’s set of candidate solutions relative to the simulated truth. Though there was a single true tree structure used to generate the observed data, we cannot simply compare the candidate solutions to the relations imposed by this true tree—the observed VAF data are noisy reflections of the true subclonal frequencies accompanying the true tree structure, and while the true tree will be consistent with the noise-free frequencies (i.e., it will not violate the constraints they impose), there may also be other consistent tree structures. Thus, our baseline must be not the single set of relationships imposed by the true tree, but the distribution over relationships implied by all tree structures consistent with the true subclonal frequencies. Determining this baseline requires that we enumerate all such trees (Section 9.9.5). We can then measure the quality of a set of proposed solution trees by the extent to which the distribution over pairwise relations they imply recapitulates the baseline. To excel according to this metric, methods must be able to recover the full set of trees permitted by the observed VAF data, rather than only a single consistent tree. Moreover, methods must be able to deal with noise inherent to the VAF observations, such that the methods find trees that make small violations of tree constraints if we take the VAFs as exact observations of the subclonal frequencies.

Suppose a dataset consists of *M* mutations. Every clone tree built for this dataset by a method places each mutation pair (*A, B*) unambiguously into one of the four pairwise relationships. We use *θ_AB_* to delineate the pairwise model for the mutation pair induced by a given clone tree. (Provided the method uses a fixed mutation clustering provided as input, the coincident relations are determined by the clustering, and so are fixed before the method is run.) Assume the method provides a distribution over different clone trees *t*, with the posterior probability of *t* represented as *p*(*t*), such that ∑*_t_ p*(*t*) = 1. In this case, we can compute the posterior probability of the *θ_AB_* relation as *p*(*θ_AB_*) = ∑*_t_ p*(*θ_AB_*|*t*)*p*(*t*), where

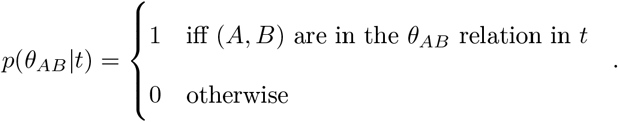

Using the set of true trees (Section 9.9.5), we will define 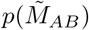 as the distribution over different relations for all *N* trees consistent with the true subclonal frequencies. For the true tree set, we will establish a uniform prior 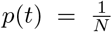, since no true tree should be privileged over another. For the mutation pair (*A, B*), we can now compute the Jensen-Shannon divergence (JSD) between a clone-tree-construction method’s *p*(*θ_AB_*) and the true 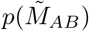, which we denote as 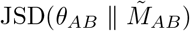. We use the base-two logarithm in computing JSD, yielding a measurement in bits.

Given *M* mutations in a dataset, there are 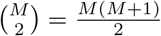 mutation pairs (*A, B*). We thus define the relationship reconstruction error *ϵ_R_* for a solution set as the mean JSD between pairs, such that

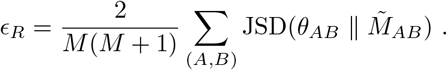

Using the mean allows us to compare *ϵ_R_* values for datasets with different numbers of mutations, so that we can understand which result sets have more or less error. As an aside, though it may be tempting to view *ϵ_R_* as the joint JSD for all mutation pairs, normalized to the number of mutation pairs, this perspective is wrong. The JSD can be defined with respect to the Kullback-Leibler (KL) divergence. Under our definition of *p*(*θ_AB_*|*t*), every pair is independently distributed, such that the KL divergence of the joint distribution over all pairs is equal to the sum of KL divergences of individual pairs. This property is not, however, true for the JSD, and so our sum over pairs does not equal the JSD of the joint distributions.

Note that relationship error is similar to the probabilistic ancestor-descendant matrix (ADM) metric developed in [24], where it is referred to as metric 3B. To represent the ground truth, given *M* mutations and a single true tree 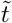, metric 3B constructs four matrices of size *M* × *M*, which can be represented by the *M* × *M* ×4 tensor denoted by *T*. Let *T_ijk_* be the binary indicator corresponding to whether mutations *i* and *j* fall into pairwise relationship *k* ∈ {ancestor, descendant, branched, coincident} (Section 9.2). Similarly, a candidate solution set can be represented with an *M* × *M* × 4 tensor denoted by *R*, with *R_igk_* indicating the probability that mutations *i* and *j* fall into relationship *k*. Both *T* and *R* are thus akin to the pairs tensor computed by Pairtree. To compute the similarity between *T* and *R*, the 3B metric concatenates the column vectors of each tensor’s constituent *M* × *M* matrices, forming vectors of length 4*M*^2^ that we denote with 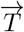 and 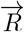. The metric 3B is then computed as the Pearson correlation between 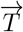 and 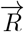, equivalent to the mean-centered cosine similarity between these vectors.

Relationship error differs from metric 3B in two ways [24]. Though both operate on information about similarity in pairwise relations between a ground truth and candidate solution set, they compute distance differently. Relationship error uses the mean JSD between all pairs, and so ranges between 0 and 1, while metric 3B uses Pearson correlation, and so ranges between −1 and 1. More importantly, relationship error’s truth is defined with respect to all trees, and thus pairwise relationships, consistent with the true subclonal frequencies. Metric 3B, conversely, defines truth with respect to the single tree structure used to generate the data. Relationship error thus better reflects a method’s performance, as it recognizes the fundamental ambiguity in tree structure.

#### 9.9.4 Tree error

Tree error complements relationship error (Section 9.9.3) to provide a metric defined solely using single trees, without accounting for a method’s ability to provide multiple solutions or the existence of multiple trees equally consistent with the noise-free lineage frequencies used to generate the observed mutation count data. Given a candidate tree *t* provided in reference to a true tree 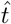, let 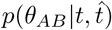 be an indicator function indicating whether the pairwise relation *θ_AB_* for mutations *A* and *B* imposed by *t* matches the relation defined by 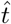. This yields

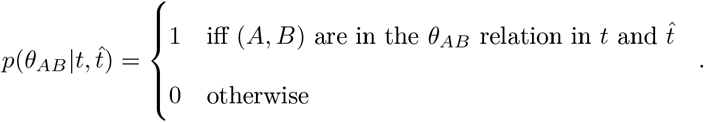

Given *M* mutations in a dataset, there are 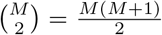 mutation pairs (*A, B*). Consequently, we define tree error 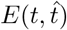 using the mean error across mutation pairs, such that

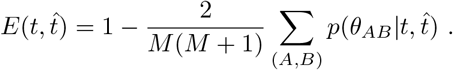

#### 9.9.5 Enumerating trees quickly

To enumerate all trees consistent with the true subclonal frequencies for a simulated dataset, henceforth termed “consistent trees,” we first construct a directed graph tau. Given *K* subclones and *S* cancer samples, tau consists of a graph of *K* + 1 nodes, with the *i*th node corresponding to the *i*th subclone, and the implicit node 0 that has no incoming edges. We place an edge from node *i* to node *j* in *tau*, such that *tau_ij_* = 1, if node *i* is a potential parent of subclone *j* in a tree consistent with the subclonal frequencies Φ = {*ϕ_ks_*}. The *τ* graph represents edges that will be present in at least one consistent tree. Thus, the spanning trees of *tau* compose a superset of the consistent trees—i.e., all consistent trees exist as a spanning tree of *tau*, but not all spanning trees of tau must be consistent trees.

By definition, *ϕ*_0*s*_ = 1 for all cancer samples *s*. Without loss of generality, assume *ϕ_is_* ≥ *ϕ*_(*i*+1)_s__ for *i* ∈ {1, 2, …, *K* – 1} for all cancer samples *s*, as the subclones can be sorted to fulfill this requirement without affecting the problem structure. We then construct *τ* as follows.

**Listing 2:**
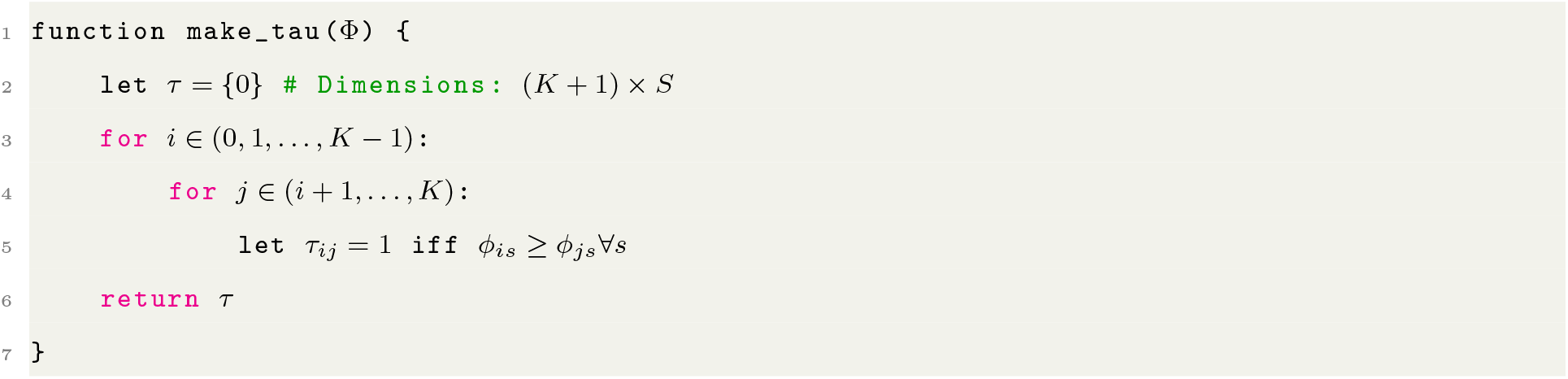
Algorithm to create graph adjacency matrix *τ*.

**Listing 3:**
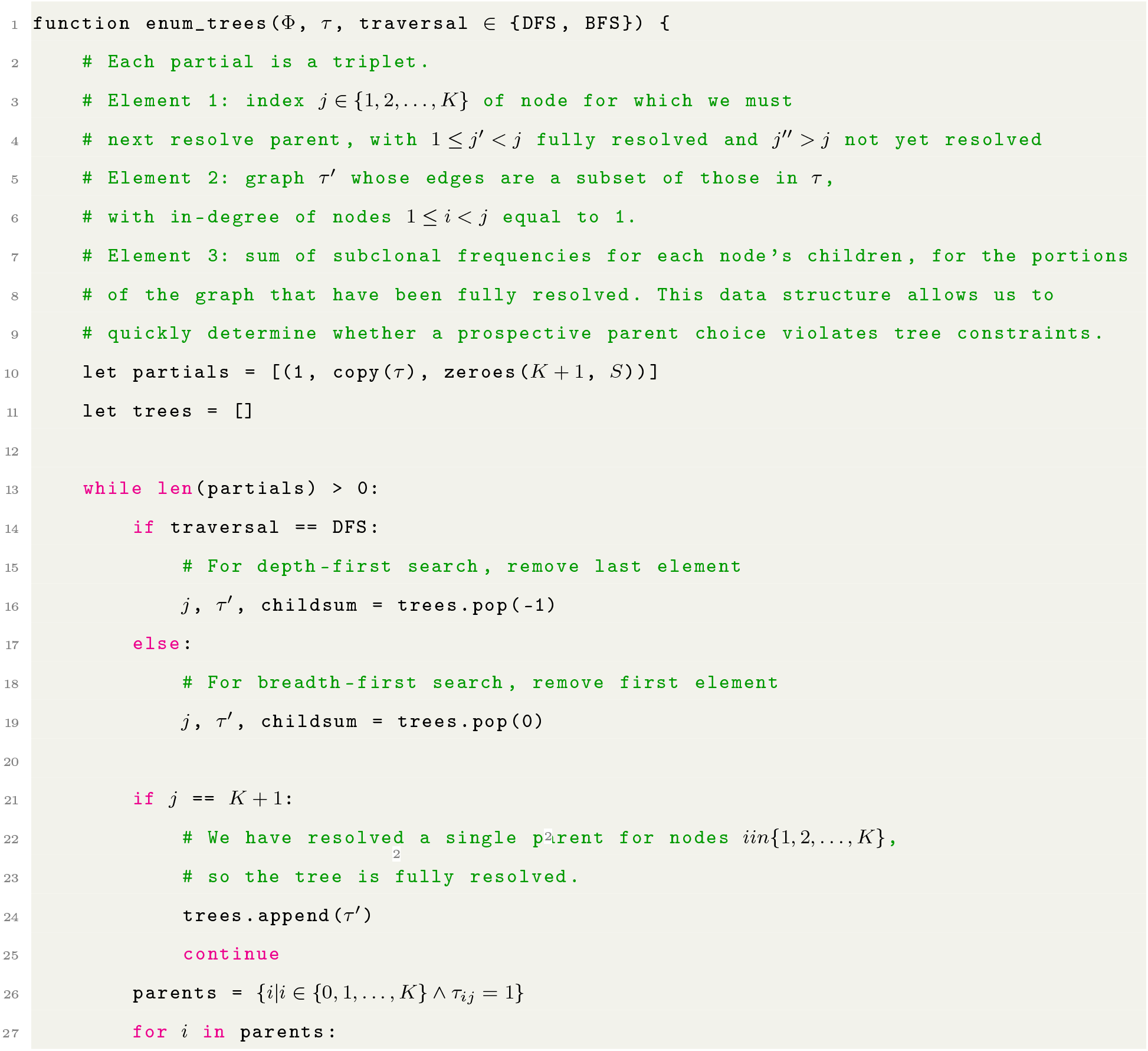
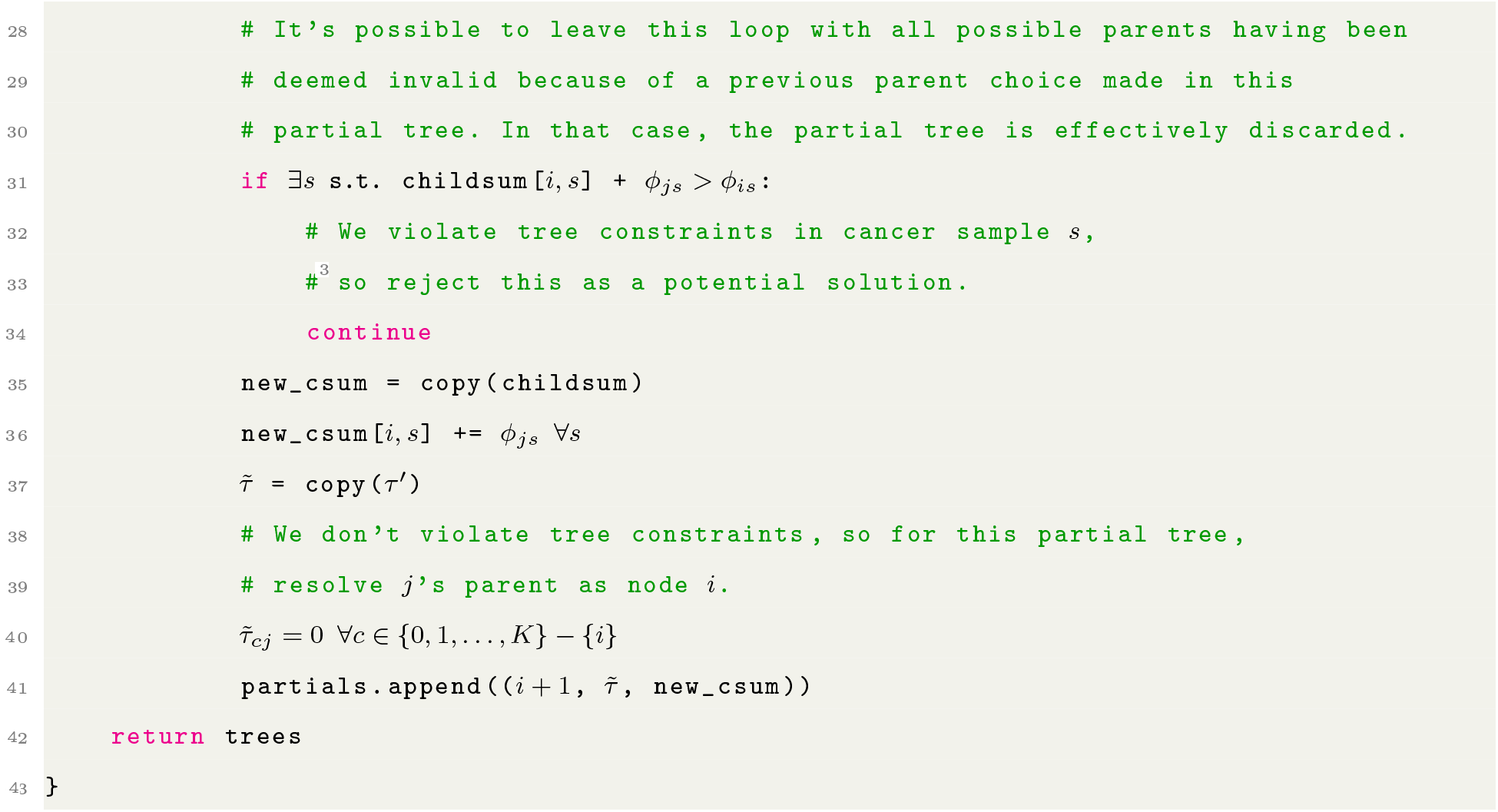
Algorithm to enumerate trees based on *τ* graph.

By implementing this algorithm in Python and exploiting Numba, we can enumerate trees for all 576 simulated datasets quickly.Improving runtime through parallelization would be trivial, given that the algorithm need make only a single pass through each *τ*′ graph, without having to backtrack “up” the graph to alter edges corresponding to fully resolved parents. Though the algorithm offers the choice of DFS or BFS when exploring the *τ* graph, DFS is generally superior. As the tree enumeration algorithm proceeds down the *τ* graph, DFS allows it to quickly determine whether a parental choice made upstream of the nodes being considered was invalid, making it impossible for a downstream node to find any parent. DFS will quickly find this parent-less downstream node and so discard the partial tree. BFS, conversely, will keep the invalid partial tree in memory as it futilely resolves parents of other nodes before locating the parent-less node, while also storing in memory other variants of the invalid partial tree that retain the erroneous parental choice. The memory demands of the BFS algorithm variant can thus be much higher than DFS, while conferring no benefit.

Additionally, we could alter the `make_tau` algorithm to remove edges that are clearly invalid before beginning enumeration. Suppose in *τ* we have a node *j* whose only possible parent is *i*, and that there exists another node *k* who is also a possible child of *i*, implying *ϕ_is_* ≥ *ϕ_js_* and *ϕ_is_* ≥ *ϕ_ks_* for all cancer samples *s*. Furthermore, suppose *ϕ_is_* – *ϕ_js_* < *ϕ_ks_* for at least one *s*. This implies that, by exploiting the knowledge that *i* must be the parent of *j*, we can eliminate *i* as being a possible parent of *k*. Moreover, by eliminating the *i*-to-*k* edge from *τ*, we may have determined with certainty the parent of *k*. Supposing this is true, we label *k*’s parent as *i*′, and can eliminate any edges from *i*′ to other possible children *k*′ that would now violate the tree constraints. In this manner, we can propagate constraints through *τ* at the algorithm’s outset to eliminate edges from consideration. We have not implemented this optimization here, as tree enumeration was already sufficiently fast for our purposes.

### 9.10 Running comparison methods

All methods were run on systems with dual Intel Xeon 6148 CPUs, with 40 CPU cores and 192 GB of RAM. Methods were allowed up to 24 hours of compute time per dataset, and were terminated if they exceeded this threshold.

We used CITUP v0.1.2 from https://anaconda.org/dranew/citup, corresponding to the most recent revision at https://bitbucket.org/dranew/citup/. CITUP offers both a quadratic integer programming (QIP) mode and a faster iterative approximation to it. We used the QIP mode because it alone was able to take a fixed clustering as input. The iterative approximation insists on clustering mutations itself, which would have unfairly disadvantaged CITUP relative to other methods, as it would not have known which mutations belonged to which clusters. Regardless, we tried running CITUP’s iterative mode with the same supervariant-based approach we used for PhyloWGS (described below), but this did not improve CITUP’s failure rate.

We used LICHeE version `26c2a701` from https://github.com/viq854/lichee. LICHeE could not compute subclonal frequencies, so we invoked Pairtree to perform this task using the tree structures LICHeE produced. LICHeE can optionally cluster mutations itself, but we gave it the correct mutation clustering as input.

We used PASTRI version `1d2fb83c` from https://github.com/raphael-group/PASTRI, which is limited to running on datasets with 15 or fewer subclones. PASTRI was given the correct mutation clusters as input.

We used PhyloWGS version `2205be16` from https://github.com/morrislab/phylowgs. PhyloWGS did not offer a means of taking a fixed clustering as input, unlike the other four methods examined, and so was disadvantaged in the method comparisons. We provided as much clustering information to PhyloWGS as possible by using *supervariants* (Section 9.3.8), preventing the method from splitting clusters such that mutations from the same cluster would be assigned to different subpopulations. Nevertheless, PhyloWGS could still merge clusters such that multiple clusters’ variants would be assigned to the same subpopulation.

### 9.11 Examining method failures

CITUP produced results for 137 of the three-subclone datasets (76%), failing on the remainder. CITUP also failed on all datasets with 10, 30, or 100 subclones. For 3- and 10-subclone failures, 137 exited with the error `failed to optimize LP`: `Infeasible`, while 34 `failed` with failed to `optimize LP`: `Unknown`. Another 52 of the 10-subclone runs failed to finish in 24 hours. All 216 datasets with 30 or 100 subclones failed with the error `create_trees failed to complete`.

LICHeE succeeded on 477 cases. its 99 failures all occurred on 100-subclone datasets, where the method failed to finish in 24 hours.

PASTRI only supports 15 or fewer subclones, and so failed on all 216 datasets with 30 or 100 subclones. For 37 datasets with 3 or 10 subclones, PASTRI succeeded in sampling at least one tree with subclonal frequencies. On 22 datasets, all of which had 10 subclones, PASTRI failed to finish within 24 hours. PASTRI terminated without sampling any trees for 220 datasets, comprising a mixture of 3- and 10-subclone cases. Additionally, on 81 datasets, PASTRI sampled one or more trees, but failed at later steps of its pipeline, without writing usable output. These 81 cases included four types of failure.

CALDER succeeded on 350 of the 576 simulated datasets. Its 226 failures reflect instances where it could not construct a graph possessing a spanning tree that corresponded to a valid clone tree.

- PASTRI failed the assertion `assert(round(slack[j],10) >= 0`) in `gabow_myers.py` for one ten-subclone case.
- PASTRI failed with a `ValueError`: `too many values to unpack` exception for other cases.
- In some cases, the trees had fewer nodes than expected, despite being given the correct number of subclones as input.
- Some cases included invalid blank lines for some of their subclonal frequencies, evidently stemming from an error when frequencies of exactly 1 were output as blanks.

PhyloWGS succeeded on 535 datasets. Amongst these, it finished all 1000 burn-in and 2500 posterior samples within 24 hours for 463. For another 72 cases, comprising a mixture of 30- and 100-subclone datasets, it finished the burn-in samples and at least one posterior sample, without finishing all 2500 posterior samples. These 72 cases were counted as successes, but assigned wall-clock times and CPU times of 24 hours (Section 9.15.2). The remaining 41 runs failed to complete their burn-in portion within 24 hours, and so were counted as failures. All such cases had 100 subclones.

### 9.12 Using tree error to evaluate performance without considering ambiguity in the solution space

Pairtree consistently shows near-zero tree error in settings with three or 30 subclones (Fig. S4). Tree error is higher with 10 subclones because, when only a small number of cancer samples are present for such datasets, many possible trees exist (Fig. S14a), which Pairtree captures and scores appropriately. By arbitrarily selecting one of these trees, the tree error metric does not reflect this ambiguity. This becomes evident when comparing to relationship error (Fig. 3c), which is near-zero in the same 10-subclone setting, reflecting both Pairtree’s ability to characterize ambiguity inherent to the problem, as well as the importance of using metrics that incorporate uncertainty.

As with the others metrics, Pairtree consistently beats other methods (Section 9.3.1). Pairtree produces results for all 576 datasets, which no other method is able to accomplish. Across different numbers of subclones, Pairtree produces lower tree error except in the 100-subclone case, where CALDER generally fares better on the 21% of datasets where it produces a solution. Success rates for LICHeE are lower than in the previous comparison (Fig. 3a) because, for tree error, we rejected solutions that did not include the full set of mutations. By comparison, when computing relationship error, we placed all excluded mutations from LICHeE’s solution set in the “garbage” relationship with respect to other mutations.

### 9.13 Examining method performance on low-depth datasets

The full set of simulated data includes datasets with read depths of 50, 200, and 1000 (Section 9.8), while the B-ALL data obtained via WES on which Pairtree was evaluated has a median read depth of 212 (Section 2.9). The read depth of 50 is meant to reflect typical WGS settings, while the higher depths both on real and simulated data reflect WES or targeted sequencing. To evaluate how well methods perform in typical WGS scenarios, we examined performance on only the subset of cases with a 50x read depth. No method consistently succeeds or fails more frequently on this subset of datasets relative to the full simulation set (Fig. S6). As on the full set, Pairtree is the only method able to produce results on every simulated dataset, and LICHeE and PhyloWGS run successfully on all datasets with 30 subclones or fewer. PASTRI produces results for only one dataset with 10 subclones or more, while CITUP fails on all such datasets. CALDER does not produce results on all datasets in any of the 3, 10, 30, or 100 subclone subsets with low depth, and produces results for only 8% of the 100-subclone cases.

VAF losses in the low-depth setting are consistently lower than when all simulated datasets are considered (Fig. S6). This is to be expected, as an inaccurate lineage frequency will impose a lower cost given fewer total reads for a mutation. Notably, Pairtree continues to perform well on low-depth datasets, with its VAF loss never exceeding 0.007 bits on trees with 30 or fewer subclones. Relationship errors are almost unchanged between settings, both in the rankings of different methods and the magnitude of the errors they produce (Fig. S7). As with the full set of simulations, Pairtree performs less favorably on 100-subclone trees, where its VAF loss and relationship error are on par with or slightly worse than CALDER and PhyloWGS. Again, however, PhyloWGS produces results for only 58% of 100-subclone datasets and CALDER for 8%, while Pairtree yields results on all.

### 9.14 Why existing algorithms failed

Given that the algorithms we compared against often failed to produce results on our simulated datasets, considering possible reasons for this poor performance is a worthwhile exercise. When building trees with few subpopulations, exhaustive enumeration algorithms are attractive, as they promise to find the single best tree by considering all possibilities. As our simulations demonstrated, however, enumeration algorithms cannot cope with more than ten subpopulations, as the number of possible trees becomes too great, even when constraints are employed to reduce possible tree configurations. Stochastic search algorithms are a superior approach when faced with numerous subpopulations, provided they can locate high-likelihood regions of tree space and limit their search to those areas. When this space is searched blindly, however, it remains difficult to navigate, given the massive number of possible clone trees formed from having many subpopulations.

We hypothesize that CITUP attempted to enumerate all trees with a given number of subpopulations, but faced too many trees to make this approach feasible when provided with more than three subpopulations. Thus, CITUP is limited to datasets with only a small number of subclones.

PASTRI attempted to overcome the difficulties of enumeration by first sampling subclonal frequencies, then enumerating only trees consistent with those frequencies. Because mutation VAFs are independent from the tree when conditioned upon the subclonal frequencies, PASTRI can treat its approximate posterior over subclonal frequencies as a proposal distribution for importance sampling, where the target is the posterior distribution over subclonal frequencies permitted by the true tree. The PASTRI implementation is nevertheless limited to 15 subpopulations [35]. Even with ten subpopulations or fewer, because PASTRI samples frequencies without considering tree structure, the frequencies are often inconsistent with any tree when the algorithm is given many cancer samples, as the frequencies collectively impose constraints that rule out all possible trees. A weakness of this approach becomes apparent in real cancer datasets, where new subpopulations often emerge when they acquire driver mutations that confer a strong selective advantage, leading to them displacing their parents such that the subclonal frequency of the child is only slightly greater than that of the parent. Indeed, this situation often occurs in the leukemias considered here. As PASTRI samples subclonal frequencies before enumerating consistent trees, the frequencies sampled for children in this situation will often by chance be slightly higher than their parent, rendering the correct tree structure impossible to recover.

LICHeE fared better than CITUP and PASTRI, as it first constructed a directed acyclic graph (DAG) containing possible trees permitted by the noisy subclonal frequency estimates provided by the VAFs, then only considered spanning trees of this graph [22]. However, this approach could not scale to most 100-subpopulation trees, presumably because the corresponding DAGs have too many spanning trees. Even in settings with 30 or fewer subclones, LICHeE exhibited considerably higher error than Pairtree both with respect to subclonal frequencies and pairwise relations, despite us computing subclonal frequencies for LICHeE’s tree structures using the same algorithm as Pairtree. This suggests that the DAGs did not include as spanning trees good tree candidates, or that the error scoring function LICHeE used to indicate tree quality did not properly reflect tree quality. Some of LICHeE’s shortcomings may have arisen because it takes as input only VAFs, rather than mutation read counts. Consequently, LICHeE has no knowledge of how precisely the VAFs should reflect underlying subclonal frequencies, unlike methods such as Pairtree that use a binomial observation model.

When PhyloWGS fared poorly, its performance could often be attributed to its inability to use a fixed clustering, unlike the other methods. Because we gave PhyloWGS supervariants rather than individual mutations in an attempt to mitigate this discrepancy, even though PhyloWGS could not split clusters into multiple subclones, the algorithm could effectively merge distinct subclones into single entities, causing considerable pairwise relationship error.

Given that non-Pairtree methods may have been particularly prone to failing on the most challenging simulations, summary statistics reported for these methods may be unfairly biased in their favour, as they would only reflect performance on less-challenging datasets. Nevertheless, when we compare Pairtree to each method on only the subset of datasets for which the comparison method succeeded (Fig. S3), we see that Pairtree almost always produces better VAF losses on the 468 datasets with 30 or fewer subclones. The only exception are two datasets with three subclones, where CALDER achieves infinitesimally lower VAF losses, such that Pairtree is within 0.002 bits of CALDER’s solution. PhyloWGS has better VAF loss than Pairtree on 16 of the 108 datasets with 100 subclones, but fails on 38% of these datasets. When CALDER can produce a result on 100-subclone datasets, it generally does better than Pairtree, producing lower VAF losses on 22 of the 23 such datasets where it runs. However, CALDER fails to produce a result on the other 85 datasets with 100 subclones, while Pairtree succeeds on all.

In general, stochastic search algorithms are a superior approach relative to exhaustive enumeration methods when faced with numerous subpopulations, since they avoid the exponential growth in number of trees as a function of number of subclones [23]. For stochastic search algorithms to work well, they must locate high-likelihood regions of tree space and limit their search to those areas. However, as data become richer, tree space is rendered more complex, such that existing search algorithms struggle to navigate through it. This was apparent with PhyloWGS, which consistently exhibited higher error for many-cancer-sample simulations than few-cancer-samples ones. By constructing the pairs tensor and using this as a guide to tree search, Pairtree is better able to cope with many cancer samples and the constraints they impose.

### 9.15 Comparing the computational costs of methods

#### 9.15.1 Criteria for measuring computational costs

Pairtree and the four methods we compared to it differed substantially in the computational costs they imposed, as well as their ability to conduct computations in parallel using multiple CPU cores, using either multiple processes or multiple threads. Pairtree, CITUP, and PhyloWGS had the ability to conduct computations in parallel, while LICHeE and PASTRI did not. We used this ability only for Pairtree, however. For CITUP, using the method’s multiple-process mode did not improve its failure rate. Though PhyloWGS allows running multiple MCMC chains in parallel, doing so was not helpful for this study—PhyloWGS’ failures stemmed from an inability to sample enough trees to form a posterior estimate in 24 hours from a single chain, and so increasing the number of chains only amplified the computational burden without improving the failure rate.

We measured runtime on each simulated dataset for each method both with respect to CPU time and wall-clock time. CPU time indicates the number of CPU seconds consumed by a method’s primary process and any subprocesses or threads it spawned, in either user or kernel mode. Wall-clock time measures the elapsed time a method took. Runs that exited with an error without producing a result, or that failed to finish in 24 hours of wall-clock time, are excluded from the results. Thus, the maximum wall-clock time observed for any method is 86,400 seconds. Considering both CPU time and wall-clock time is worthwhile—CPU time reflects the total computational burden imposed by a method, while wall-clock time indicates how long a method will take to finish in a multi-CPU environment. We conducted all experiments on compute nodes using dual Intel Xeon Gold 6148 CPUs, such that 40 CPU cores were available to each method. On systems with only one CPU, we expect that wall-clock time will generally be slightly more than CPU time, as that single CPU must also be used for the operating system and other concurrent tasks. In our experiments, however, non-Pairtree methods that used only a single CPU core for a run typically achieved wall times that were less than CPU times, given that system or library calls they made (e.g., to numerical routines in the Python library NumPy) could be parallelized.

#### 9.15.2 Examining method runtime

In cases with 3, 10, or 30 subclones, we see similar patterns of CPU time consumed for Pairtree, LICHeE, and PhyloWGS (Fig. S10). These three methods succeeded on all simulations with 30 or fewer subclones, simplifying comparisons. Across datasets with 3, 10, or 30 subclones, LICHeE was fastest, realizing median CPU times of 0.46 seconds, 1.6 seconds, and 2,722 seconds, respectively. This characterization is unfair to other methods, however, as LICHeE did not compute subclonal frequencies for the tree structures it produced. To overcome this deficiency, we invoked Pairtree to compute subclonal frequencies for LICHeE’s results, but did not include the time this step took in LICHeE’s CPU time or wall-clock time measurements. Pairtree was slower than LICHeE, taking median times of 993 seconds, 1506 seconds, and 4391 seconds in settings with 3, 10, or 30 subclones, respectively. PhyloWGS was faster than Pairtree for 3-subclone cases, needing only a median CPU time of 509 seconds, but slower in 10- and 30-subclone cases, taking median times of 1,781 and 35,472 seconds. When we compare each method’s CPU time to Pairtree’s on only the subset of datasets for which each method succeeded, these observations are reinforced, with LICHeE usually being faster than Pairtree excepted for outliers corresponding to 100-subclone cases, and PhyloWGS usually being slower than Pairtree (Fig. S12). As CITUP could not produce results for datasets with more than three subclones, and PASTRI failed on most three- and ten-subclone cases, we do not consider their performance in depth, except to note that CITUP and PASTRI are generally fast when they can produce results for three-subclone cases, while PASTRI is slower than all other methods on the 4% of 10-subclone datasets where it ran successfully (Fig. S10).

When examining wall-clock times, however, we see that Pairtree fares better because of its use of multiple CPU cores. In few-subclone cases, Pairtree is still slower than LICHeE, with Pairtree taking median wall times of 55 seconds and 69 seconds in the 3- and 10-subclone settings, respectively, while LICHeE took 0.326 and 0.93 seconds, respectively (Fig. S11). Conversely, Pairtree is faster than LICHeE in settings with more subclones. For 30-subclone datasets, Pairtree takes a median 148 seconds, while LICHeE takes 2,685 seconds. PhyloWGS was considerably slower with respect to wall-clock time than LICHeE and Pairtree across all three settings. When runtime on individual datasets is examined, Pairtree demonstrates a comparable or superior wall-clock time relative to PhyloWGS and LICHeE (Fig. S13).

Datasets with 100 subclones warrant separate consideration. Pairtree took a median 23,827 seconds of CPU time on 100-subclone cases (Fig. S10), but only a median 675 seconds of wall-clock time (Fig. S11). LICHeE produced results for only 8% of these datasets, where it took a median 74,790 seconds of CPU time. PhyloWGS yielded output for 62% of such datasets, taking median times of 86,400 seconds for both CPU time and wall-clock time. The method’s median times being equal to 24 hours reflects how we handled incomplete runs. According to the (default) parameter settings used for these experiments, PhyloWGS discards the first 1000 samples from its MCMC chain as burn-in samples not reflective of the true posterior, then takes an additional 2500 posterior samples. If the method finished the 1000 burn-in samples within the 24-hour wall-clock period permitted, but completed fewer than the 2500 posterior samples, we used whatever partial set of posterior samples the algorithm produced to evaluate its accuracy, while recording its runtime as 24 hours. The median times being 24 hours indicate that most successful 100-subclone runs fell into this category. Conversely, the 68% of 100-subclone cases where we recorded no output correspond to instances where PhyloWGS could not finish its initial 1000 burn-in samples.

CALDER was generally the least costly method on datasets with 30 subclones or fewer both with respect to CPU time (Fig. S10) and wall time (Fig. S11), on the subset of such datasets where it could produce a result. On datasets with 100 subclones, it was often the most costly with respect to wall time (Fig. S13), and consumed vastly more CPU time than any other method on the fraction of 100-subclone cases where it produced a result (Fig. S12).

#### 9.15.3 Evaluating the performance costs of Pairtree’s two stages

The two primary steps composing the Pairtree algorithm are computing pairwise relations between subclones and searching for trees. Tree search includes computing MAP subclonal frequencies for each tree structure. The amount of computation needed to build the pairs tensor is fixed, as a distribution over relations for every pair must be computed regardless of how many CPU cores are available. As relations for each subclone pair are independent of all other subclones, the pairwise computations are embarrassingly parallel, such that they can be trivially computed in parallel for all pairs. Thus, though the total computational burden represented by CPU time is constant, the wall-clock time can be greatly reduced by using more CPU cores, with N cores reducing the time needed for this stage nearly by a factor of N. By comparison, tree search requires that each MCMC chain acquire samples serially, such that any one chain cannot be parallelized. Multiple chains, however, can execute in parallel, increasing CPU time consumed in proportion to the number of chains, but with little effect on wall-clock time.

In the Pairtree experiments illustrated throughout this paper, we used all available 40 CPU cores on our compute nodes to calculate pairwise relations in parallel, and to run 40 parallel MCMC chains for tree search. Doing so greatly inflated CPU time relative to wall-clock time, but likely was not necessary to realize good results. Results of nearly equal quality could perhaps have been obtained from Pairtree using fewer chains—while any one chain may become mired in pathological regions of tree space corresponding to a local optimum, such that multiple chains initialized from different positions can yield better samples, we likely did not need all 40 chains to realize this benefit. Nevertheless, even if all 40 chains were necessary to produce results of this quality, running those chains serially on a single CPU would have been feasible. In this case, the wall-clock time would have been approximately equal to the CPU time. Amongst the 576 simulations, Pairtree’s longest run was on a 100-subclone, 100-cancer-sample dataset that took 1,110 seconds of wall-clock time (Fig. S11) and 36,606 seconds of CPU time (Fig. S10). Running all 40 chains serially on a single CPU would thus have resulted in a wall-clock time of slightly over 10 hours.

We can understand the relative computational costs of Pairtree’s two primary steps by comparing the runtimes of the full Pairtree algorithm to the portion that computes the pairwise relations, denoted as *pairs tensor*. By subtracting the pairs tensor runtime from that of full Pairtree, we reveal the cost of tree search alone. Comparisons are most informative for the 100-subclone, 100-cancer-sample datasets, where the runtimes are longest and differences are thus clearest. For instance, the single most costly Pairtree run took 1,110 seconds of wall-clock time and 36,606 seconds of CPU time, as above (Figs. S10 and S11). Computing the pairs tensor alone took 81 seconds of wall-clock time and 2,666 seconds of CPU time. Whether we consider CPU times or wall-clock times, we see 7% of Pairtree’s time went to computing pairwise relations, while 93% went to tree search. If the number of CPU cores dedicated to this run were cut tenfold to four CPUs rather than 40, we would expect the wall-clock cost of computing pairwise relations to increase proportionally to 810 seconds, while the CPU time would remain constant. Conversely, the wall-clock cost of tree search could be kept constant at 1,110 seconds by reducing the number of MCMC chains to four, at a potential cost in result quality. In this instance, we would expect Pairtree to take 810 + 1, 110 = 1, 920 seconds, with tree search consuming 58% of the total. Thus, the relative burdens of computing the pairs tensor and performing tree search depend both on the number of CPU cores used in parallel, and on the number of MCMC chains from which the user elects to sample trees.

### 9.16 Multiple trees are often consistent with observed data, which Pairtree can accurately characterize

When building trees, algorithms draw on the subclonal frequencies of constituent subclones across cancer samples and relationships between these frequencies to determine possible tree structures. Thus, to assess method performance on simulated data, we can enumerate all tree structures consistent with the true subclonal frequencies used to generate the data, yielding a distribution over trees. This distribution will include the true tree used to generate the data, as well as any other tree structures that are also consistent with the subclonal frequencies. A perfect method would be able to recover this distribution exactly, despite being given only noisy estimates of the true subclonal frequencies via the observed mutation frequencies. To evaluate a method, we can then determine the extent to which its tree distribution matches the true distribution of all trees consistent with the true subclonal frequencies.

Amongst our 576 simulated datasets, if only one cancer sample is provided, there are usually multiple trees consistent with the data (Fig. S14a), regardless of how many subclones are in the tree. This reaches an extreme in our ten-subclone, single-sample simulations. This illustrates the importance of understanding uncertainty in these reconstructions, rather than simply producing a single answer (Section 2.10)—the perfect method should represent all of these trees as being equally consistent with the data, such that the user should have no reason to prefer any one structure over the others. Drawing on more cancer samples reduces this uncertainty, with most ten-sample datasets possessing only a single possible tree across the three-, ten-, and 30-subclone settings (Fig. S14a). With 100 subclones, ten samples still permits little uncertainty, with the number of possible trees rarely exceeding ten. Note, however, that in this simulated setting, multiple samples are likely to be more powerful than they would be for real cancers. Here, each sample had its subclonal frequencies generated independently from other samples, increasing the chance that the sample induces tree structure constraints because its frequencies are different from all other samples. In reality, samples are likely to have correlated frequencies, given that they may be taken from similar spatial or temporal sites in the cancer that have similar population proportions.

By computing the entropy of tree distributions, we can characterize how many high-confidence trees exist in the distribution. Effectively, the entropy is a posterior-weighted count of the number of trees, with the weights in the true tree distribution being uniform because all solutions are equally consistent with the data. To determine how many high-confidence solutions was Pairtree was finding relative to the number of possible solutions, we compared Pairtree’s tree entropy for each simulated dataset to the entropy of the true tree distribution (Fig. S14b). Pairtree’s entropy generally tracked the true entropy well, suggesting that Pairtree’s uncertainty was usually consistent with the uncertainty in the true tree distribution. Notably, in settings where the number of cancer samples was higher than the number of subclones, there was only ever one true tree (Fig. S14a), while Pairtree’s tree distribution entropy exceeded the true distribution’s entropy by more than 5.9 × 10^-6^ bits with only one exception across 181 simulations (Fig. S14b). These results demonstrate that, when the data is sufficiently high-resolution as to permit only a single solution, Pairtree finds only a single solution.

Though examining tree distribution entropies reveals the number of high-confidence trees Pairtree finds, it says nothing about the quality of those trees. To gain further insight, we can view a distribution over trees as inducing a distribution over the *parents* of each subclone. For a given dataset, to compare the Pairtree-computed tree distribution to the distribution of trees consistent with the true subclonal frequencies, we can consider the joint Jensen-Shannon divergence between parent distributions induced by these tree distributions, normalized to the number of subclones in the tree such that the divergence will always lie between zero bits and one bit. We refer to this metric as the *parent JSD*. Even if the tree distributions have no overlap—which could occur, for instance, if there is only a single true tree that Pairtree fails to locate—the parent JSD nevertheless allows the distributions to have a small divergence if they agree on parent choice for most subclones. We see that the parent JSD falls as the number of samples increases for a given number of subclones (Fig. S14c), suggesting that Pairtree can efficiently exploit the constraints provided by additional cancer samples to produce higher-quality trees. Moreover, when the number of samples exceeds the number of subclones such that there is only one tree consistent with the true subclonal frequencies (Fig. S14a), the parent JSD is effectively always zero, complementing the tree entropy analysis (Fig. S14b) to show that the one tree Pairtree finds is almost perfectly consistent with the true tree. Additionally, when the pairwise relation error is examined at a more granular level (Fig. S14d), we see that for a given number of subclones and samples it is always less than the parent JSD. This suggests that, even when Pairtree doesn’t perfectly determine the parents of each subclone, the distributions over relationships between subclones (e.g., ancestor-descendant or on-different-branches) are closer to the truth. The quality difference between pairwise relation distributions and parent distributions is stark for the 100-subclone setting. Though Pairtree only rarely finds the correct parents, demonstrated by the parent JSDs that are close to one (Fig. S14c), the pairwise relation errors are much lower (Fig. S14d), indicating that the higher-level relationships between subclones are closer to being correct.

### 9.17 Characteristics of simulated data

#### 9.17.1 Trees are dominated by small subclones

Examining statistics of simulated data illustrates factors that affect each clone-tree-reconstruction algorithm’s ability to recover good solutions. The nodes of each clone tree correspond to populations, with subclones consisting of sub-trees made up of a population and all its descendants (Section 2.1). Thus, a tree with *K* populations defines *K* subclones. Subclones are nested within trees—a subclone with population *i* at its head and *c* total populations is also part of a subclone with *i*’s parent at its head and *c* +1 total populations (excluding the root subclone that corresponds to the entire tree, which has no parental subclone). Characterizing subclone composition within simulated data is helpful, as several properties of the simulated trees depend on how many populations compose each subclone.

A fully linear tree with no branching that contains *K* populations would yield a uniform distribution over subclones consisting of 1, 2, …, *K* populations, with exactly one subclone of each size. Branching within trees depletes the contribution of larger subclones, replacing them with smaller ones. Because of how we constructed simulated tree structures (Section 9.8.2), we see that small subclones dominate regardless of the number of populations within a tree (Fig. S15), with most subclones consisting of ten or fewer populations in the 30- or 100-subclone trees. In the tree generation algorithm, we choose parents for each population in turn, selecting the preceding population as parent with 75% probability, and otherwise choosing a parent uniformly from the other nodes already in the tree. As a result, the length of linear chains of populations within the tree roughly follows a geometric distribution. Linear chain length deviates from the distribution, however, because a node may choose as its parent the end of a different chain, allowing that chain to continue extending under a new geometric process.

#### 9.17.2 Tree construction becomes increasingly difficult with more subclones

Large trees containing many subclones are more difficult to reconstruct than small trees. In part, this is because the number of possible tree structures scales exponentially with the number of populations [23]. We must also consider, however, how relationships between subclones become more difficult to infer as the number of subclones grows, which is a factor independent of tree structure. Given how we generated the simulated data (Section 9.8.2), we can derive statistics of the simulated data, then use them to show how the difficulty of inferring relationships between subclones changes according to the numbers of subclones and cancer samples.

In determining the proper placement of a population within a clone tree, two properties related to population frequencies affect the difficulty of this task. Firstly, if a population *k* has a near-zero population frequency *η_ks_* in a cancer sample *s*, the VAFs associated with its mutations in that sample will be difficult to distinguish from the VAFs of mutations in *k*’s parent, which we will denote as population *j*. This occurs because the VAFs for mutations that arose in each population are sampled based on the subclonal frequencies of the populations’ subclones (Section 9.8.2), which are computed from the sum of the population frequencies composing the subclone (Section 9.4.1). Thus, when *η_ks_* ≈ 0, we have *ϕ_ks_* ≈ *ϕ_js_*, and the VAFs in *k* and *j* will be nearly the same. Assuming there are no cancer samples other than sample *s*, we could thus swap the positions of *k* and *j* in the tree without affecting tree likelihood—both populations would have nearly the same subclonal frequency fit to them in the tree, which would fit the two sets of VAFs almost equally well. Larger population frequencies avoid this situation, making clearer the proper ordering of parents and children.

Intuitively, as more populations appear in a tree, the *η_ks_* frequencies will become smaller on average, as the unit mass apportioned by the Dirichlet distribution from which the frequencies are drawn must be split amongst more entities. Indeed, by the properties of the Dirichlet distribution, for *K* subpopulations in a sample *s* with [*η*_0*s*_, *η*_1*s*_, …, *η_Ks_*] ~ Dirichlet(*α, α*, …, *α*) (Section 9.8.2), we have 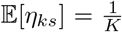. This is evident when we examine the distribution over *η_ks_* frequencies for each population in the simulated trees (Fig. S16A), where the largest frequency observed across cancer samples for each population is typically close to 1 for trees with three subclones, but gets progressively smaller as the number of subclones increases, with populations in 100-subclone trees dominated by small frequencies. To distinguish a population from its parent, it need have a non-negligible *η_ks_* frequency in only one sample *s*, which is part of why adding cancer samples is so helpful in resolving evolutionary relationships between populations, and ultimately reconstructing an accurate clone tree.

The second property related to population frequency that affects the difficulty of clone tree reconstruction is the variance over cancer samples *s* in a subclone *k*’s frequencies *ϕ_ks_*. Suppose you are trying to resolve the position of two subclones *A* and *B* in a tree, using the frequencies in cancer samples *s* and *s*′. To gain the greatest benefit from having two samples rather than only one, we want there to be as much variance as possible in the subclonal frequencies between samples. The power of multiple samples comes from these differences—for instance, if *ϕ_As_* > *ϕ_Bs_*, but *ϕ_As′_* < *ϕ_Bs′_*, we conclude that *A* cannot be the ancestor of *B*, and *B* cannot be the ancestor of *A*, since an ancestral subclone must have a frequency at least as high as its descendants across every cancer sample. This is termed the *crossing rule* [40], and leads to the conclusion that *A* and *B* must occur on separate tree branches. Unfortunately, as we observe only a noisy estimate of the subclonal frequencies through the VAFs, if the subclonal frequencies for *A* and *B* are nearly the same in both samples, the noise in VAFs can obscure this relationship. The less variance there is between *ϕ_As_* and *ϕ_As′_*, and between *ϕ_Bs_* and *ϕ_Bs′_*, the more likely that max(|*ϕ_As_* – *ϕ_Bs_*|, |*ϕ_As′_* – *ϕ_Bs′_*| < *ϵ* for some near-zero *ϵ*, and the more difficult it will be to utilize the crossing rule with our noisy observations.

Suppose we have a subclone *C* composed of |*C*| ≤ *K* populations, such that *C* ⊆ {0,1, …, *K*}. As before, given cancer sample *s*, we have population frequencies [*η*_0*s*_, *η*_1*s*_, …, *η_Ks_*] ~ Dirichlet(*α, α*, …, *α*) (Section 9.8.2), and *ϕ_Cs_* = ∑_*i*∈*C*_ *η_is_*. By the properties of the Dirichlet distribution, we know that the sum of Dirichlet-distributed variables is itself Dirichlet-distributed, such that

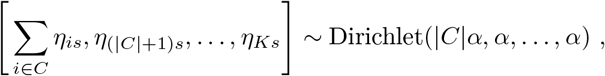

where the first element of the vector represents the subclonal frequency ∑_*i*∈*C*_ *η_is_* = *ϕ_Cs_*, and the final *K* – |*C*| elements represent the population frequencies of all populations not in subclone *C*. From this, we get

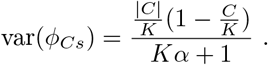

From the denominator, we see that variance is reduced either with more populations *K*, or with a larger Dirichlet parameter *α*. By plotting both the (theoretical) population standard deviation and (empirical) sample standard deviation (Fig. S16B), we see that the latter conforms to the former, and that variance is maximized for subclones with 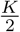 populations, conferring the greatest benefit from multiple cancer samples to populations near the root of the tree, such that they have half the populations as descendants. Conversely, subclones with less variance in frequency across samples will either be at the very top of the tree, with almost all populations as descendants, or at the bottom of the tree, with few populations as descendants. Note that, in Fig. S16, the sample standard deviation appears less than the population standard deviation, particularly in the three- and ten-subclone cases. This effect is exaggerated for those settings because they include single-sample datasets with zero sample standard deviation, whereas the 30- and 100-subclone datasets do not.

#### 9.17.3 Simulated data often include subclones that are impossible to resolve

If a population *k* has a near-zero population frequency *η_ks_* across all cancer samples *s*, its position in a clone tree relative to its parent *j* is difficult or impossible to resolve. Since *k*’s subclonal frequency *ϕ_ks_* is equal to the sum of the population frequencies of all populations in the subclone, when *η_ks_* ≈ 0, we have *ϕ_ks_* ≈ *ϕ_js_*. When this occurs, we will have two candidate trees that fit the data equally well—one in which *k* is the parent of *j*, and one in which *j* is the parent of *k*. Both tree structures would permit tree-constrained subclonal frequencies that fit the observed VAF data almost equally well. Well-behaved algorithms should find both tree structures. Thus, populations whose frequencies are negligible across all cancer samples lead to their subclonal frequencies being nearly equal across all cancer samples, which leads to ambiguity. In real data, we are unlikely to be faced with this situation. The observed VAFs for two variants serve as noisy estimates of their subclones’ subclonal frequencies. When the observation noise exceeds the negligible differences in the subclonal frequencies, we will deem the two variants as having originated from the same subclone, such that the variants are placed in a single cluster.

Nevertheless, examining how often this situation occurs in simulated data is worthwhile, as it grants insight into how well algorithms deal with ambiguity. Note that noisy observations of near-zero population frequencies are not the only source of ambiguity—ambiguity can exist even given noise-free frequencies, or with large population frequencies. All cases where tree enumeration using the noise-free subclonal frequencies found multiple trees (Section 9.9.5) are demonstrations of this alternative ambiguity. Tree-reconstruction algorithms should be able to deal with both sources of ambiguity by finding the full range of solutions permitted for a dataset. With respect to our evaluation metrics, VAF loss (Section 2.4) does not capture algorithms’ performance in this respect, since it penalizes discrepancies between VAFs and tree-constrained subclonal frequencies, and so algorithms can do well regardless of whether they find a single good solution or multiple equivalent solutions. Relationship reconstruction error (Section 2.4), however, properly reflects algorithms’ performance in the face of ambiguity—in the example above in which subclones *j* and *k* had nearly equal subclonal frequencies across all cancer samples, the solutions recovered by a tree-reconstruction algorithm should show both that *k* could be an ancestor of *j*, and *j* could be an ancestor of *k*.

To understand the role near-zero population frequencies play in introducing ambiguity, we must first define a threshold *ϵ* on population frequencies, such that we will say a population frequency *η* is near-zero if *η* < *ϵ*. This *ϵ* should ideally be defined as a function of read depth, since depth determines how precisely the observed VAFs reflect the underlying subclonal frequencies, and ultimately how small population frequencies can get before they are swamped by noise. To set this threshold, consider a fixed read depth of *D* = 200, such that with *V* variant reads and *R* reference reads we have *D* = *V* + *R* = 200. By our simulation framework, we have *V* ~ Binom(*D, ωϕ*), yielding [*E*](*V*) = *ωϕD*. We will define a non-negligible population frequency as that which produces a difference of one read in the mean read counts. While this is a subtle difference, we must remember that, in tree search, the read counts for all variants belonging to a cluster will be summed, exaggerating the difference in observations for the two clusters. Thus, for populations *j* and *k*, we will assume we have subclonal frequencies *ϕ_j_* and *ϕ_k_* with *ϕ_j_* > *ϕ_k_*. Moreover, assume *j* is the parent of *k*, such that *ϕ_j_* = *ϕ_k_* + *η_j_*. This gives us

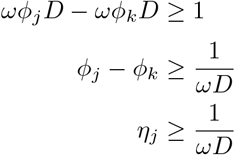

With 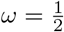, this results in a non-negligible population frequency of *η_j_* ≥ 0.01 for read depth *D* = 200. Conversely, we will define a near-zero population frequency as the complement of this, resulting in a threshold *ϵ* = 0.01. To simplify the analysis, we will use this threshold regardless of read depth. With read depths *D* ∈ {50,200,1000} (Section 9.8.2), this choice of *ϵ* will yield a greater difference in binomial mean for *D* = 1000, and a smaller difference for *D* = 50. Nevertheless, the conclusions we reach for fixed *ϵ* will be broadly applicable regardless of read depth.

First, we will consider how many populations within each simulated dataset have population frequencies less than *ϵ* = 0.01 across all cancer samples *s*. Let *η_ks_* denote the population frequency of population *k* in cancer sample *s*. For *K* subpopulations, we have [*η*_0*s*_, *η*_1*s*_, …, *η_Ks_*] ~ Dirichlet(*α, α*, …, *α*). By the properties of the Dirichlet distribution, we have

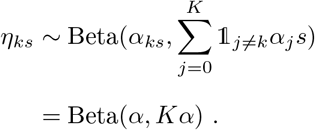

Consequently, since each cancer sample’s population frequencies are independent of every other, for *S* cancer samples we get

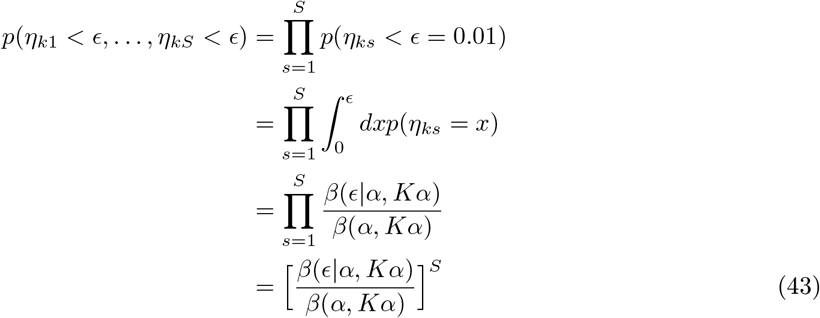

Here, *β*(*ϵ*|*α, *K* α*) refers to the incomplete beta function, and *β*(*α, K α*) refers to the complete beta function. Empirically, the proportion of simulated populations with near-zero population frequencies across samples agrees with the result predicted above (Fig. S17). Datasets with 30 or 100 populations and one or three cancer samples would have at least 38% of populations with near-zero population frequencies in all cancer samples, rendering their positions in the tree difficult to resolve. This would create excessive ambiguity, which is why we did not include such datasets in our simulated data.

The relationship reconstruction error we used to evaluate method performance on simulated data reflected how algorithms dealt with two sources of ambiguity: firstly, the multiple tree structures potentially permitted by the noise-free frequencies (Section 9.16); and, secondly, the additional tree structures permitted by making noisy observations of populations with near-zero population frequencies. For the second factor, as we established above, if a population *k* has near-zero population frequencies across all cancer samples, the subclonal frequencies of *k* and its true parent *j* will be almost equal, such that the noisy VAF observations will render difficult the task of determining whether *j* is the parent of *k* or vice versa. Observe that 14% of populations in 100-subclone, 10-sample trees have noise-free population frequencies less than *ϵ* = 0.01 across cancer samples. In the average tree, these would correspond to 14 populations with near-zero frequencies. Since each such population could be swapped with its parent while minimally affecting tree likelihood, these would generate 2^14^ ≈ 16,000 additional trees. This assumes that none of the populations with near-zero frequencies have edges between them; chains of two or more populations with near-zero frequencies would further increase the number of potential tree configurations. The first factor, by comparison, is less important. In the 100-subclone, 10-sample setting, none of the 36 simulated datasets permitted more than 42 trees given the noise-free frequencies (Fig. S14), which is a value far smaller than the 16,000 trees we expect to be permitted by the noisy observations. Thus, we expect noisy observations to be the dominant source of ambiguity.

This analysis also helps us understand how many cancer samples we must simulate to remove ambiguity in tree search arising from noisy observations for a given number of subclones. Taking our threshold *ϵ* = 0.01, we can ask how many cancer samples we need before *p*(*η*_*k*1_ – *ϵ*, …, *η_kS_* < *ϵ*). By solving for *S* in Eq. (43), we find that we need 24 or more samples before the probability of a population frequency being less than *ϵ* across all samples falls below 1%. This has implications for variant clustering as well, since a population’s variants become distinguishable from other variants by the clustering algorithm only when one or more cancer samples with non-negligible frequencies for the associated population render the VAFs clearly distinct.

To complement the above analysis concerning lone populations, we will also examine the probability of our simulation process created trees containing sub-trees that consist entirely of populations whose frequencies are less than *ϵ* = 0.01. We define a sub-tree to consist of a subset of the full tree’s nodes, as well as all edges between them, ensuring the sub-tree is connected. Thus, a sub-tree can correspond to a subclone (Section 2.1), but is more general in that it may omit parts of the subclone defined by the ancestral population at the root of the sub-tree. For this analysis, we did not conduct an empirical examination of the simulated data, but used only theoretical results derived from the Dirichlet distribution properties. Given a complete tree composed of *K* populations as well as the root node 0, and a sub-tree composed of populations *T* ⊆ {0, 1, …, *K*} with size |*T*|, we have in cancer sample *s* the result

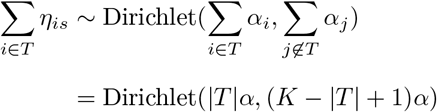

Note that if the sub-tree *T* = {*j*} ∪ {*k*|*k* is descendent of *j*}, then *T* is equivalent to the subclone with population *j* at its head, and ∑_*i*∈*T*_ *η_is_* = *ϕ_js_*. By using the Dirichlet′s marginal beta distribution, as in the previous analysis, we can compute the probability of the arbitrary sub-tree *T* consisting exclusively of populations whose summed frequencies across cancer samples are small, such that ∑_*i*∈*T*_ *η_is_* < *ϵ* = 0.01 for every cancer sample *s* (Fig. S18). For instance, in the 100-subclone, single-sample case, we have a 6% probability of an arbitrary eleven-population sub-tree having a near-zero population frequency sum. With |*T*| populations in such a sub-tree, there are (*T* +1)! orderings of nodes in the sub-tree that would permit nearly equal tree-constrained subclonal frequencies, and thus nearly equal tree likelihood. In the eleven-population case, there would thus be (11 + 1)! = 4.79 × 10^8^ solution trees resulting from this single ambiguous sub-tree.

To compute the probability of observing such a case in the simulated trees, we must first consider how many linear chains of *J* populations exist in a tree with *K* nodes, as each has an equal chance of being assigned these small frequencies. If a tree is fully linear with no branching, there would be (*K* + 1) – *J* + 1 chains of *J* nodes, such that our chain of 11 populations in a 100-subclone tree would have 101 – 11 + 1 = 91 sub-trees, assuming that tree was fully linear. This in turn yields a (100% – 6%)^91^ = 0.36% chance that we would not observe any near-zero-frequency 11-population chains in our tree—i.e., with near certainty, we would encounter such a chain. Any degree of branching in a tree can reduce the number of node chains of a given length, thereby lessening the chance we would see this scenario. Nevertheless, the probability can remain considerable, which is another reason we omitted the many-subclones, few-samples cases from our simulated data. Amongst the settings we included, we see, for instance, that in ten-subclone, single-sample trees, 6% of five-population chains will have small population frequency sums, yielding a 35% chance that we would encounter such a case in a fully linear tree.

#### 9.17.4 Justifying our choice of the Dirichlet parameter for generating simulated data

In Sections 9.17.1 to 9.17.3, we saw that our choice of the Dirichlet parameter α when generating simulated data (Section 9.8.2) affects multiple aspects of simulated data.

1. A smaller *α* leads to more variance in population frequencies between samples, increasing the chance that multiple samples will make clear the proper pairwise relations between subclones.
2. A smaller *α* also leads, however, to a greater probability of observing near-zero frequencies for a population across all cancer samples, inhibiting tree-reconstruction algorithms’ attempts to infer the proper place for such populations in the tree. (We do not present results with alternative α values here, but used these analyses to inform our choice of *α*.)

Our chosen *α* = 0.1 thus achieved a compromise between three factors.

1. It led to sufficient variance in population frequencies between cancer samples for algorithms to benefit from having access to multiple cancer samples.
2. It avoided creating too many populations with near-zero frequencies across samples, which would have created excessive ambiguity.
3. Yet it created enough such populations so that we could evaluate how algorithms dealt with ambiguity stemming from this source.

### 9.18 Using non-uniform priors on tree structure and subpopulation frequency

Pairtree assumes a uniform prior over both tree structure (Section 9.3) and the subpopulation frequencies that determine subclonal frequencies when taken alongside tree structure (Section 9.4). Incorporating non-uniform priors on either or both into the tree-scoring step used during tree search would be straight-forward. However, these priors may make the optimization problem non-convex (Section 9.4.3), which could compromise the accuracy of the Laplace approximation of the marginal likelihood we use. While the current MAP estimate of a tree sample’s subclonal frequencies is proportional to a Laplace approximation of the marginal likelihood (Section 9.3.2), the quality of this approximation (Section 9.4.4) would likely decrease if the posterior over frequencies becomes non-convex.

The nature of selection we expect a cancer to undergo can inform the choice of prior. For example, if strong selection is present, we may observe a linear tree that has been pruned of many branches, and we should expect only a few populations to dominate the composition of each sample. Conversely, if neutral evolution occurs, we should see much more branching in the tree structure, with more populations represented in each cancer sample [14, 54, 55]. Moreover, we expect that a sensible non-uniform prior should depend on the cancer type. For instance, blood cancers should be much more spatially uniform than solid tumours, and should allow for positive selection to more rapidly remove unfit clones.

Other work reveals possible prior choices. For instance, [56, 57] uses driver events to characterize the set of evolutionary trajectories observed in a given cancer type. We could thus use this set as a prior over tree structure, on the assumption that each of our clones contained a corresponding driver mutation that would have favoured one of the previously observed trajectories. Conversely, [43] focuses on building clone trees from longitudinal samples. Given a known time ordering of samples, the method places a prior on tree structures that requires that, once a lineage has gone extinct, it cannot return at a later timepoint. Additionally, the method encourages a sparse representation of subpopulation frequencies by using regularization and a threshold on minimum allowed frequency. This can be viewed as a prior on frequencies that rewards solutions with more populations that go extinct, preventing overfitting of solutions in which extinct populations would maintain a low but nonzero frequency to allow their later return. In Pairtree, we could add an optional time-ordering input for longitudinal studies, then use this to enforce a prior that prevents the return of extinct populations. Additionally, we could structure this prior to reward sparse solutions, just as in [43]. However, if we are able to incorporate a non-uniform prior only when do tree search and not when computing pairwise relations, this could weaken the value of the Pairs Tensor for use as a proposal distribution when making tree modifications.

## 10 Supplementary figures

**Figure S1:**
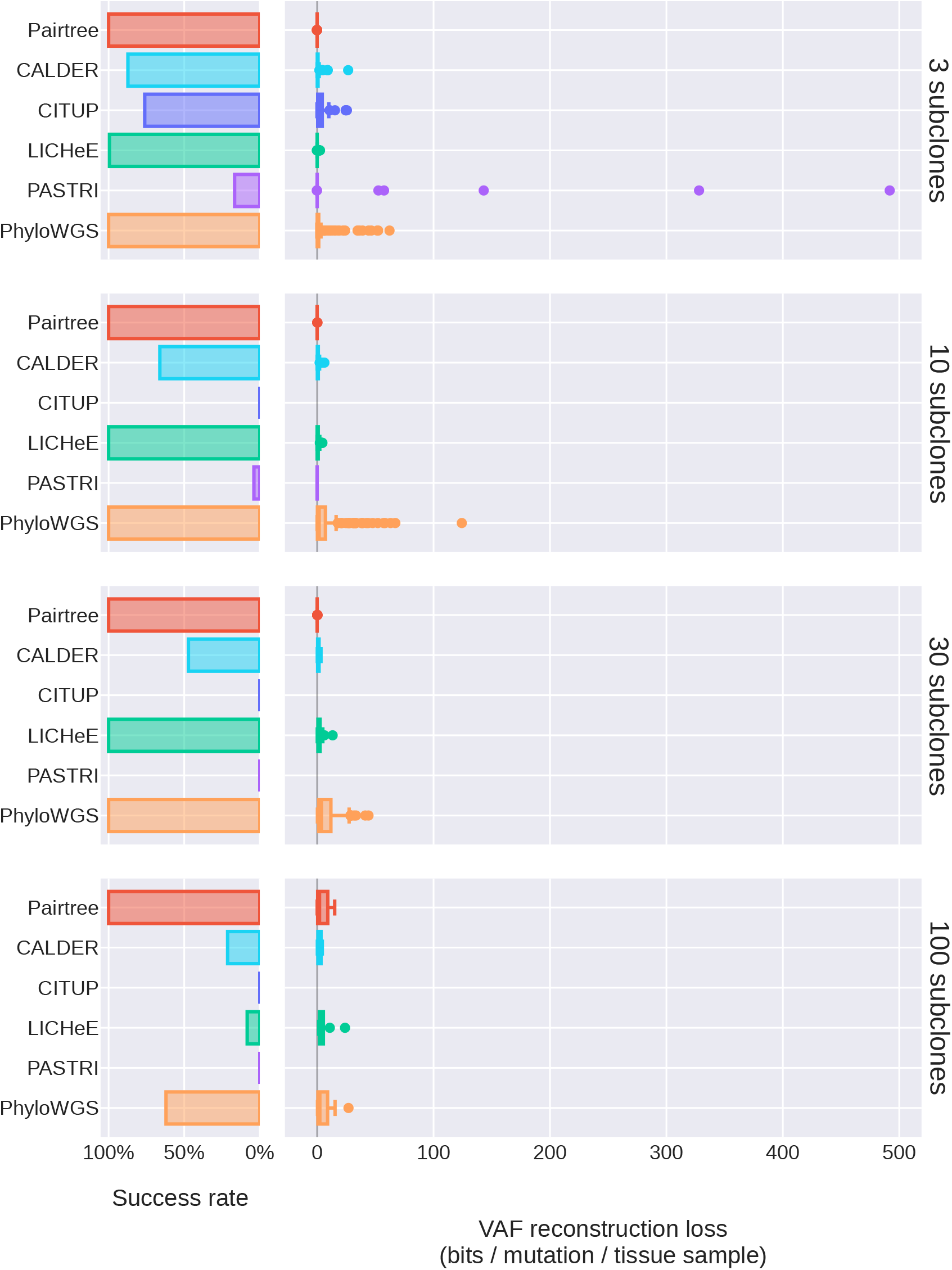
Untruncated VAF reconstruction losses on 576 simulated datasets. These results are the same as in Fig. 3b, but without axis truncation. As in the truncated plots, results reflect each method’s performance on the subset of datasets where it succeeded in running.

**Figure S2:**
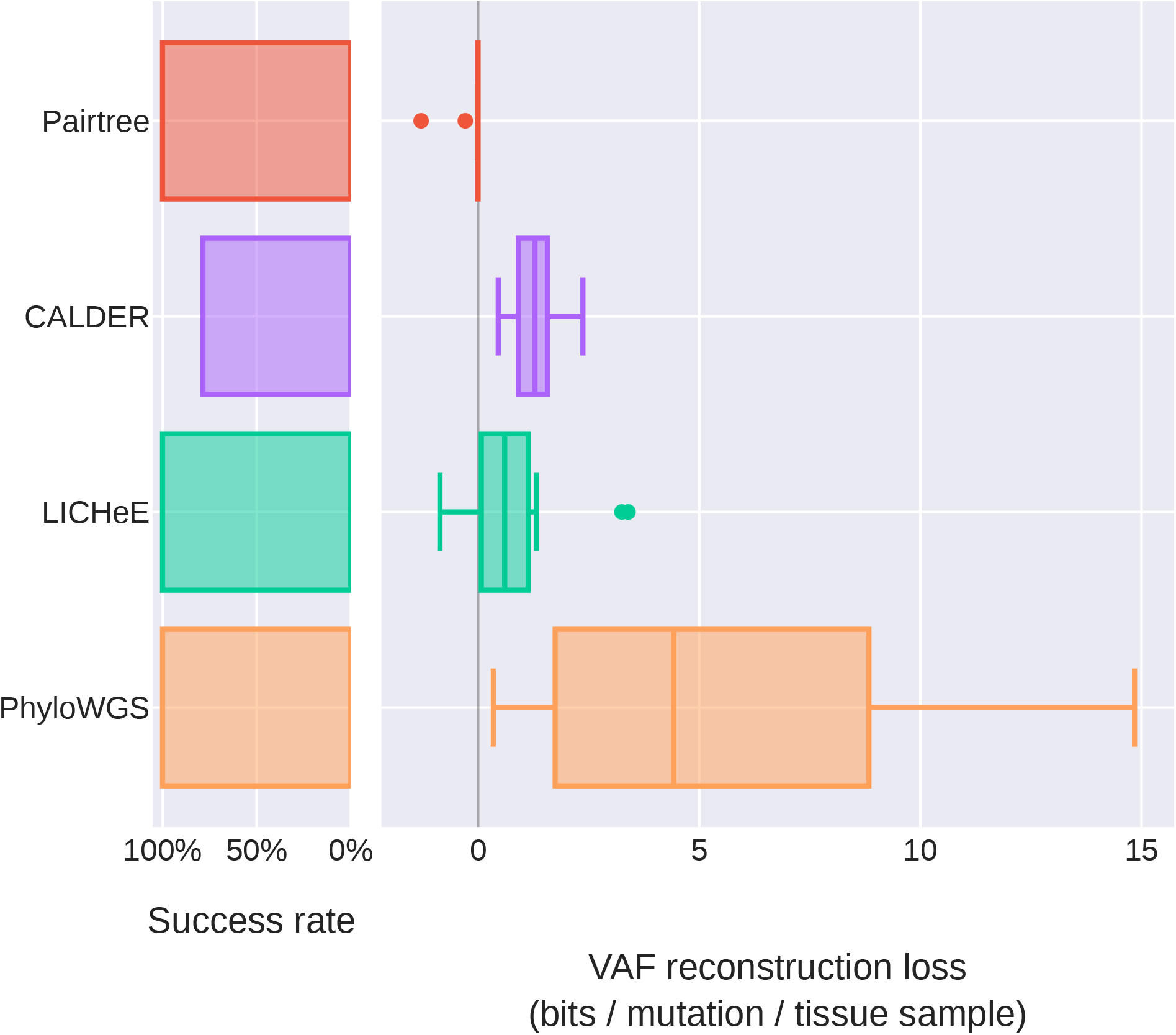
Untruncated VAF reconstruction losses on 14 B-ALL datasets. These results are the same as in Fig. 5, but without axis truncation. As in the truncated plots, results reflect each method’s performance on the subset of datasets where it succeeded in running.

**Figure S3:**
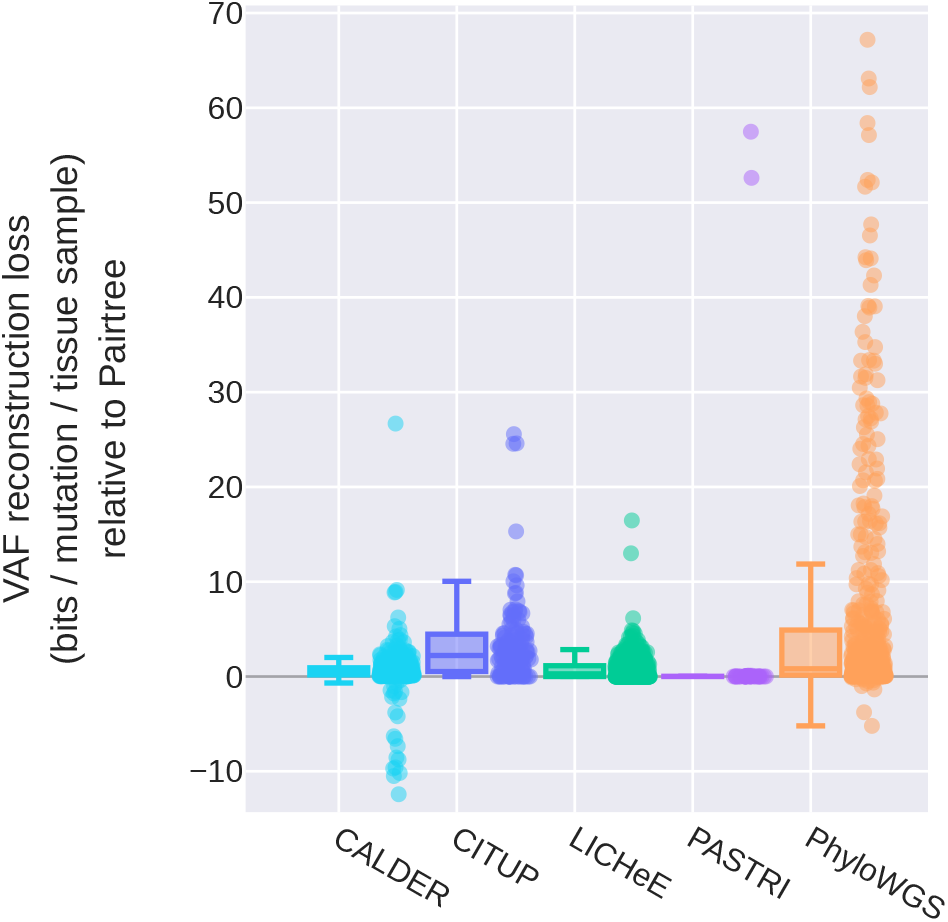
VAF reconstruction loss of each method relative to Pairtree. Each point represents a method’s VAF reconstruction loss on a simulated dataset relative to Pairtree, with positive values indicating worse error. As each method failed on different simulations (Fig. 3a), values are reported only on the subset of 576 datasets where a method produced a result.

**Figure S4:**
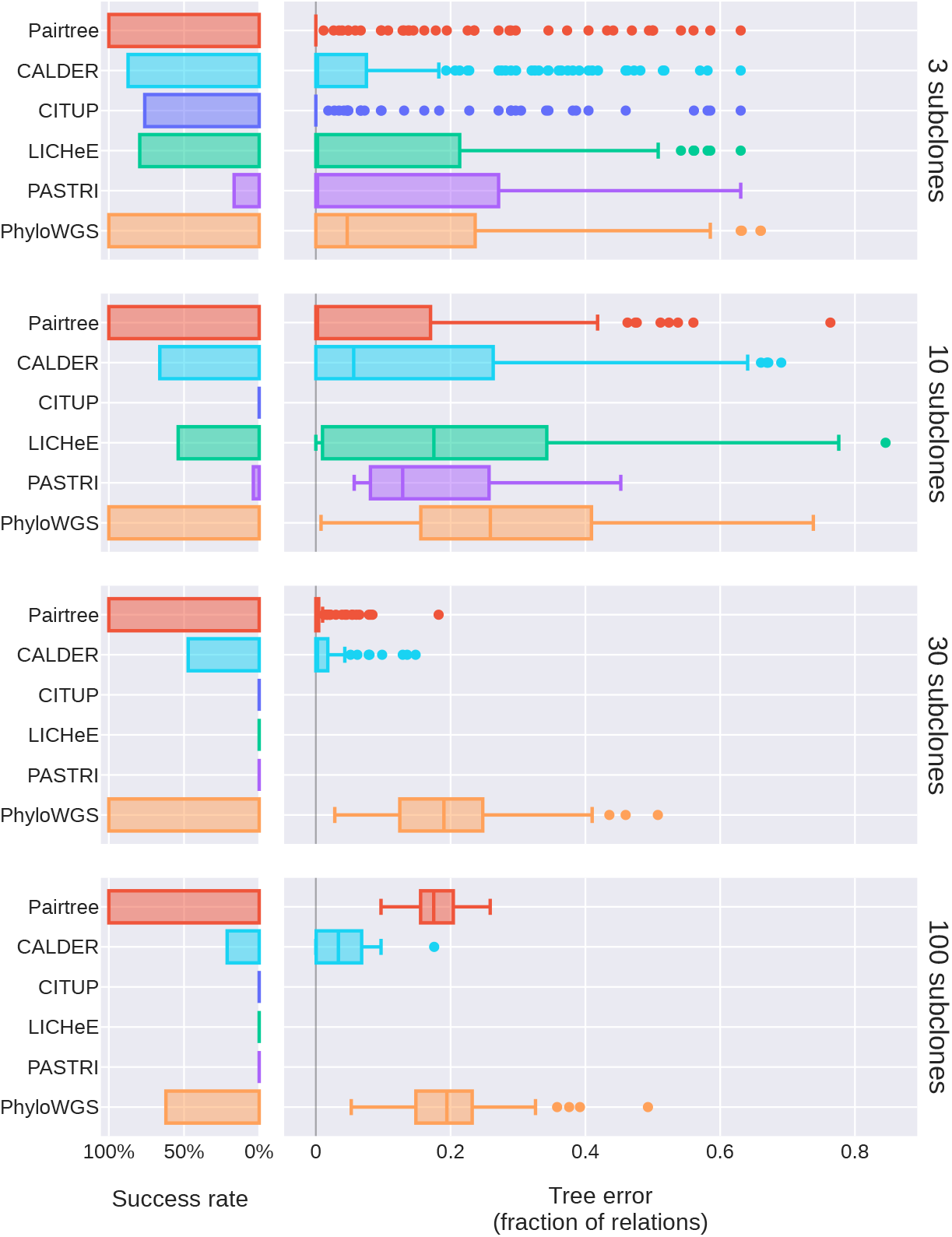
Tree error of each method on 576 simulated datasets. Simulations are grouped by number of subclones (rows). **a.** Bar plots show each method’s success rate in the group. Successes are reconstruction problems for which the method produced at least one tree in 24 hours (wall-clock time) and did not crash. **b.** Boxplots show distributions of tree error for a method on a problem group. Scores reflect only datasets where a method ran successfully. Tree error is the proportion of pairwise relationships in a method’s highest-scored solution that correctly match the relations in the true tree. Unlike relationship error (Fig. 3), tree error does not reflect that multiple trees may be equally consistent with the noiseless lineage frequencies used to generate the tree. Mid-lines in box plots indicate medians.

**Figure S5:**
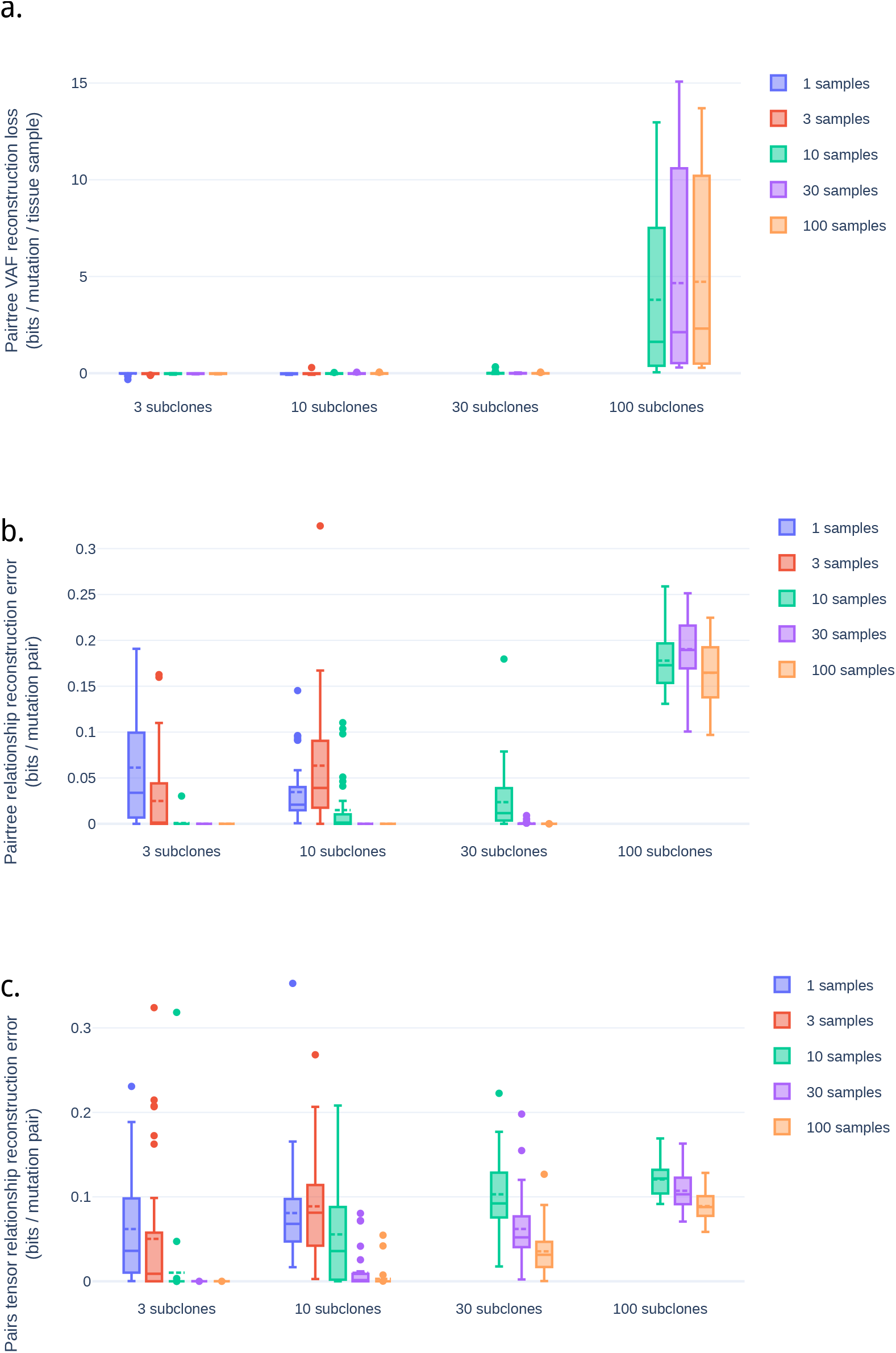
Pairtree’s performance on different numbers of subclones and cancer samples. **a.** Pairtree’s VAF reconstruction loss for each number of subclones and number of cancer samples. **b.** Pairtree’s relationship reconstruction error for each number of subclones and number of cancer samples. **c.** Pairtree’s Pairs Tensor’s relationship reconstruction error for each number of subclones and number of cancer samples.

**Figure S6:**
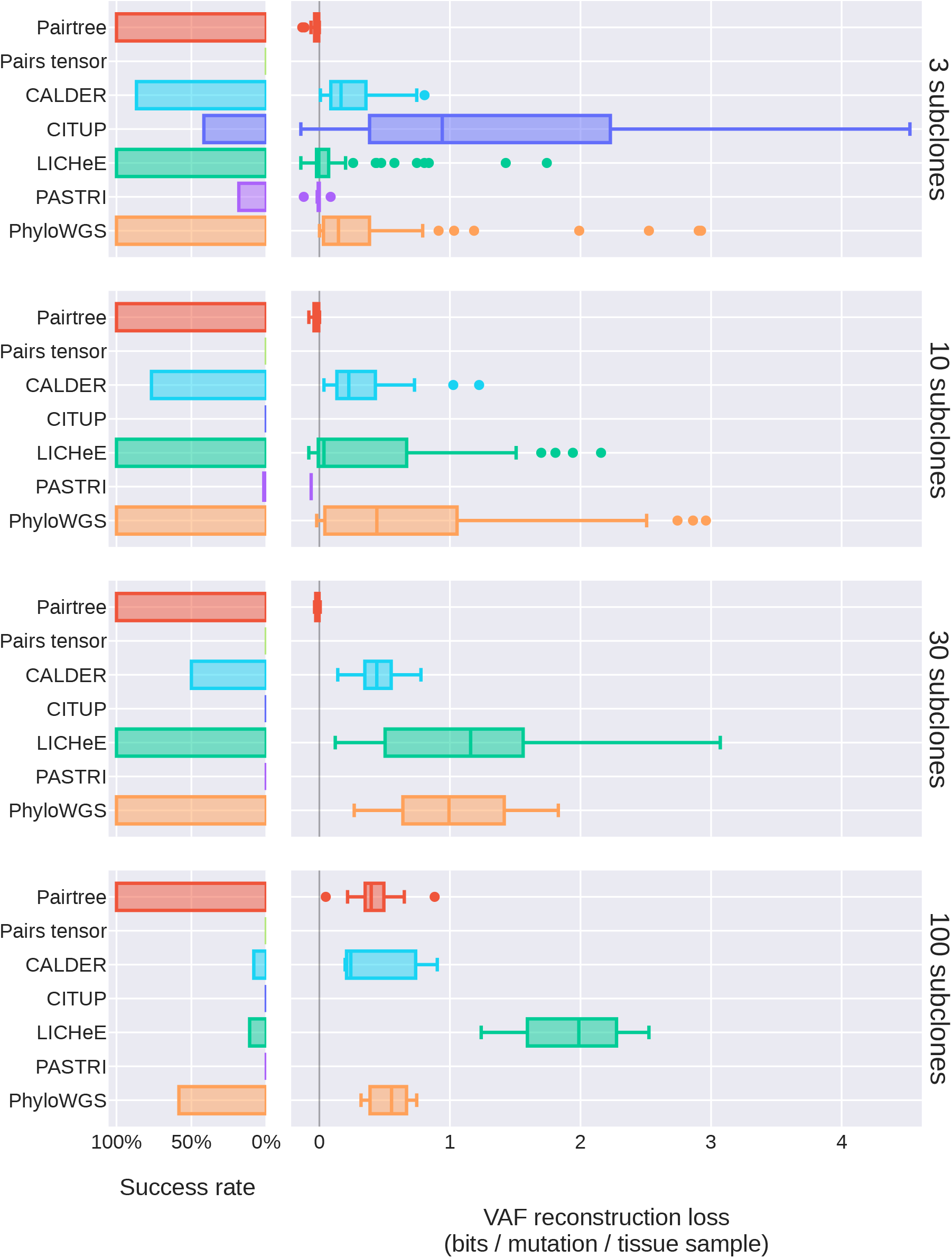
Pairtree’s VAF loss on 192 simulated low-depth datasets where the read depth was 50x.

**Figure S7:**
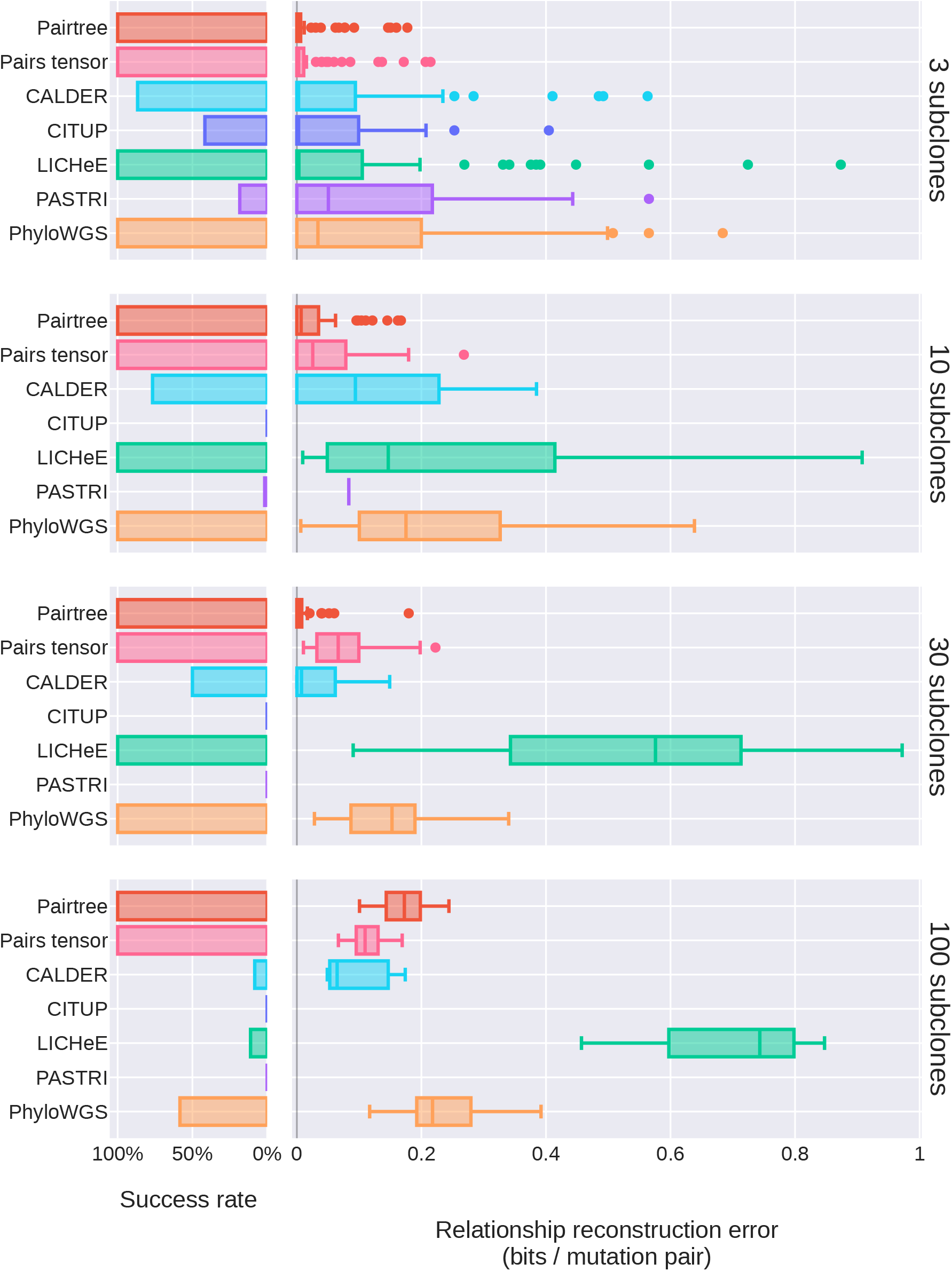
Pairtree’s relationship reconstruction error on 192 simulated low-depth datasets where the read depth was 50x.

**Figure S8:**
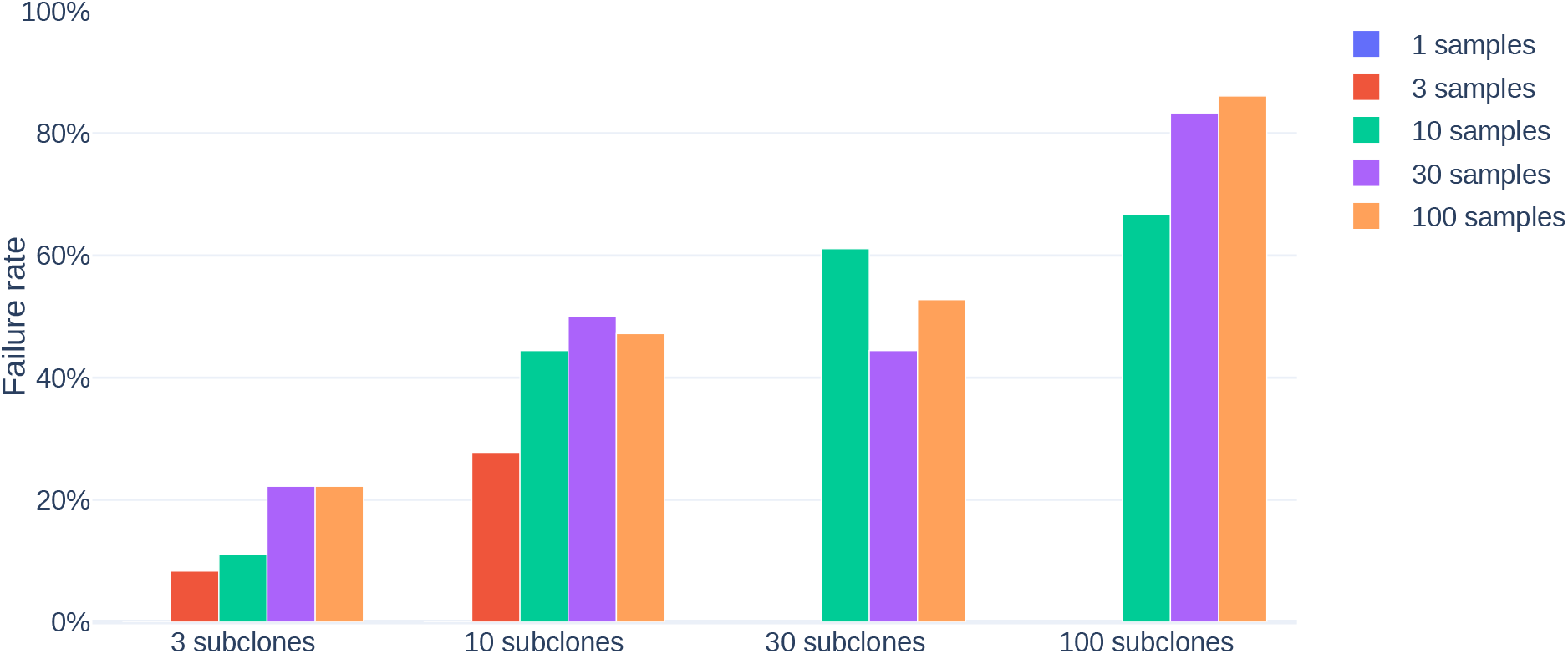
CALDER’s failure rate on 576 simulated datasets for different number of subclones and cancer samples.

**Figure S9:**
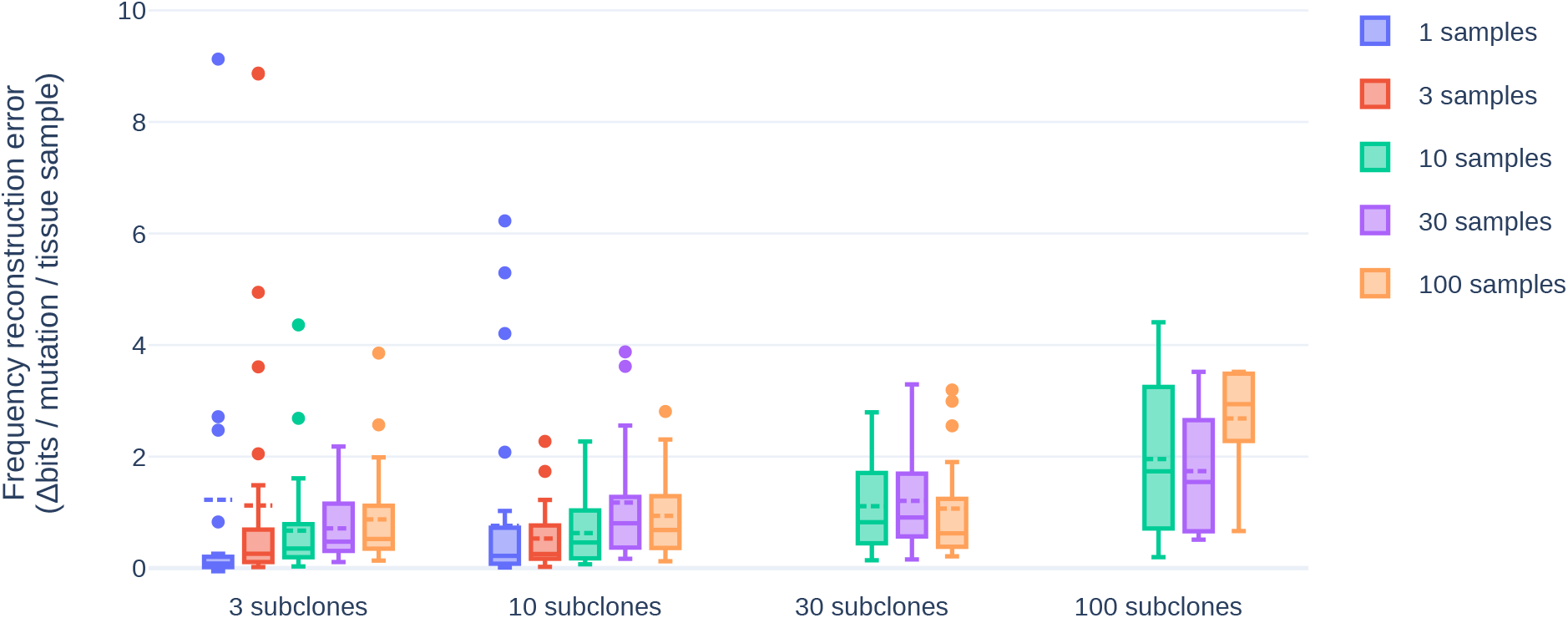
CALDER’s VAF loss on 576 simulated datasets for different number of subclones and cancer samples.

**Figure S10:**
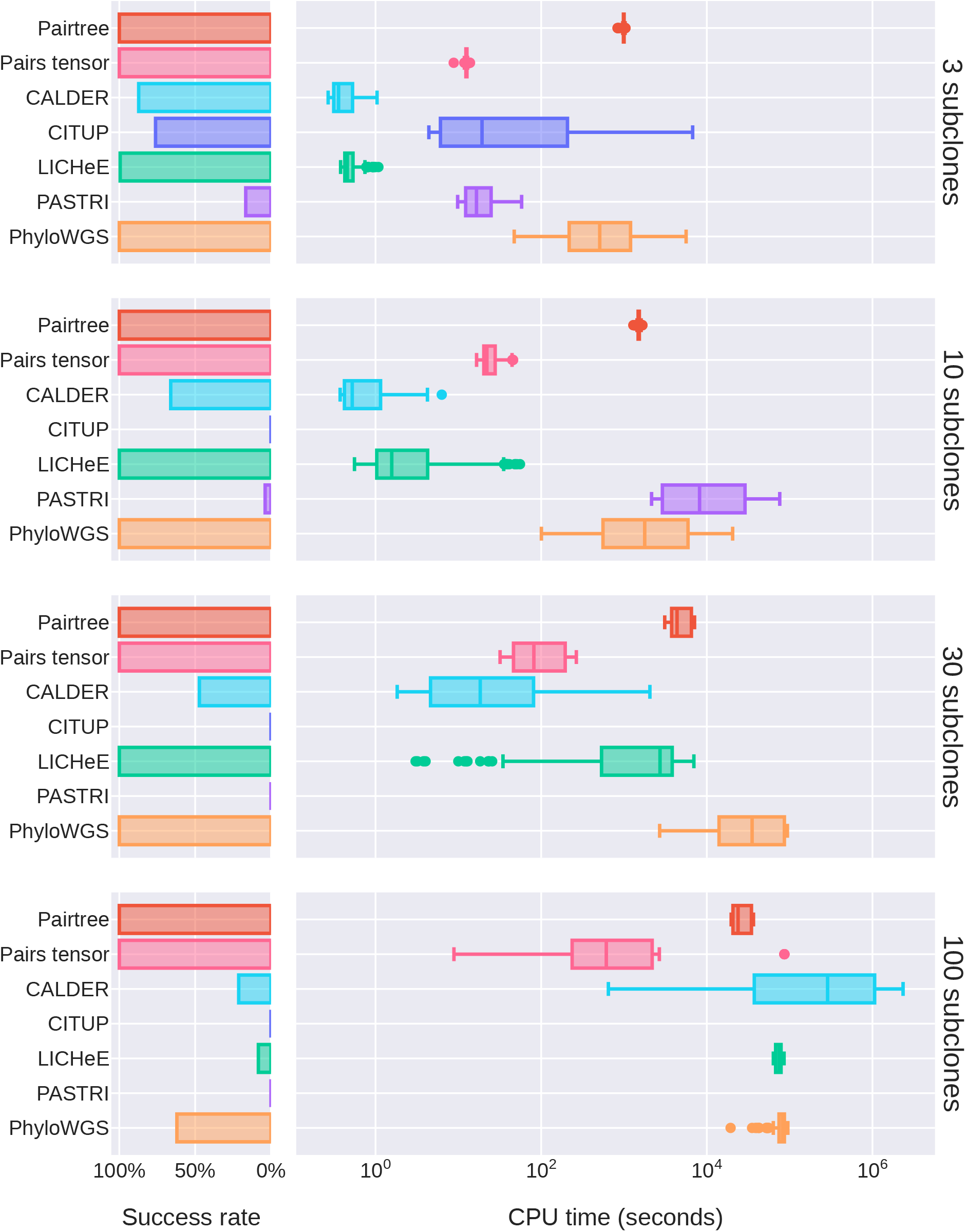
Number of CPU seconds methods took to produce results. Box mid-lines indicate medians. When using multiple CPU cores, these numbers can be much higher than elapsed wall-clock time (Fig. S11). Results for each method reflect only its performance on the datasets where it could produce a result.

**Figure S11:**
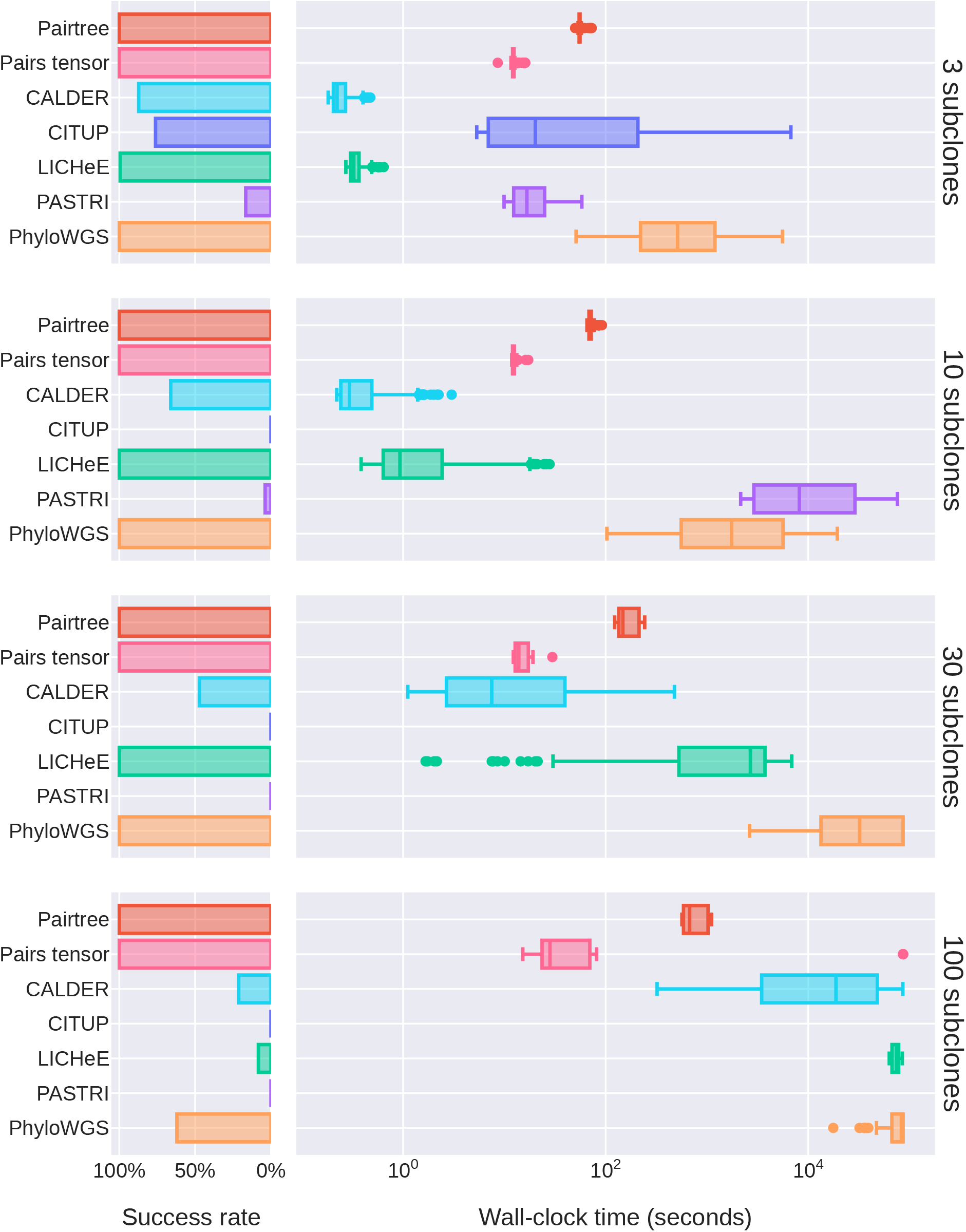
Elapsed wall-clock seconds methods took to produce results. Box mid-lines indicate medians. When using multiple CPU cores, these numbers can be much lower than the number of CPU seconds consumed (Fig. S10). Results for each method reflect only its performance on the datasets where it could produce a result.

**Figure S12:**
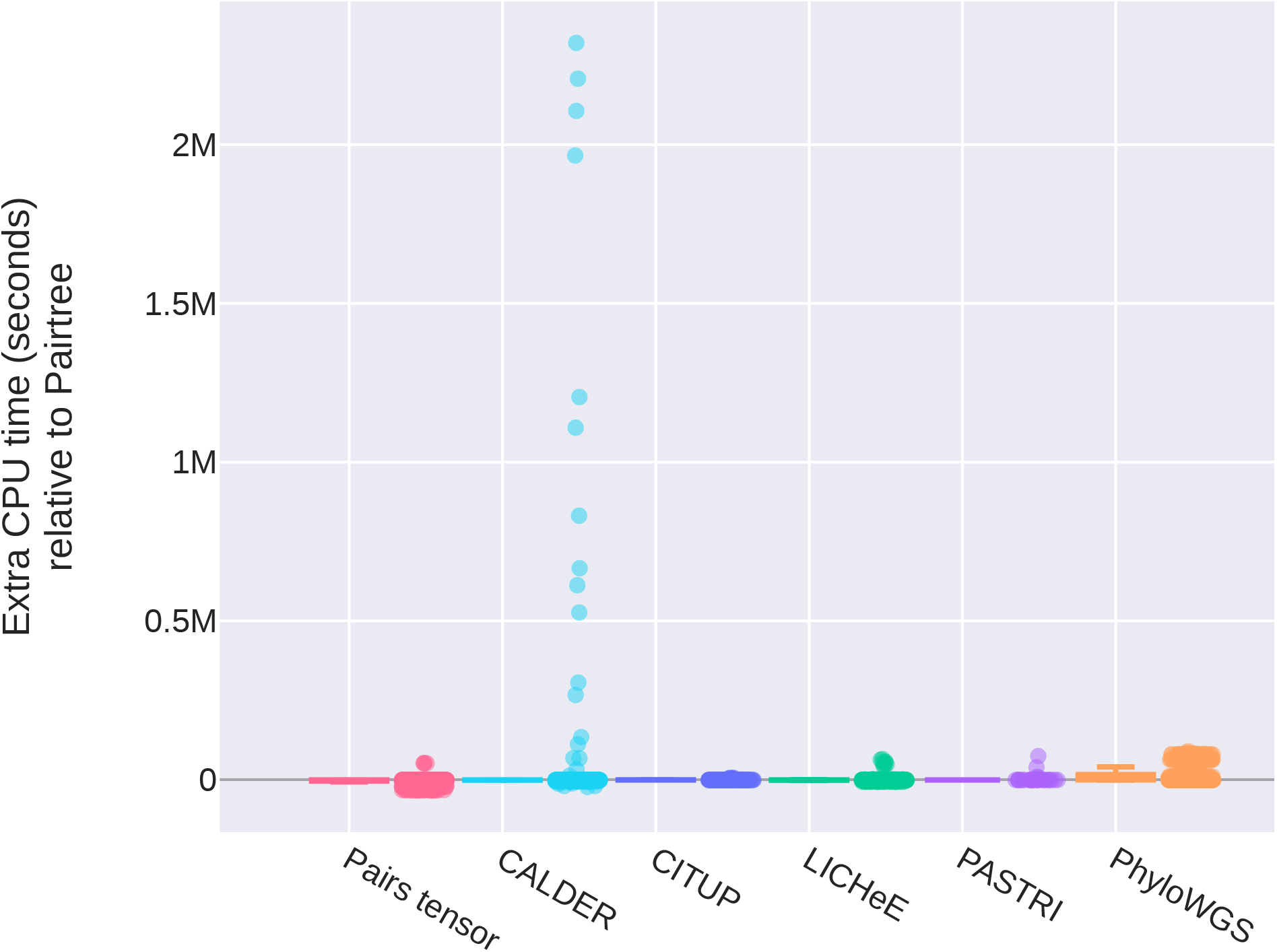
Number of CPU seconds each method took to produce results relative to Pairtree. Each point indicates the number of additional CPU seconds a method took on a dataset relative to Pairtree on that dataset. Points below zero indicate a method took less time than Pairtree on those datasets.

**Figure S13:**
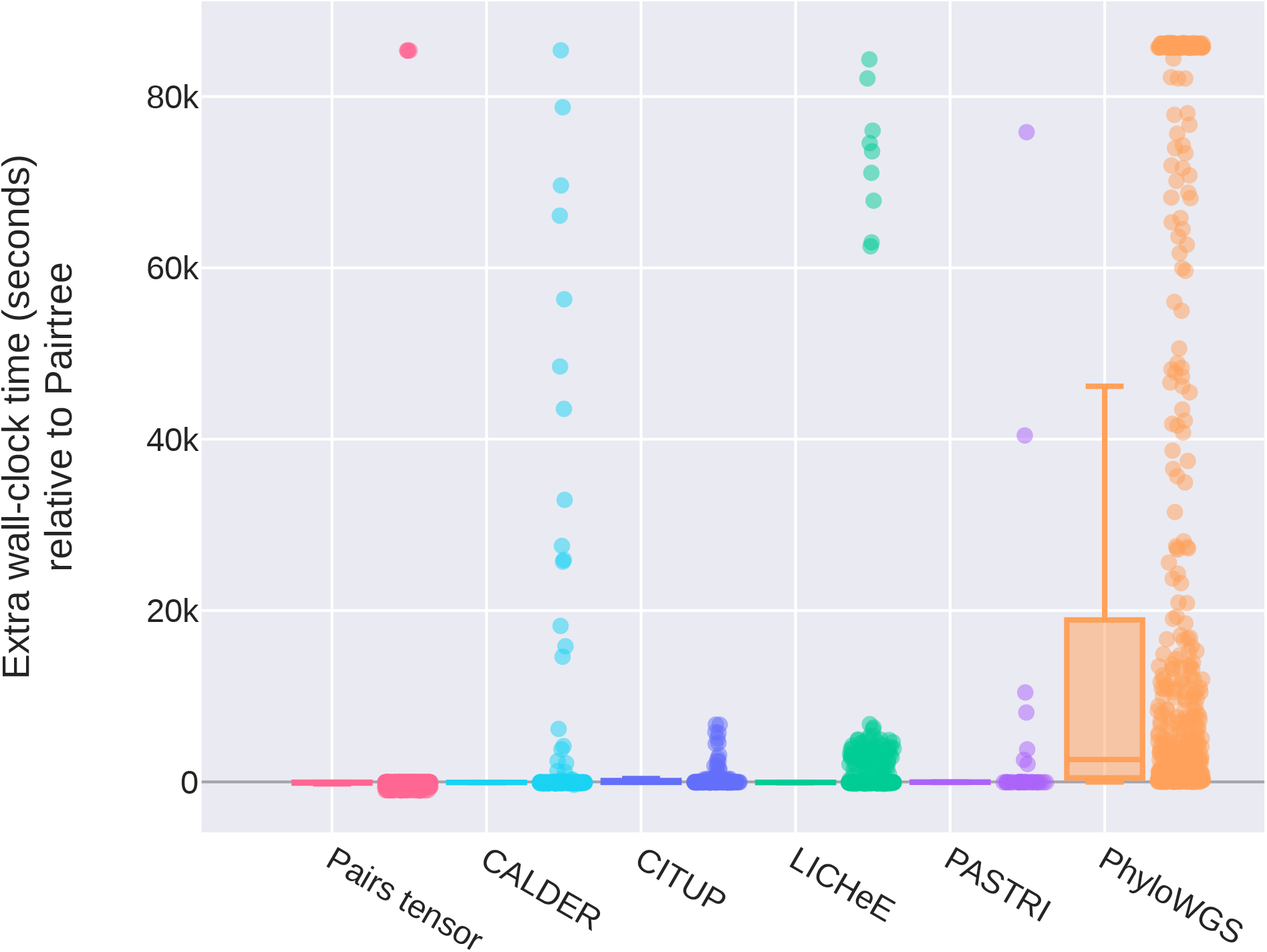
Elapsed wall-clock seconds each method took to produce results relative to Pairtree. Each point indicates the number of additional wall-clock seconds a method took on a dataset relative to Pairtree on that dataset. Points below zero indicate a method took less time than Pairtree on those datasets.

**Figure S14:**
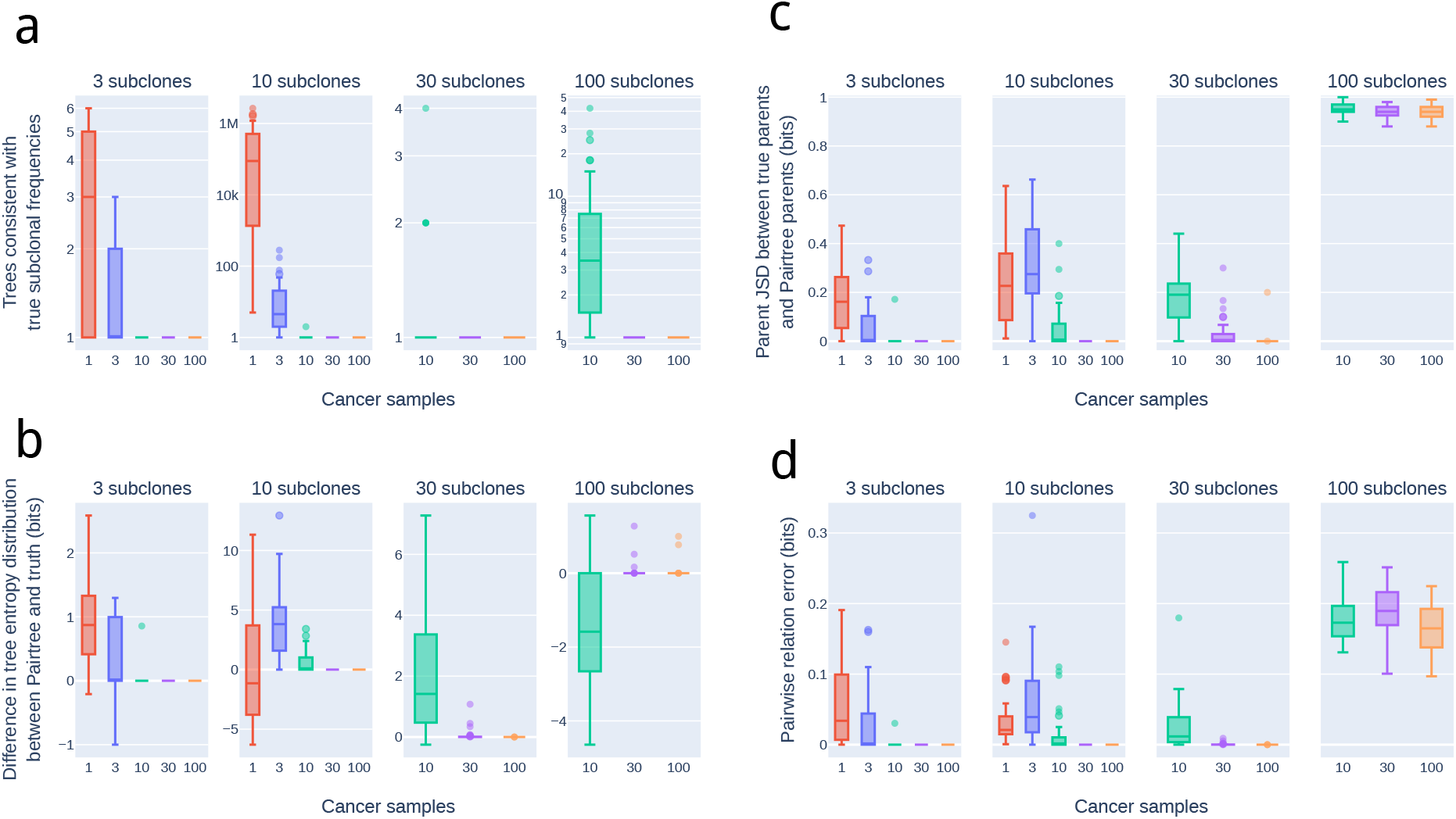
Characteristics of the distributions over possible trees for the 576 simulated clone tree reconstruction problems. Mid-lines in box plots indicate medians. **a.** Regardless of the number of subclones, with one cancer sample there are usually multiple trees consistent with the true subclonal frequencies. The highest median number of true trees (88,860) is reached for 10-subclone, single-sample reconstructions problems. Given ten or more samples, the tree becomes highly constrained, and there is usually only a single consistent tree. **b.** The entropies of the Pairtree-recovered tree distribution and true tree distribution reflect how many high-confidence trees Pairtree recovers relative to the number of possible trees. In general, Pairtree recognizes when the true tree is highly constrained, and returns only one high-confidence tree. **c.** For a simulated dataset, a distribution over possible trees induces a distribution over parent choice for every population represented in the tree. Shown are the joint Jensen-Shannon divergence between parent distributions for Pairtree relative to truth for each simulated dataset, normalized to the number of subclones in each tree. These divergences range between zero and one, with small values indicating that parent choices are nearly always correct. For a given number of subclones, Pairtree generally exhibits lower divergences with more cancer samples, indicating it was able to use the information provided by those samples to improve its solution set. **d.** Relationship reconstruction errors show that, even when the parents chosen for subclones are sometimes incorrect (panel c), the relationship reconstructions can be more accurate. This is the same information as presented in Fig. 3b, but partitioned by number of cancer samples.

**Figure S15:**
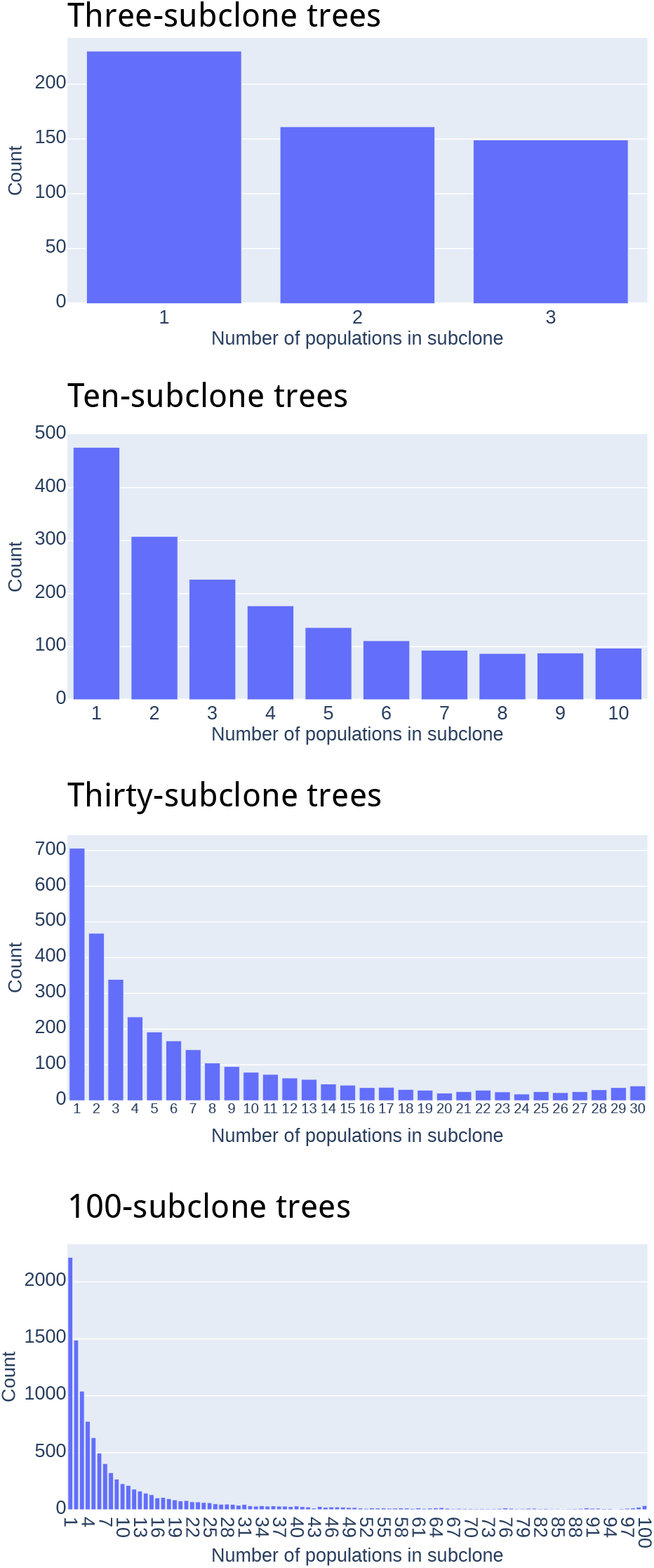
Prevalence of different subclone sizes within simulated trees. Subclone size indicates the number of subpopulations present within a subclone, reflecting the number of subpopulations that are descendants of the subpopulation that initiated the subclone.

**Figure S16:**
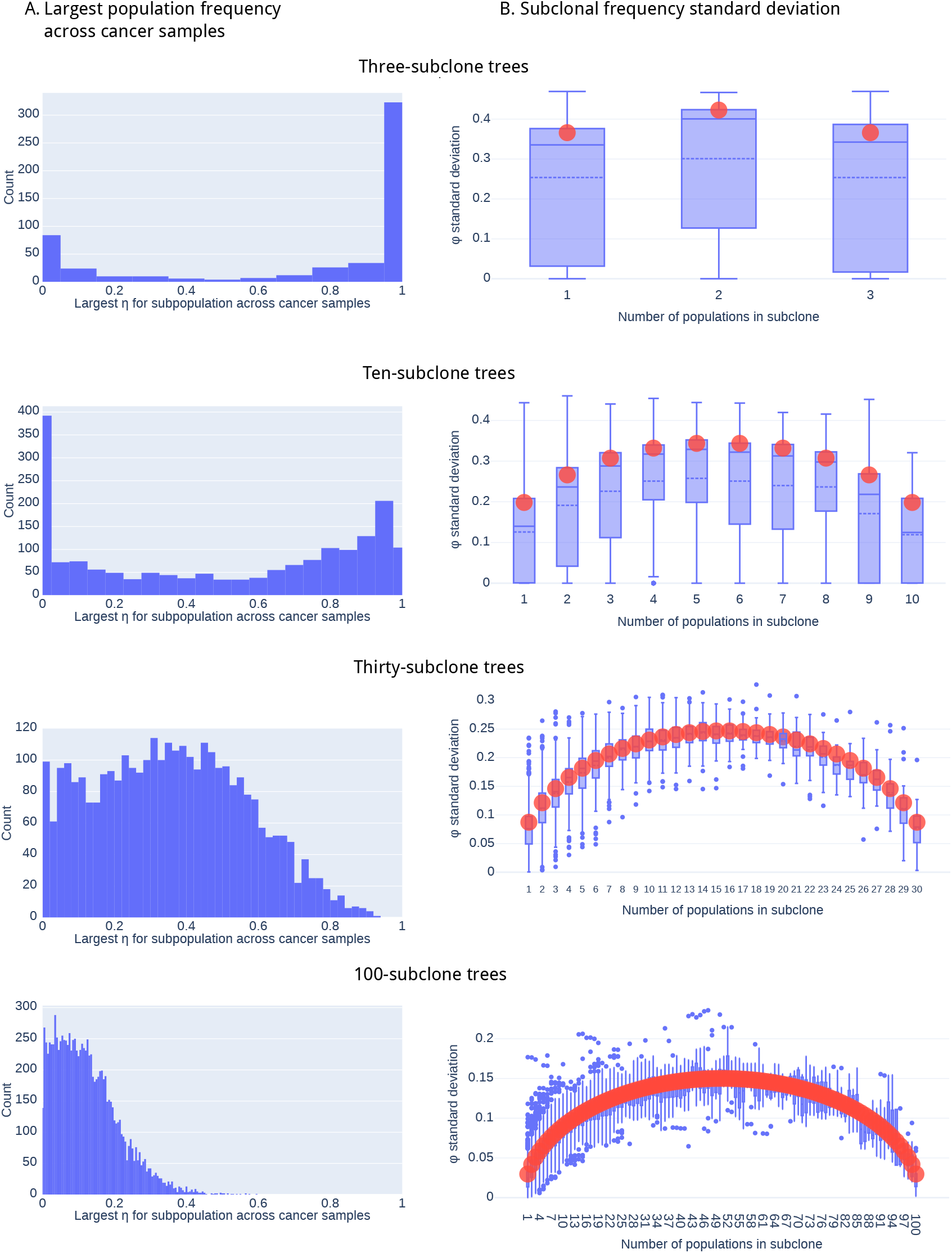
Properties of population and subclone frequencies. **a.** Largest population frequency *η_ks_* for each population *k* across cancer samples *s* in simulated data. **b.** Standard deviation of subclonal frequencies *ϕ_ks_* for each subclone *k* across cancer samples *s* in simulated data, as a function of the number of populations in the subclone. Box plots show the empirical standard deviation measured in (noise-free) simulated data, with solid line indicating the median and dashed line showing the mean. Orange circles show the predicted standard deviation derived from Dirichlet distribution properties.

**Figure S17:**
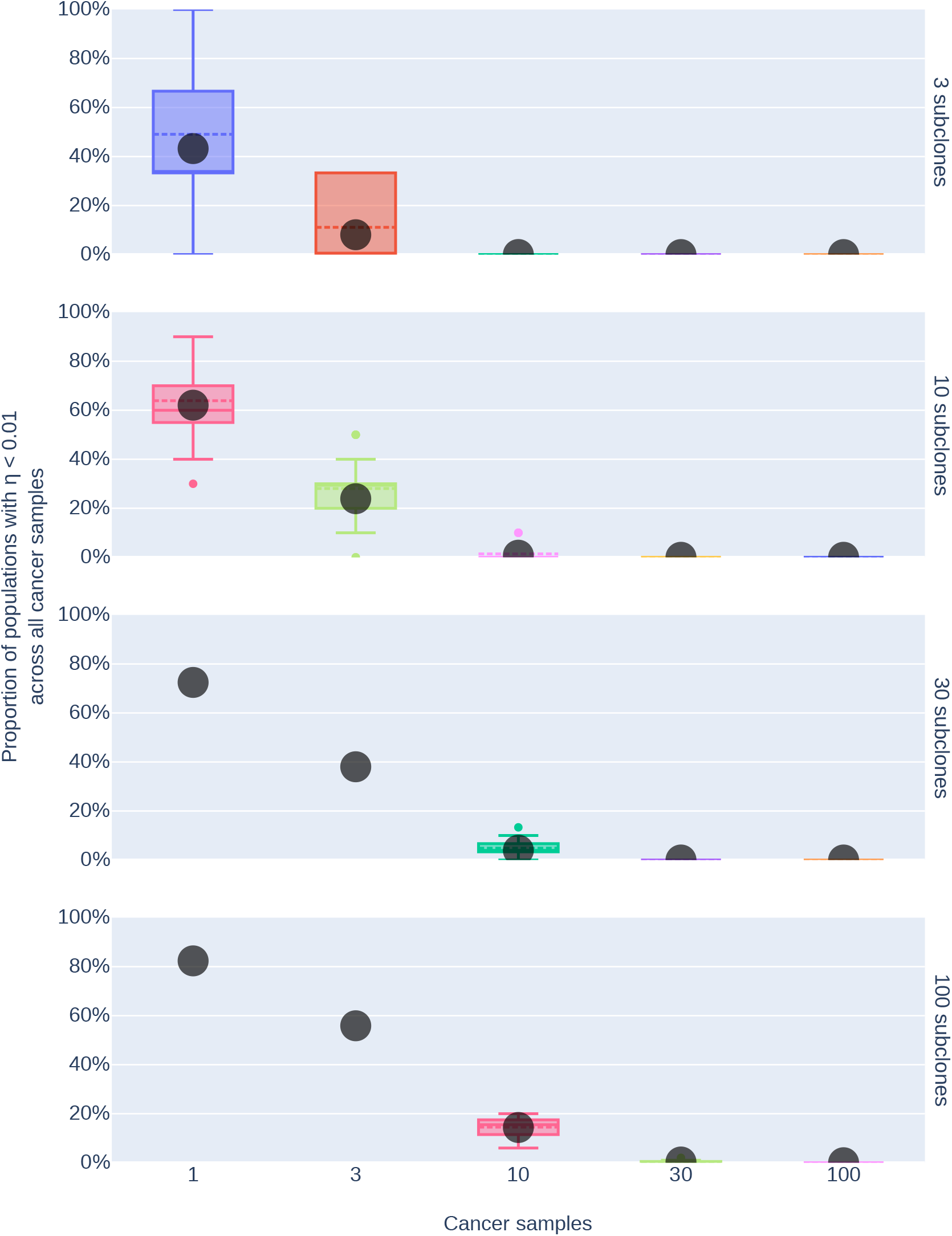
Proportion of populations with small population frequencies in all cancer samples. Proportion of populations *k* with population frequencies *η_ks_* < 1% across all cancer samples *s*. Box plots show the empirical proportions measured in (noise-free) simulated data, with solid line indicating the median and dashed line showing the mean. Grey circles show the predicted proportions derived from Dirichlet distribution properties.

**Figure S18:**
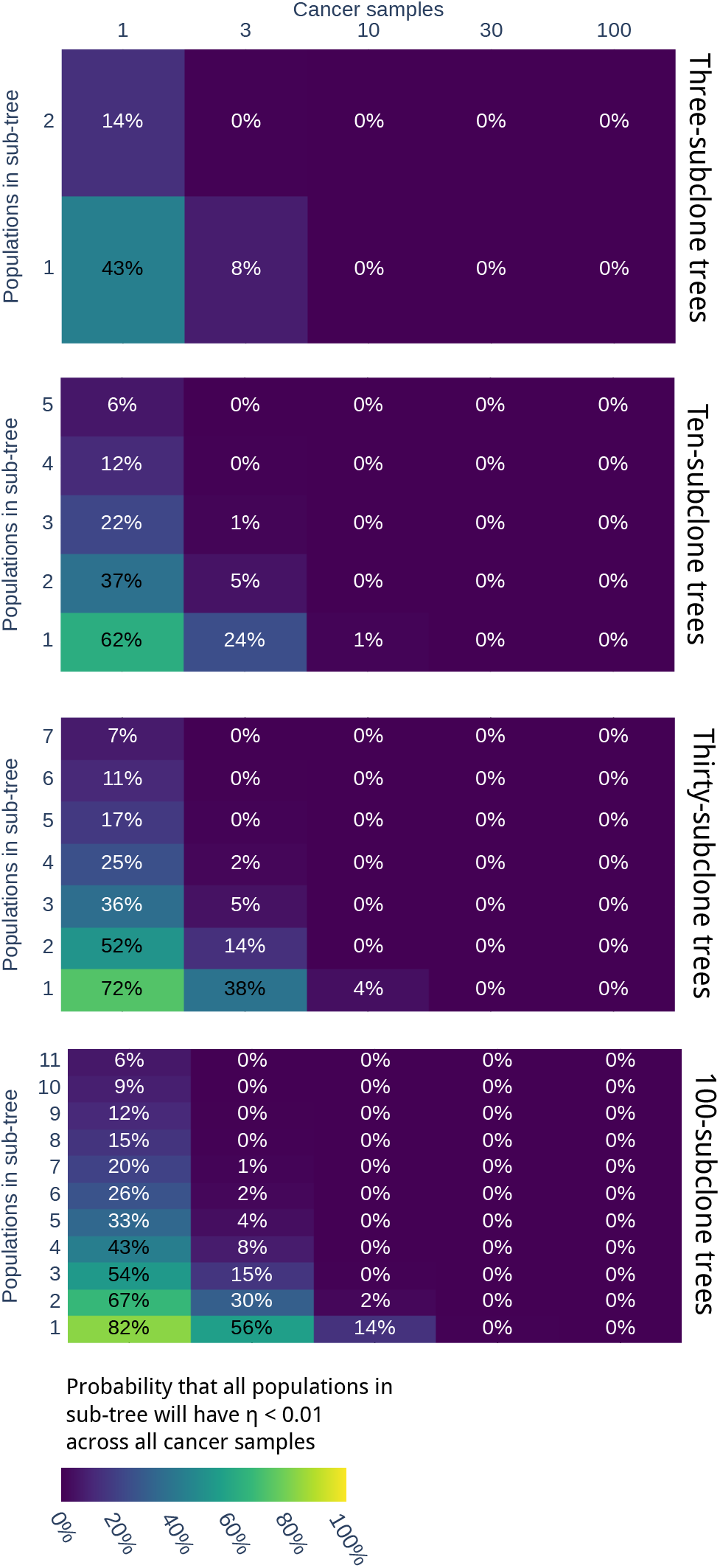
Probability that sub-trees will consist entirely of populations with small frequencies in all cancer samples. Probability that sub-tree containing given number of populations will have population frequencies *η_ks_* < 1% for all populations *k* in the sub-tree across all cancer samples *s*, computed using properties of Dirichlet distribution. A sub-tree consists of a subset of nodes from the full-tree and all edges between those nodes. By this definition, all subclones are sub-trees, but a sub-tree need not be a subclone.

**Figure S19:**
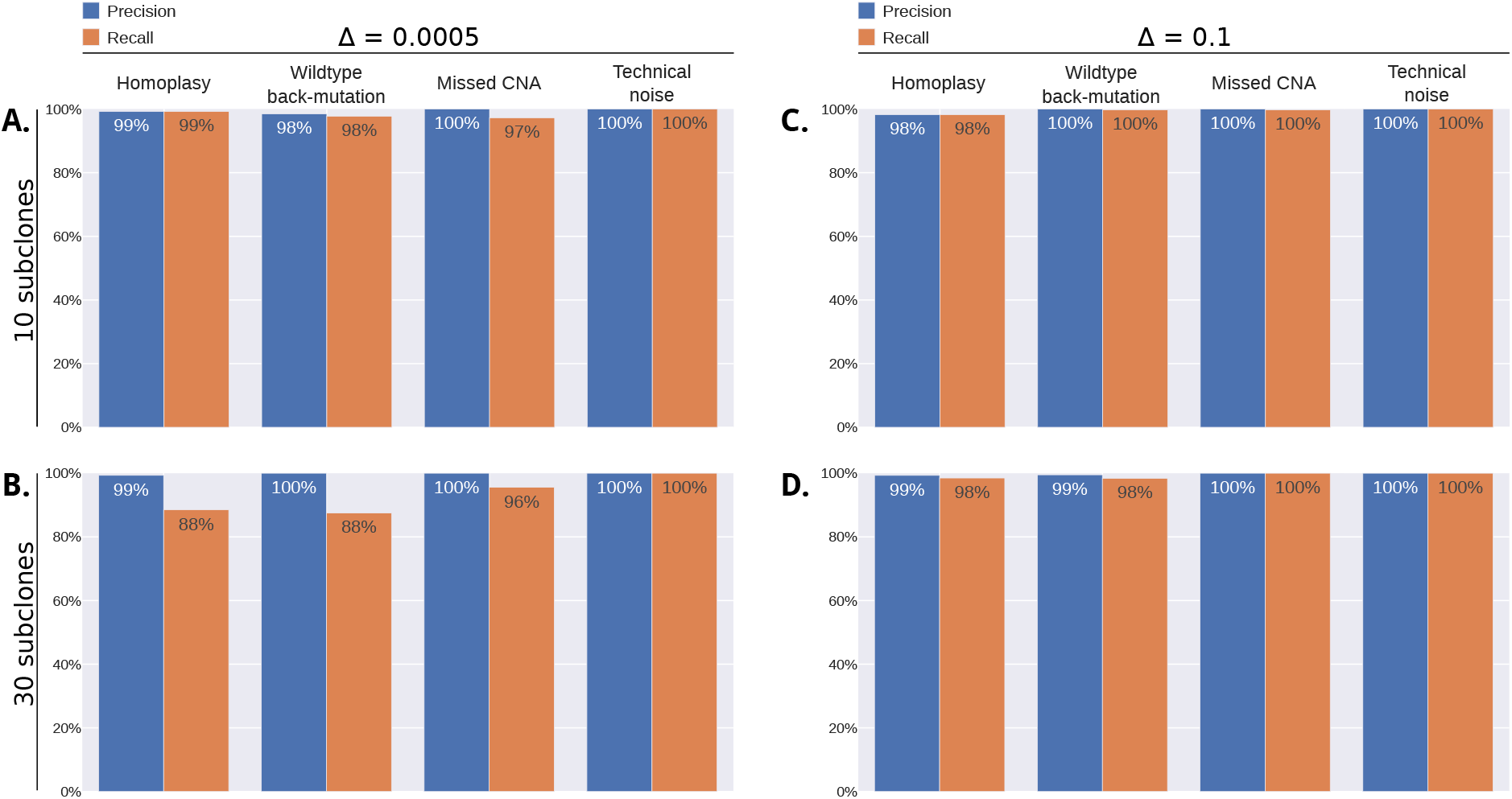
Ability to detect simulated garbage mutations using pairwise relations. Garbage mutations are mutations that do not fit in the tree because of ISA violations (homoplasy and wildtype back mutations), incorrect input ploidies (missed CNAs corresponding to LOH events), or technical noise. Pairtree can use pairwise relations to detect and discard such mutations. Shown are the precision and recall for this garbage detection procedure on 160 simulated trees with 10 subclones, 200 non-garbage mutations, and 20 garbage mutations per tree (**a.**), and on 30 subclones, 600 non-garbage mutations, and 60 garbage mutations per tree (**b.**). The number of true positives, false positives, and true negatives were combined across datasets in each class to produce single values for precision and recall.

**Figure S20:**
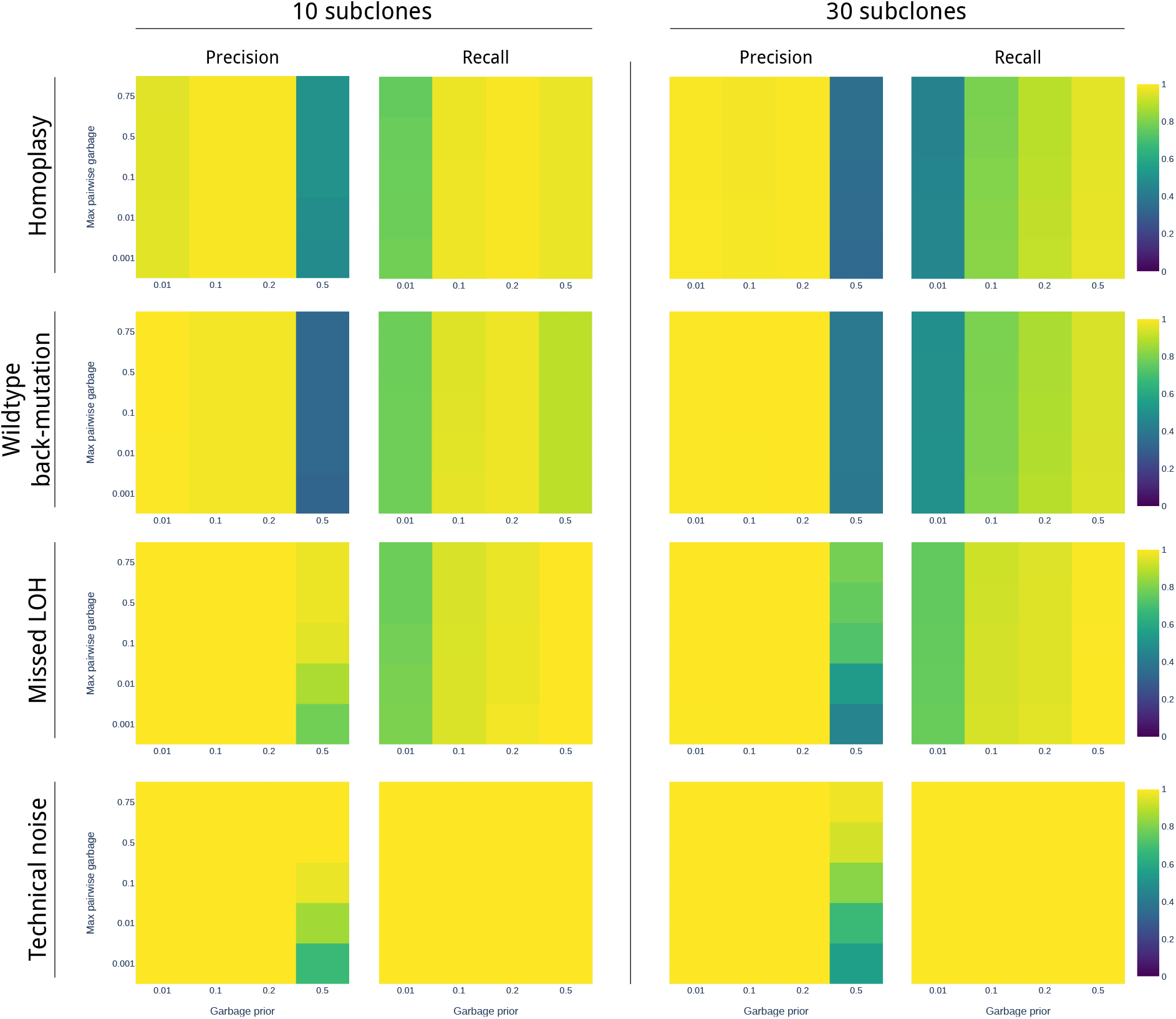
Influence of hyperparameters on garbage detection precision and recall. The hyperparameters represent the prior probability of the garbage relationship between mutation pairs (*γ*) and the maximum pairwise garbage probability permitted amongst mutations classified as not garbage (*ρ*). Performance is reported on the same 160 simulated clone trees used in Fig. S19.

**Figure S21:**
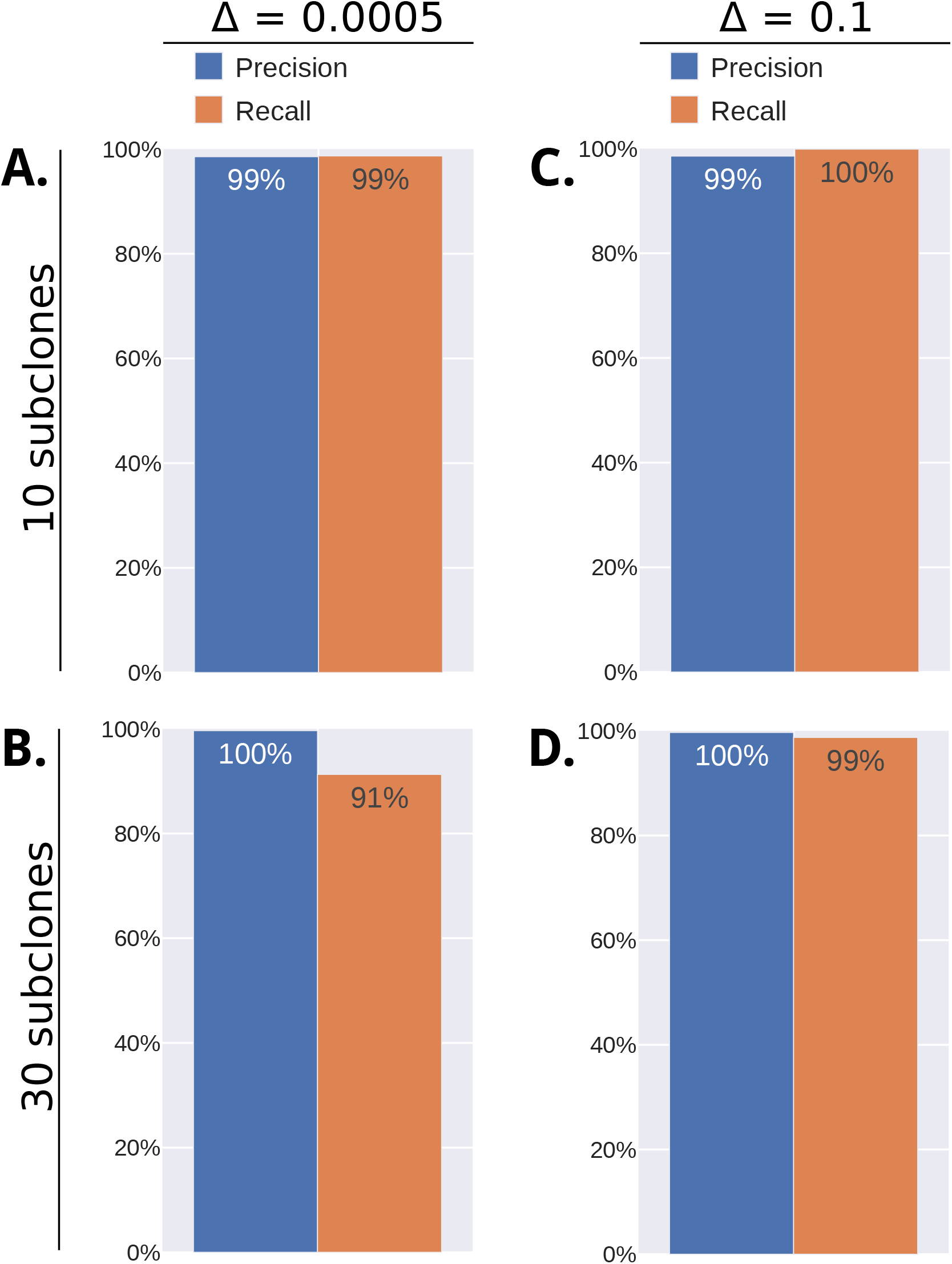
Ability to detect missed LOH events without using pairwise relations. If the input data given to Pairtree does not report CNA events corresponding to LOH, the mutation VAFs can imply subclonal frequencies outside a realistic range. Pairtree can detect such problems without having to use pairwise relationships to legitimate mutations. Shown are precision and recall for 100 trees with 10 subclones (200 legitimate mutations, 20 garbage mutations) and 100 trees with 30 subclones (600 legitimate mutations, 60 garbage mutations). The number of true positives, false positives, and true negatives were combined across datasets in each class to produce single values for precision and recall.

## References

1. Dentro, S. C. et al. Pervasive intra-tumour heterogeneity and subclonal selection across cancer types. Accepted at Cell (2021).

2. Hanahan, D. & Weinberg, R. a. Hallmarks of cancer: the next generation. Cell 144, 646–74. ISSN: 1097-4172. http://www.ncbi.nlm.nih.gov/pubmed/21376230 (Mar. 2011).

3. Gerstung, M. et al. The evolutionary history of 2,658 cancers. Nature 578, 122–128. ISSN: 1476-4687. https://www.nature.com/articles/s41586-019-1907-7 (2020)(Feb. 2020).

4. Espiritu, S. M. G. et al. The Evolutionary Landscape of Localized Prostate Cancers Drives Clinical Aggression. Cell 173, 1003–1013.e15. ISSN: 0092-8674. http://www.sciencedirect.com/science/article/pii/S009286741830309X (2018) (May 3, 2018).

5. Nik-Zainal, S. et al. The life history of 21 breast cancers. Cell 149, 994–1007 (2012).

6. Jamal-Hanjani, M. et al. Tracking the Evolution of Non–Small-Cell Lung Cancer. New England Journal of Medicine 376, 2109–2121. ISSN: 0028-4793, 1533-4406. http://www.nejm.org/doi/10.1056/NEJMoa1616288 (2018) (June 2017).

7. Gundem, G. et al. The evolutionary history of lethal metastatic prostate cancer. Nature 520, 353–357. ISSN: 1476-4687 (Apr. 16, 2015).

8. Sakamoto, H. et al. Evolutionary Origins of Recurrent Pancreatic Cancer. bioRxiv, 811133. https://www.biorxiv.org/content/10.1101/811133v1 (2019) (Oct. 31, 2019).

9. Alves, J. M., Prado-López, S., Cameselle-Teijeiro, J. M. & Posada, D. Rapid evolution and biogeographic spread in a colorectal cancer. Nature Communications 10. Number: 1 Publisher: Nature Publishing Group, 5139. ISSN: 2041-1723. https://www.nature.com/articles/s41467-019-12926-8 (2020) (Nov. 13, 2019).

10. Hu, Z. et al. Quantitative evidence for early metastatic seeding in colorectal cancer. Nature Genetics, 1. ISSN: 1546-1718. https://www.nature.com/articles/s41588-019-0423-x (2019) (June 17, 2019).

11. Dobson, S. M. et al. Relapse-Fated Latent Diagnosis Subclones in Acute B Lineage Leukemia Are Drug Tolerant and Possess Distinct Metabolic Programs. Cancer Discovery. Publisher: American Association for Cancer Research Section: Research Articles. ISSN: 2159-8274, 2159–8290. https://cancerdiscovery.aacrjournals.org/content/early/2020/03/12/2159-8290.CD-19-1059 (2020) (Feb. 21, 2020).

12. Hu, Z., Li, Z., Ma, Z. & Curtis, C. Multi-cancer analysis of clonality and the timing of systemic spread in paired primary tumors and metastases. Nature Genetics 52. Number: 7 Publisher: Nature Publishing Group, 701–708. ISSN: 1546-1718. https://www.nature.com/articles/s41588-020-0628-z (2020) (July 2020).

13. Zahir, N., Sun, R., Gallahan, D., Gatenby, R. A. & Curtis, C. Characterizing the ecological and evolutionary dynamics of cancer. Nature Genetics 52. Number: 8 Publisher: Nature Publishing Group, 759–767. ISSN: 1546-1718. https://www.nature.com/articles/s41588-020-0668-4 (2020) (Aug. 2020).

14. Williams, M. J. et al. Quantification of subclonal selection in cancer from bulk sequencing data. Nature Genetics, 1. ISSN: 1546-1718. http://www.nature.com/articles/s41588-018-0128-6 (2018) (May 28, 2018).

15. Pogrebniak, K. L. & Curtis, C. Harnessing Tumor Evolution to Circumvent Resistance. Trends in Genetics 34, 639–651. ISSN: 01689525. https://linkinghub.elsevier.com/retrieve/pii/S0168952518300921 (2018) (Aug. 2018).

16. Coorens, T. H. H. et al. Inherent mosaicism and extensive mutation of human placentas. Nature 592. Number: 7852 Publisher: Nature Publishing Group, 80–85. ISSN: 1476-4687. https://www.nature.com/articles/s41586-021-03345-1 (2021) (Apr. 2021).

17. Bizzotto, S. et al. Landmarks of human embryonic development inscribed in somatic mutations. Science 371. Publisher: American Association for the Advancement of Science Section: Report, 1249–1253. ISSN: 0036-8075, 1095-9203. https://science.sciencemag.org/content/371/6535/1249 (2021) (Mar. 19, 2021).

18. Jiang, Y., Qiu, Y., Minn, A. J. & Zhang, N. R. Assessing intratumor heterogeneity and tracking longitudinal and spatial clonal evolutionary history by next-generation sequencing. Proceedings of the National Academy of Sciences 113, E5528–E5537. ISSN: 0027-8424, 1091-6490. http://www.pnas.org/content/113/37/E5528 (2017) (Sept. 13, 2016).

19. Malikic, S., McPherson, A. W., Donmez, N. & Sahinalp, C. S. Clonality inference in multiple tumor samples using phylogeny. Bioinformatics 31, 1349–1356. ISSN: 1460-2059, 1367-4803. https://academic.oup.com/bioinformatics/article-lookup/doi/10.1093/bioinformatics/btv003 (2019) (May 1, 2015).

20. Malikic, S., Jahn, K., Kuipers, J., Sahinalp, S. C. & Beerenwinkel, N. Integrative inference of subclonal tumour evolution from single-cell and bulk sequencing data. Nature Communications 10. Number: 1 Publisher: Nature Publishing Group, 2750. ISSN: 2041-1723. https://www.nature.com/articles/s41467-019-10737-5 (2020) (June 21, 2019).

21. Deshwar, A. G., Vembu, S. & Morris, Q. Comparing nonparametric Bayesian tree priors for clonal reconstruction of tumors. Pacific Symposium on Biocomputing. Pacific Symposium on Biocomputing, 20–31. ISSN: 2335-6936 (2015).

22. Popic, V. et al. Fast and scalable inference of multi-sample cancer lineages. Genome Biology 16. ISSN: 1465-6906. https://www.ncbi.nlm.nih.gov/pmc/articles/PMC4501097/ (2017) (2015).

23. Satas, G. & Raphael, B. J. Tumor phylogeny inference using tree-constrained importance sampling. Bioinformatics 33, i152–i160. ISSN: 1367-4803. https://academic.oup.com/bioinformatics/article/33/14/i152/3953987/Tumor-phylogeny-inference-using-tree-constrained (2017) (July 15, 2017).

24. Salcedo, A. et al. A community effort to create standards for evaluating tumor subclonal reconstruction. Nature Biotechnology 38. Number: 1 Publisher: Nature Publishing Group, 97–107. ISSN: 1546-1696. https://www.nature.com/articles/s41587-019-0364-z (2020) (Jan. 2020).

25. Dentro, S. C., Wedge, D. C. & Van Loo, P. Principles of Reconstructing the Subclonal Architecture of Cancers. Cold Spring Harbor Perspectives in Medicine, a026625 (2017).

26. Kuipers, J., Jahn, K., Raphael, B. J. & Beerenwinkel, N. Single-cell sequencing data reveal widespread recurrence and loss of mutational hits in the life histories of tumors. Genome Research 27, 1885–1894. ISSN: 1088-9051. https://www.ncbi.nlm.nih.gov/pmc/articles/PMC5668945/ (2020) (Nov. 2017).

27. Singer, J., Kuipers, J., Jahn, K. & Beerenwinkel, N. Single-cell mutation identification via phylogenetic inference. Nature Communications 9, 5144. ISSN: 2041-1723. https://www.nature.com/articles/s41467-018-07627-7 (2020) (Dec. 4, 2018).

28. McPherson, A. et al. Divergent modes of clonal spread and intraperitoneal mixing in high-grade serous ovarian cancer. Nature Genetics 48, 758–767. ISSN: 1546-1718 (2016).

29. Bonizzoni, P., Ciccolella, S., Della Vedova, G. & Soto, M. Does relaxing the infinite sites assumption give better tumor phylogenies? An ILP-based comparative approach. bioRxiv. http://biorxiv.org/lookup/doi/10.1101/227801 (2019) (Dec. 3, 2017).

30. Ciccolella, S. et al. Inferring Cancer Progression from Single-cell Sequencing while Allowing Mutation Losses. bioRxiv. http://biorxiv.org/lookup/doi/10.1101/268243 (2019) (Apr. 13, 2018).

31. Satas, G., Zaccaria, S., Mon, G. & Raphael, B. J. SCARLET: Single-Cell Tumor Phylogeny Inference with Copy-Number Constrained Mutation Losses. Cell Systems 10, 323–332.e8. ISSN: 2405-4712. http://www.sciencedirect.com/science/article/pii/S2405471220301150 (2020) (Apr. 22, 2020).

32. Roth, A. et al. PyClone: statistical inference of clonal population structure in cancer. Nature methods (2014).

33. Miller, C. A. et al. SciClone: Inferring Clonal Architecture and Tracking the Spatial and Temporal Patterns of Tumor Evolution. PLoS Computational Biology 10 (ed Beerenwinkel, N.) e1003665. ISSN: 1553-734X. JSTOR: {PMC}4125065. http://www.ncbi.nlm.nih.gov/pmc/articles/PMC4125065/ (Aug. 2014).

34. Gillis, S. & Roth, A. PyClone-VI: Scalable inference of clonal population structures using whole genome data. bioRxiv. Publisher: Cold Spring Harbor Laboratory Section: New Results, 2020.08.31.276212. https://www.biorxiv.org/content/10.1101/2020.08.31.276212v1 (2020) (Sept. 1, 2020).

35. Satas, G. & Raphael, B. PASTRI source code Aug. 16, 2019. https://github.com/raphael-group/PASTRI (2020).

36. Tarabichi, M. et al. A practical guide to cancer subclonal reconstruction from DNA sequencing. Nature Methods 18. Number: 2 Publisher: Nature Publishing Group, 144–155. ISSN: 1548-7105. https://www.nature.com/articles/s41592-020-01013-2 (2021)(Feb. 2021).

37. Hastings, W. K. Monte Carlo Sampling Methods Using Markov Chains and Their Applications. Biometrika 57. Publisher: [Oxford University Press, Biometrika Trust], 97–109. ISSN: 0006-3444. https://www.jstor.org/stable/2334940 (2020) (1970).

38. Jahn, K., Kuipers, J. & Beerenwinkel, N. Tree inference for single-cell data. Genome Biology 17, 86. ISSN: 1474-760X. http://dx.doi.org/10.1186/s13059-016-0936-x (2016) (2016).

39. Jiao, W., Vembu, S., Deshwar, A. G., Stein, L. & Morris, Q. Inferring clonal evolution of tumors from single nucleotide somatic mutations. BMC bioinformatics 15, 35. ISSN: 1471-2105. http://www.pubmedcentral.nih.gov/articlerender.fcgi?artid=3922638&tool=pmcentrez&rendertype=abstract (Jan. 2014).

40. Deshwar, A. G. et al. PhyloWGS: Reconstructing subclonal composition and evolution from whole genome sequencing of tumors. Genome Biology 16, 35 (2015).

41. Jia, B., Ray, S., Safavi, S. & Bento, J. Efficient Projection onto the Perfect Phylogeny Model. arXiv:1811.01129 [cs]. arXiv: 1811.01129. http://arxiv.org/abs/1811.01129 (2018) (Nov. 2, 2018).

42. Sundermann, L. K., Wintersinger, J., Rätsch, G., Stoye, J. & Morris, Q. Reconstructing tumor evolutionary histories and clone trees in polynomial-time with SubMARine. PLOS Computational Biology 17. Publisher: Public Library of Science, 1–28. https://doi.org/10.1371/journal.pcbi.1008400 (2021).

43. Myers, M. A., Satas, G. & Raphael, B. J. CALDER: Inferring Phylogenetic Trees from Longitudinal Tumor Samples. Cell Systems 8. Publisher: Elsevier, 514–522.e5. ISSN: 2405-4712. https://www.cell.com/cell-systems/abstract/S2405-4712(19)30191-7 (2021) (June 26, 2019).

44. Strino, F., Parisi, F., Micsinai, M. & Kluger, Y. TrAp: a tree approach for fingerprinting subclonal tumor composition. Nucleic Acids Research 41, e165–e165. ISSN: 1362-4962, 0305-1048. https://academic.oup.com/nar/article/41/17/e165/2411666 (2021) (Sept. 1, 2013).

45. Hudson, R. R. & Kaplan, N. L. Statistical Properties of the Number of Recombination Events in the History of a Sample of DNA Sequences. Genetics 111, 147–164. ISSN: 0016-6731. https://www.ncbi.nlm.nih.gov/pmc/articles/PMC1202594/ (2019) (Sept. 1985).

46. Riedmiller, M. & Braun, H. A direct adaptive method for faster backpropagation learning: The RPROP algorithm in IEEE international conference on neural networks (1993), 586–591.

47. Hinton, G. Neutral Networks for Machine Learning: Lecture 6a: Overview of min-batch gradient descent. 2014. https://www.cs.toronto.edu/~tijmen/csc321/slides/lecture_slides_lec6.pdf.

48. Bei, J. & Bento, J. libprojectppm source code original-date: 2018-10-26T22:41:45Z. Apr. 8, 2019. https://github.com/bentoayr/Efficient-Projection-onto-the-Perfect-Phylogeny-Model.

49. Gusfield, D. Efficient algorithms for inferring evolutionary trees. Networks 21, 19–28. ISSN: 10970037. http://onlinelibrary.wiley.com/doi/10.1002/net.3230210104/abstract (2017) (Jan. 1, 1991).

50. El-Kebir, M., Oesper, L., Acheson-Field, H. & Raphael, B. J. Reconstruction of clonal trees and tumor composition from multi-sample sequencing data. Bioinformatics (Oxford, England) 31, i62–70. ISSN: 1367-4811 (June 15, 2015).

51. Stewart, C. A. et al. Single-cell analyses reveal increased intratumoral heterogeneity after the onset of therapy resistance in small-cell lung cancer. Nature Cancer 1. Number: 4 Publisher: Nature Publishing Group, 423–436. ISSN: 2662-1347. https://www.nature.com/articles/s43018-019-0020-z (2020) (Apr. 2020).

52. Miles, L. A. et al. Single cell mutational profiling delineates clonal trajectories in myeloid malignancies. bioRxiv. Publisher: Cold Spring Harbor Laboratory Section: New Results, 2020.02.07.938860. https://www.biorxiv.org/content/10.1101/2020.02.07.938860v1 (2020) (Feb. 9, 2020).

53. Lähnemann, D. et al. Eleven grand challenges in single-cell data science. Genome Biology 21, 31. ISSN: 1474-760X. https://doi.org/10.1186/s13059-020-1926-6 (2020) (Feb. 7, 2020).

54. Sun, R. et al. Between-region genetic divergence reflects the mode and tempo of tumor evolution. Nature Genetics 49. Bandiera_abtest: a Cg_type: Nature Research Journals Number: 7 Primary_atype: Research Publisher: Nature Publishing Group Subject_term: Cancer;Computational biology and bioinformatics;Genomics Subject_term_id: cancer;computational-biology-and-bioinformatics;genomics, 1015–1024. ISSN: 1546-1718. https://www.nature.com/articles/ng.3891 (2021) (July 2017).

55. Williams, M. J., Werner, B., Barnes, C. P., Graham, T. A. & Sottoriva, A. Identification of neutral tumor evolution across cancer types. Nature Genetics 48, 238–244. ISSN: 1546-1718 (Mar. 2016).

56. Caravagna, G. et al. Detecting repeated cancer evolution from multi-region tumor sequencing data. Nature Methods 15, 707–714. ISSN: 1548-7091, 1548-7105. http://www.nature.com/articles/s41592-018-0108-x (2021) (Sept. 2018).

57. Caravagna, G. et al. Subclonal reconstruction of tumors by using machine learning and population genetics. Nature Genetics 52, 898–907. ISSN: 1546-1718 (2020).

